# Forecasting trade and biosecurity risk under climate change

**DOI:** 10.1101/2024.06.30.601437

**Authors:** James Camac, Matthew Cantele, Van Ha Pham, Christine Li, Andrew Robinson, Tom Kompas

**Affiliations:** The Centre of Excellence for Biosecurity Risk Analysis, The University of Melbourne; Crawford School of Public Policy, Australian National University

## Abstract

1.

It is well known that the increasing globalisation of human movement and trade has led to substantial range expansions of many taxa, and as a consequence, is increasingly exposing countries to novel biosecurity threats. This increased exposure to novel threats is expected to be exacerbated by a changing climate as trade patterns and species distributions change. In many countries biosecurity regulators are already struggling to contend with the increased exposure to pests and diseases, and as such, must shift towards a pro-active strategy of anticipating risk. However, to anticipate risk, regulators must make better use of the data they collect (e.g. border interceptions) and develop models capable of forecasting biosecurity propagule pressure and establishment exposure as a function of changing trade, human population, and species distributions.

In this project, we develop a highly innovative model framework that simulates annual changes in international trade patterns (imports & exports) as a function of climate change impacts on crop productivity, labour productivity (via heat stress) and the amount of arable land. These simulated changes in global trade are then coupled with border interception data, and annual predicted changes in the geographic distributions of both human populations and threat climate suitability to estimate climate-induced changes in: 1) contamination rates for imports from different trading partners, 2) the total amount of biosecurity risk material arriving at Australia’s borders, and 3) its exposure to an establishment event (Figure 1.1).

Figure 1.1.:

Conceptual diagram of innovative biosecurity forecasting model. The model is made up of two primary sub-modules: 1) The Economic submodel, which incorporates climate change damages (e.g. crop productivity & heat stress) and simulates these impacts on global import and export trade patterns; 2) The Ecological submodel that estimates threat climate suitability, country-by-commodity contamination rates, and then links these to predicted changes in both global trade flows (i.e. outputs from the Economic submodel) and human population to approximate country-specific propagule pressure and establishment exposure.

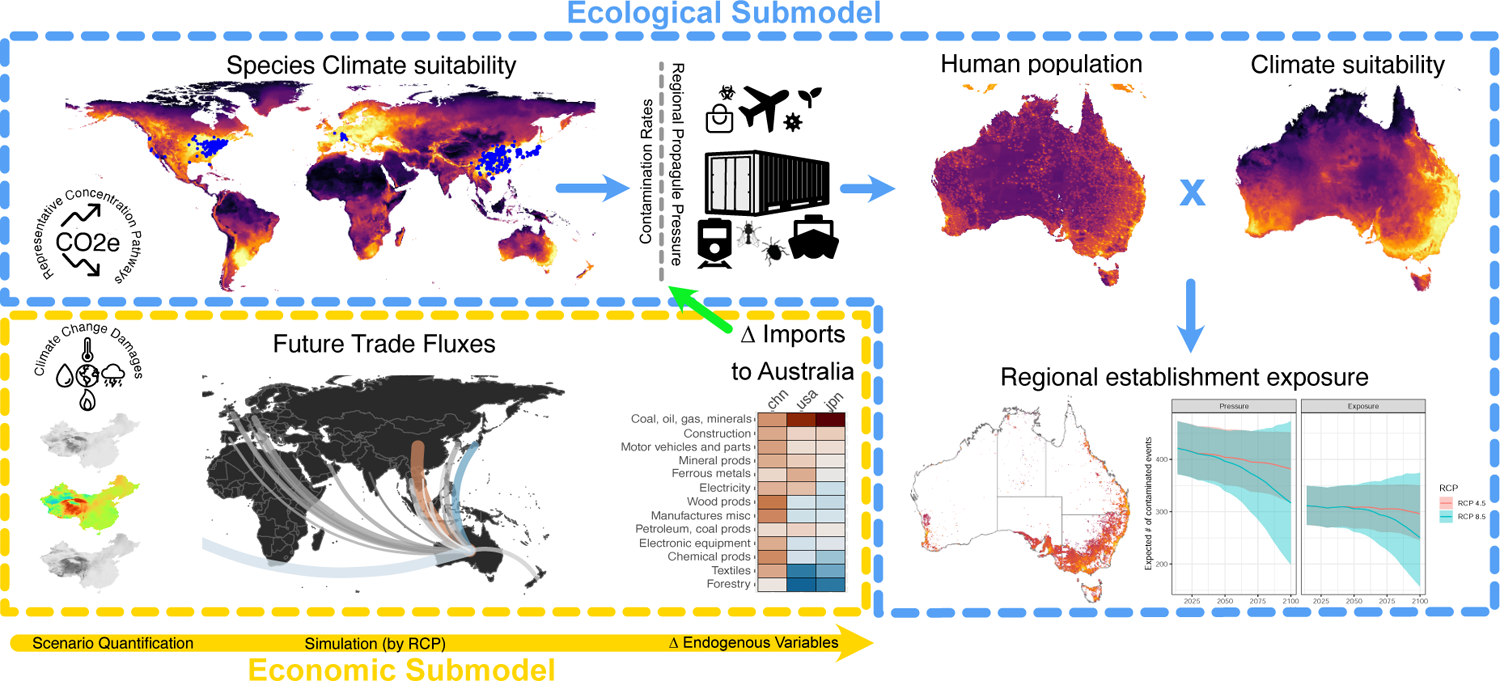

Here, we provide an in-depth description of both the trade and biosecurity risk sub-models. We then illustrate the model’s implementation and outputs under two different climate scenarios (RCP 4.5 and RCP 8.5) for the purposes of forecasting temporal changes in: 1) global and regional economic trade; and 2) the amount of biosecurity risk material arriving at Australia’s border (i.e. propagule pressure) and what it may mean for establishment exposure. In illustrating the biosecurity risk, we focused on five plant pests deemed by the Department of Agriculture, Fisheries and Forestry (DAFF) as an exemplar of different hitch-hiking functional groups. The exemplar threats examined were:

- **Overwintering** – brown marmorated stink bug (BMSB; *Halyomorpha halys*)
- **Egg laying** – Spongy moth (*Lymantria dispar*)
- **Nesting** – Asian honey bee (*Apis cerana*)
- **Sheltering** – Giant African snail (*Lissachatina fulica*)
- **Internal storage** – Khapra beetle (*Trogoderma granarium*).

**Key findings & model utility:** *Global and regional economic trade consequences of climate change:* The climate-enhanced trade model shows significant changes in trade patterns globally. Climate change impacts various countries differently, causing substantial changes in national incomes, productive capacity, and imports and exports around the world. This occurs even with relatively limited damage functions from global warming. Indeed, with limited damage functions (crop productivity losses, labour productivity losses due to heat stress, and loss of arable land from ocean inundation), the impact on national income in Australia from climate change is relatively small (e.g., damages from more substantive sea level rise and storm surge are not included). However, damages to trading partners and changes in imports to Australia, as well as the rest of the world, are considerable. To allow for easy interrogation of these predicted trade impacts, we have developed an interactive dashboard^1^ that allows users to examine changes in trade patterns – by country, commodity and climate change scenario – for user selected imports by country and total values. The following summarise the key findings from the select damage functions employed within this analysis:

- Western African regions fare the worst in terms of overall GDP impacts, with the region at large experiencing roughly a 20% loss to GDP. The second most affected region is South Asia^2^ with Pakistan bearing the greatest damages. Some countries emerged relatively unscathed, prominently those located in North America, Europe, Russia, and Oceania.
- In terms of climate change related damages to key Australian trading partners, we find that losses in China and Southeast Asia^3^ are the most consequential. Here labour productivity shocks due to heat stress lead to losses in manufacturing sector imports such as *Motor vehicles* and *Electronic equipment*. As the largest exporter of manufacturing goods and a key region affected by climate change, China sees the largest decline in exports, however most of South and Southeast Asia fares worse in terms of relative losses with India and Indonesia seeing over 10% losses. Conversely, net gains are made across several European regions, particularly Germany and the UK.
- The greatest heterogeneity within macroeconomic sector was observed among agricultural imports and in particular crop sectors in large part due to the varied mix of exporters for each crop type and differential impacts experienced. The largest absolute increase among food- and agriculture-related sectors is seen within *Vegetables, fruit, nuts* with large increases in exports from OECD regions and China.
- The climate change damage functions employed here likely vastly underestimate the true costs of climate change impacts for a number of reasons. These include the following:
  1. Recent scholarship suggests that large damages to GDP are already being experienced (Romanello *et al*., 2023) whereas the earlier work used here (with the exception of crop productivity) includes minimal damages at early timesteps. Further, the anticipated impacts from warming may be far larger (particularly at later timesteps) if climate sensitivities have been misestimated (leading to more extreme associated temperature paths) or tipping points are triggered.
  2. The application of damages used within the current study is limited in terms of scope, with damages due to human health impacts and changes in tourism trends missing. In addition to the limited scope, some damages that have been implemented have not been fully allocated to all affected sectors. For example, sea level rise is understood entirely within the purview of loss of arable land while physical capital depreciation and impacts to manufacturing hubs located on coastal areas is not currently simulated.
  3. Sociopolitical consequences of climate change such as armed conflict and forced migration are not included. Diminished social resilience due to climate change related impacts (e.g., rising food insecurity) has the potential to feed back into the economy in myriad ways, potentially cascading into additional economic damages and/or altered trade patterns.
- Although limited climate change damages have been implemented, large changes in socioeconomic trends including economic and demographic growth are anticipated to occur within the long-term time horizon used within the current study. Such changes affect key Australian trade partners and have the potential to alter the composition of imports at a magnitude equal to or surpassing climate change impacts. Full simulation of both socioeconomic pathways as well as climate change impacts is currently a CEBRA priority.

*Changes in pest propagule pressure & establishment exposure hitting Australia.:* While this report focuses on long-term trends in annual propagule pressure and establishment exposure hitting Australia’s border, the model is capable of estimating such trends for all 70 global regions and can be readily updated with new interception data. As such, to facilitate interrogation of model outputs, we have developed leaflet interactive maps for each of the five threats. Included in these maps are information^4^ about temporal trends in pest pressure, establishment exposure (if threat is influenced by climate suitability) and proportion of risk attributable to each infected exporters and commodity sector. The following summarise the key findings we found by running the model for the five exemplar threats:

- Out of the five exemplar threats, brown marmorated stink bug (BMSB; *Halyomorpha halys*) posed the greatest propagule pressure and established risk to Australia. This was followed by giant African snail (*Lissachatina fulica*), Khapra beetle (*Trogoderma granarium*), Asian honey bee (*Apis cerana*), and then spongy moth (*Lymantria dispar*).
- Imports of manufacturing commodities (e.g. electronic equipment, motor vehicles and parts, plastics and metals) from infected countries posed the greatest risk of hitch-hiker contamination, with some exporting regions posing greater risk than others.
- We detected marginal (albeit uncertain) positive impacts of potential area of suitable climate on the contamination rates coming from infected exporting regions for BMSB and to a lesser extent Spongy moth. This meant that in exporting regions where climate suitability is predicted to increase (e.g. Canada & United Kingdom for BMSB), contamination rates will be higher. By contrast, in regions where climate suitability is expected to decrease (e.g. Italy for BMSB), contamination rates are expected to decline. Area of suitable climate had no discernable impacts on contamination rates associated with Asian honey bee or giant African snail.^5^
- Differences in trends in predicted propagule pressure (i.e. number of contaminated lines) hitting Australia’s border between the two climate scenarios were found to be small and highly uncertain for all threats examined in this report. Multiple reasons are attributable to this:
  1. There are compensatory effects occurring, whereby reductions (due to either decreased trade or climate-induced declines in contamination likelihoods) in contaminated items from one infected exporter is made up by increases in another.
  2. Climate impacts on exports and imports of manufactured goods – the commodity types with highest likelihoods of containing most of the modelled hitch-hikers used in this study – are likely to be severely underestimated. Recent work (Romanello *et al*., 2023) suggests that the existing set of damages included may severely underestimate the true impacts of climate change, especially on manufacturing sectors. Extending the trade model to include a larger set of damage functions, particularly those that encompass physical capital depreciation due to sea level rise and other disasters magnified by climate change (e.g., bushfires), is expected to result in significantly greater economic impacts within affected manufacturing and related sectors. This in turn will manifest in more significant impacts on global export and import flows, and thus, the total pest pressure expected at Australia’s border.
  3. There remains significant uncertainty on how climate suitability in exporting regions impacts contamination rates. While our model detected marginal positive relationships between contamination rates and area of suitable climate for some threats (e.g. BMSB), uncertainty is partly driven by the number of infected exporting countries containing the threat and how variable climate suitability is within and among those trading partners. This uncertainty could be further reduced by including additional interception data from other countries (e.g. New Zealand) as well as incorporating additional information on other factors likely to impact contamination likelihoods such as known extent of established populations, time since establishment, or other factors that can inform population size in exporting country.
- Trends in Australia’s exposure to potential BMSB or Spongy moth establishment were expected to decline overtime due to a combination of reduced propagule pressure (caused by declines in volume of trade from high risk trading partners and/or climate-induced declines contamination rates), and declines in climatic suitability at Australian locations where goods are likely to be unpacked. These declines were most pronounced under the worst-case climate scenario (RCP 8.5). By contrast, no discernable trends were found for Asian honey bee or giant African snail – mostly due to 1) minor impacts on trade volumes from infected exporters; 2) negligible impacts of climate suitability on contamination rates; and 3) climate suitability in locations import goods are most likely to be unpacked remaining relatively stable (i.e. where urban centres are predicted to remain).

*Potential model utility:* We believe that this new model provides the necessary information needed for governments and industry to better anticipate – and therefore manage and adapt to – future climate change impacts on domestic and international economies, global trade, and exposure to biosecurity risk. The current work may serve as a foundation for informing policy decisions as well as a range of downstream models. For example model outputs can be used:

1. to directly inform both short-term and long-term policies associated with border screening effort among different commodities and exporters, threat prioritisation and developing or updating Import Risk Assessments (IRAs).
2. as inputs into other models used by DAFF such as: the post-border pest establishment likelihood model edmaps (Camac *et al*., 2020, 2021b)^6^ for informing post-border surveillance & establishment potential, the Risk Return Resource Allocation (RRRA) model, and CEBRA’s value model (Dodd *et al*., 2021) for informing potential damages accrued over time by an exotic threat and the benefits of different mitigation strategies.
3. to identify regions and economic sectors most susceptible to changing climate, and thus, provide governments with the necessary foresight to plan and adapt to potential impacts.

## 3. Background

The globalisation of human movement and trade has led to substantial range expansions of many taxa, and as a consequence, has increasingly exposed countries to novel biosecurity threats (Hulme, 2009; Seebens *et al*., 2017, 2021). Changing climate is expected to have both antagonistic and synergistic impacts on this exposure. It can directly alter geographic distributions of suitable habitat required by species, with some ranges expanding while others contract. Climate change is also expected to significantly impact humans – the primary disperser of biosecurity risk – by changing where people live and the activities they can undertake (i.e. what can they farm/produce). This raises a critical question for pest risk analysts: *How can one estimate and forecast biosecurity risk under changing climate and trade conditions*? The answer to which will ultimately guide how one may mitigate threat entry, establishment and spread risk using a variety of pre-border (e.g. trade restrictions), border (e.g. screening risk profiles), and post-border (e.g. surveillance) interventions.

Integrated trade, climate and pest risk modelling can be used to shed light on this question. Camac *et al*. (2021a) recently implemented such an integrated model for the Australian Department of Agriculture, Fisheries and Forestry. This global, innovative, and pragmatic model estimates both the expected number of threat-specific contamination events arriving at a country’s border (i.e., propagule pressure), and the expected number of these events likely to be unloaded in climatically suitable locations within a country (i.e., establishment exposure). It achieved this by integrating border interceptions with data on: 1) past international trade flows, 2) known pest occurrence, and 3) the geographic distributions of both human population and threat climate suitability. This model was originally applied to Brown Marmorated Stink Bug (BMSB) – a significant horticultural threat that has rapidly spread from its native range in East Asia (China, Japan, the Korean Peninsula and Taiwan) into Europe, North America and Canada, and in doing so, has caused significant agricultural losses (Rice *et al*., 2014; Valentin *et al*., 2017). While the original model was successful at making short-term predictions of country propagule pressure and establishment exposure of hitch-hiking threats, its predictions were predicated on the basis that recent (1-5 year) past trade flows, geographic distributions of human population and threat climate suitability did not change. As such, predictions were short-term focused (i.e., 1 year in advance), where changes in these factors were expected to be small.

Within the current project we build upon the above approach by simulating the impact of temperature change according to two long-term climate change scenarios within a large-dimensional, temporally dynamic constrained optimisation model known as a computable general equilibrium (CGE) model^1^. Within the current CGE model developed by Kompas *et al*. (2018), an equilibrium state representing all major economic expenditures as well as trade flows according to the Global Trade Analysis Project (GTAP) database is shocked to incorporate select impacts from climate change. This is accomplished through the use of select damage functions applied to technological variables within the CGE model, effectively approximating changes in productivity (and relative comparative advantage) for 70 regions and 40 commodity sectors across the entire globe. The magnitude, origin, and destination of trade flows adjust through the time horizon in response to altered productivity levels.

This integration of spatio-temporal predictions in trade, threat climate suitability and human population allows the model to predict how propagule pressure and establishment exposure for different hitch-hiking threats may vary over time under a range of temperature trajectories. Moreover, it allows users to examine expected changes in global temporal patterns in imports and exports, which vary tremendously from country to country. The combination of which can be used to anticipate future pathways of biosecurity risk, and thus, inform and prioritise how pre-border, border and post-border risk mitigation interventions should be utilised. This framework allows for long-term forecast of region-specific propagule pressure and establishment exposure of hitch-hiking threats under different climate change scenarios. We believe that this model, and the outputs it can deliver, will improve both Pest Risk Assessments (PRAs) and Import Risk Assessments (IRAs) undertaken by regulators. More importantly, as the model utilises hard-won data collected by governments and is capable of forecasting risk, we believe it will aid regulators to move from reactionary risk management towards proactive risk management in terms of both biosecurity threats and potential economic risks.

## 4. Modelling regional impacts of climate change on primary production and trade

The current modelling approach builds upon the standard GTAP model (Hertel, 1997) and previous work modeling the impacts of climate change Kompas *et al*. (2018); Kom-pas & Ha (2021). Within the current application, a computationally appropriate model is developed with a focus on major trading partners for Australia and elaborated climate change damages. Here an inter-temporal version of the GTAP model Ha *et al*. (2017) is used to simulate climate change impacts on the world economy. The Global Trade Analysis Project (GTAP) model is an inter-regional economic model where countries/regions of the world can produce, consume, and trade goods and services. The model is implemented through the solving of a large systems of equations which dictate the relationships between variables and parameters of interest.

The original GTAP model is a static model where the initial capital stock stays constant within a single period of time. The solution to a static model optimises over this single period, however, static models can be solved recursively to simulate a dynamic scenario. The end of period capital stock can be accumulated by investment minus depreciation. Investment goods are produced by combining goods and services (investment demand) and add to the capital stock. Investment is funded by household saving and borrowing (or lending) money from all over world (see Hertel, 1997). This type of dynamic model is often considered “myopic” because shocks in future time steps are not considered within the solution of the current time step (Babiker *et al*., 2009). Indeed, price expectations for the next period (only) are based solely on current and a weighted average of past prices.

The model used within the current analysis, however, is a GTAP variant which uses inter-temporal dynamics, solving the long run inter-temporal profit (dividend) maximisation problem for producers (Ha *et al*., 2017). In this regard, Kompas *et al*. (2018) has shown the importance of long run dynamic economic models in evaluating climate change impacts, which vary considerably from region to region.^1^ Our version of the dynamic CGE model is not subject to the same “myopic” forecasting rules and associated criticisms surrounding recursive models.

In particular, the component solution for the forward-looking producer problem that is different from the traditional recursive GTAP model can be summarised in terms of a system of motion equations:

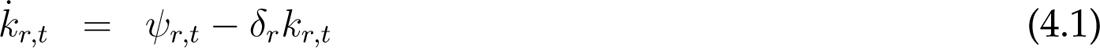

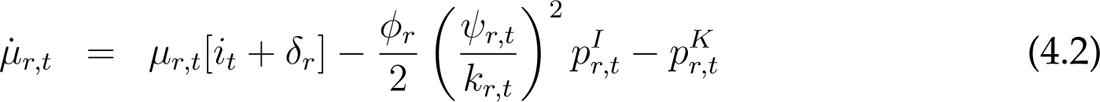

where *p_I_^K^* and *k_r,t_* are the rental price of capital and the capital stock in region *r* at time is the price of an investment good; *δ_r_* is the depreciation rate; *ψ_r_* is the capital increment from the (gross) investment activity; *i_t_* is the global interest rate; *ϕ_r_* is an investment increment coefficient; and *µ_r,t_* is the shadow price of capital.

While the capital accumulation process in a forward-looking model (Equation 4.1) is similar to that of a standard recursive GTAP model, the shadow price of capital equation (Equation 4.2) allows for a connection between future price dynamics to the producer’s current period decision making process. This offers three key advantages. First, the framework can be placed in a large dimensional setting incorporating a large number of produced commodities and countries or regions. This allows model output to capture the full heterogeneity of damages from global warming across countries. Averaging across countries or using small dimensional platforms misses the full distribution of damages and severely distorts the results. Second, forward looking firms (or households) can incorporate future changes into current planning (e.g., emissions reduction targets, a price on carbon, price and output changes from global warming, etc.), rather than ‘forecasting’ next year’s prices/outputs based only on current and past values. This matters greatly for climate change modelling. Finally, the model can be fully integrated with established Representative Concentration Pathway (RCP) and Shared Socioeconomic Pathways (SSPs), connecting global warming to changes in country output and international trade.

In the model, total demand within each region is divided into private and government expenditure, with each of these expenditures further divided into final consumption for each good or service. The final consumption of goods and services together with intermediate and investment demands form a global total demand for a good or service satisfied by production of that good or service in each country. Household income is formed by selling productive factors to producers. Household supply of land, labour and natural resource supply is met by domestic demand, while capital can be transferred to producers in other countries.

In each country/region there are multiple producers representing firms, each of which produces a single good or service employing intermediate goods/services and primary factors: land, labour, capital, and natural resources. Primary factors represent production constraints while serving as inputs to intermediate and final goods. Technological variables influence the degree of productivity at regional, sectoral, and temporal scales. Demand in each region is simulated through a single homogeneous household which consumes domestically produced as well as imported goods and services. The relative proportions of domestically produced and imported goods are influenced by range of factors including trade flows in the base year as well as region- and sector-specific parameters, which remain constant throughout the model run.

The current model utilises the GTAP 10 database (Aguiar *et al*., 2016) and is aggregated to 70 regions and 40 commodities (see Tables 4.1, 4.2) for reasons related to computational feasibility (see e.g., (Kompas & Van Ha, 2019). To run the model a “closure” determines which variables are exogenous (specified by the modeller) and endogenous (allowed to adjust to a new equilibrium). In the case of the current intertemporal model, exogenous variables of interest (e.g., labour productivity) are then provided long-term trajectories representing a deviation from the baseline (see Section 4.1.2) which are consistent with forecasted climate change impacts, effectively simulating the specific scenario of interest (see Figure 4.1). In response to exogenous changes in productivity, endogenous variables including those governing primary production and subsequent capacity for exports (as well as demand for imports) shift accordingly.

**Table 4.1.:**
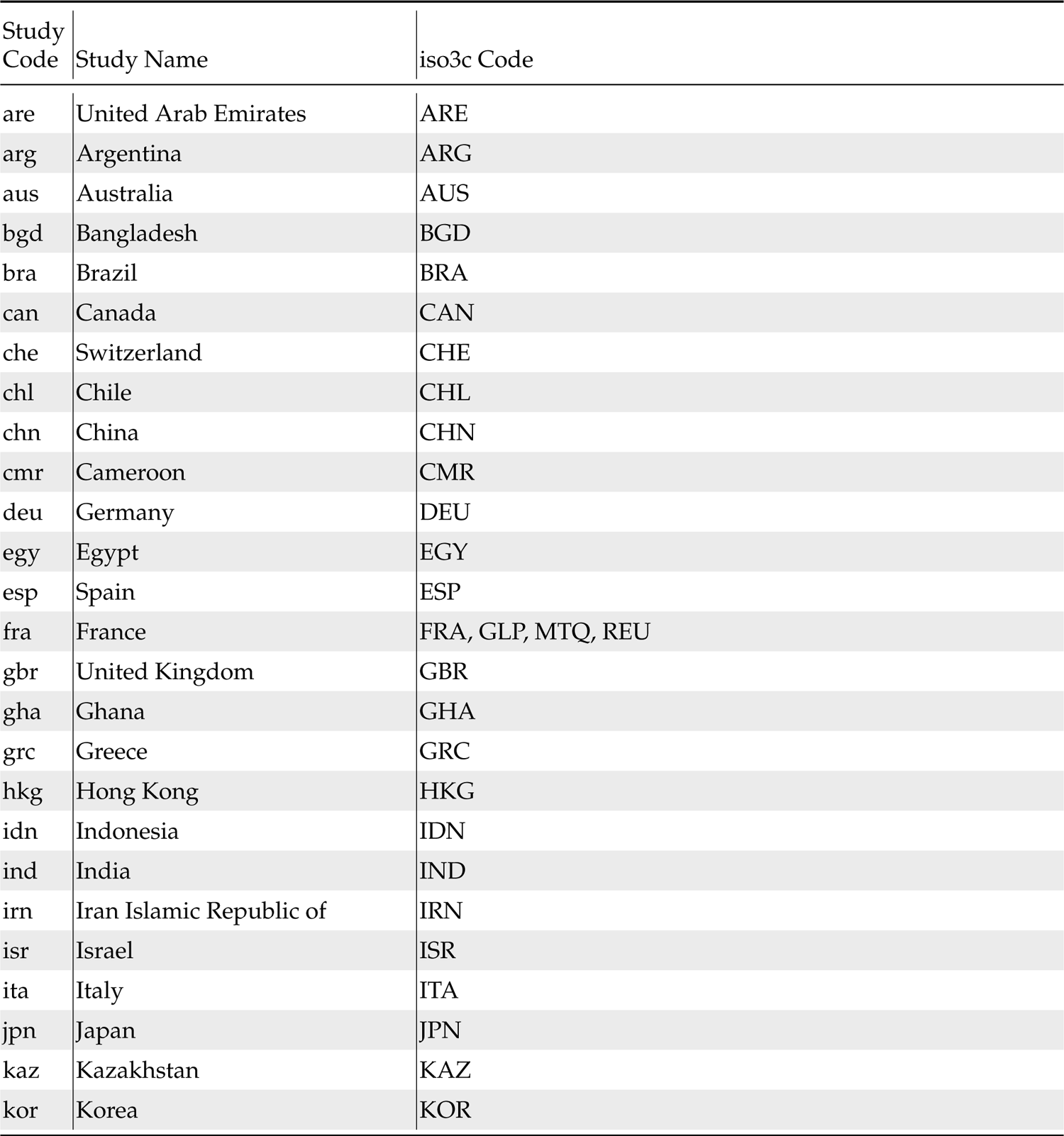
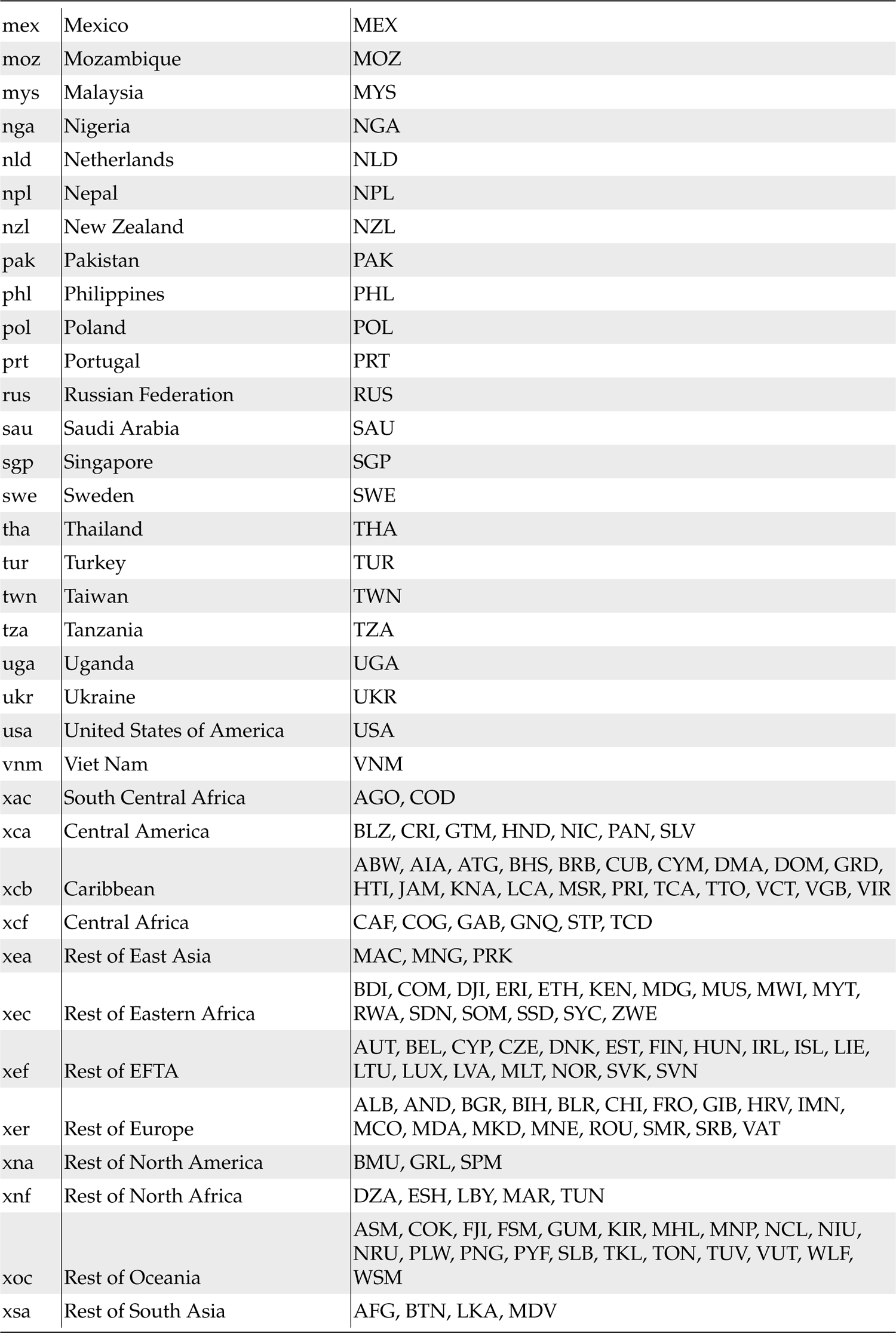
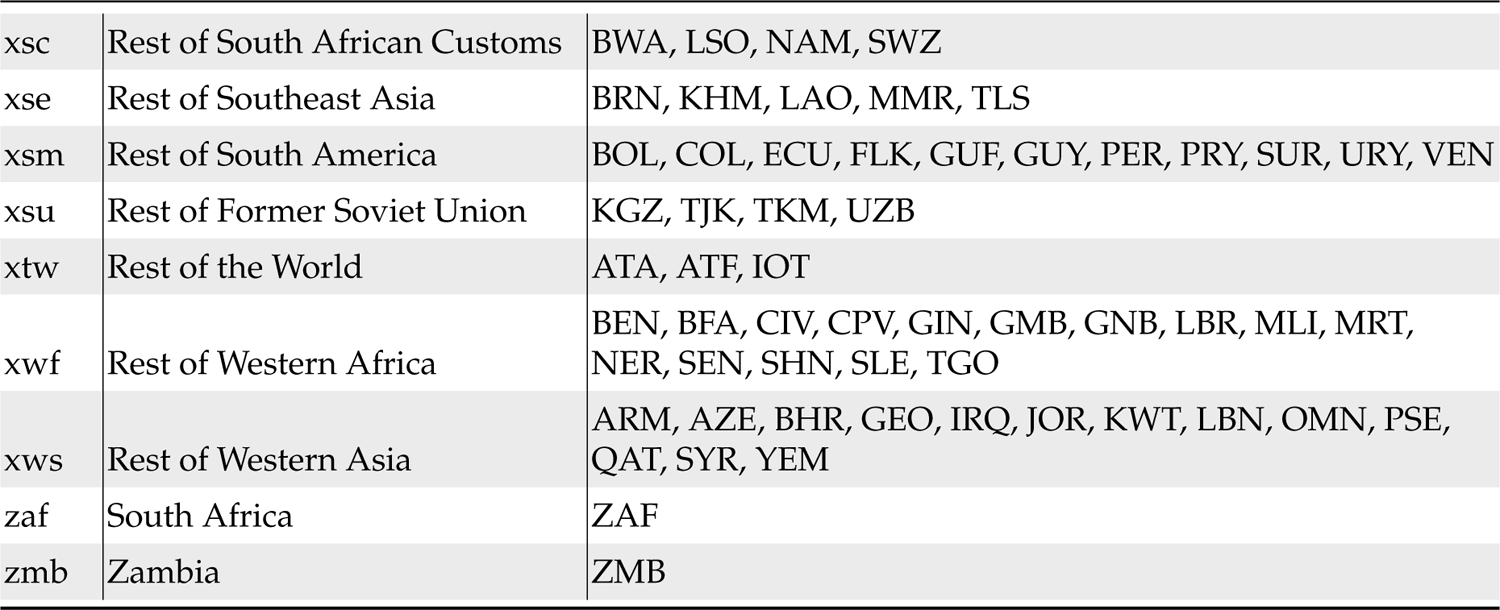
Region mappings: GTAP regional codes and how they map onto ISO3C country codes and the aggregated regions used in this study.

**Table 4.2.:**
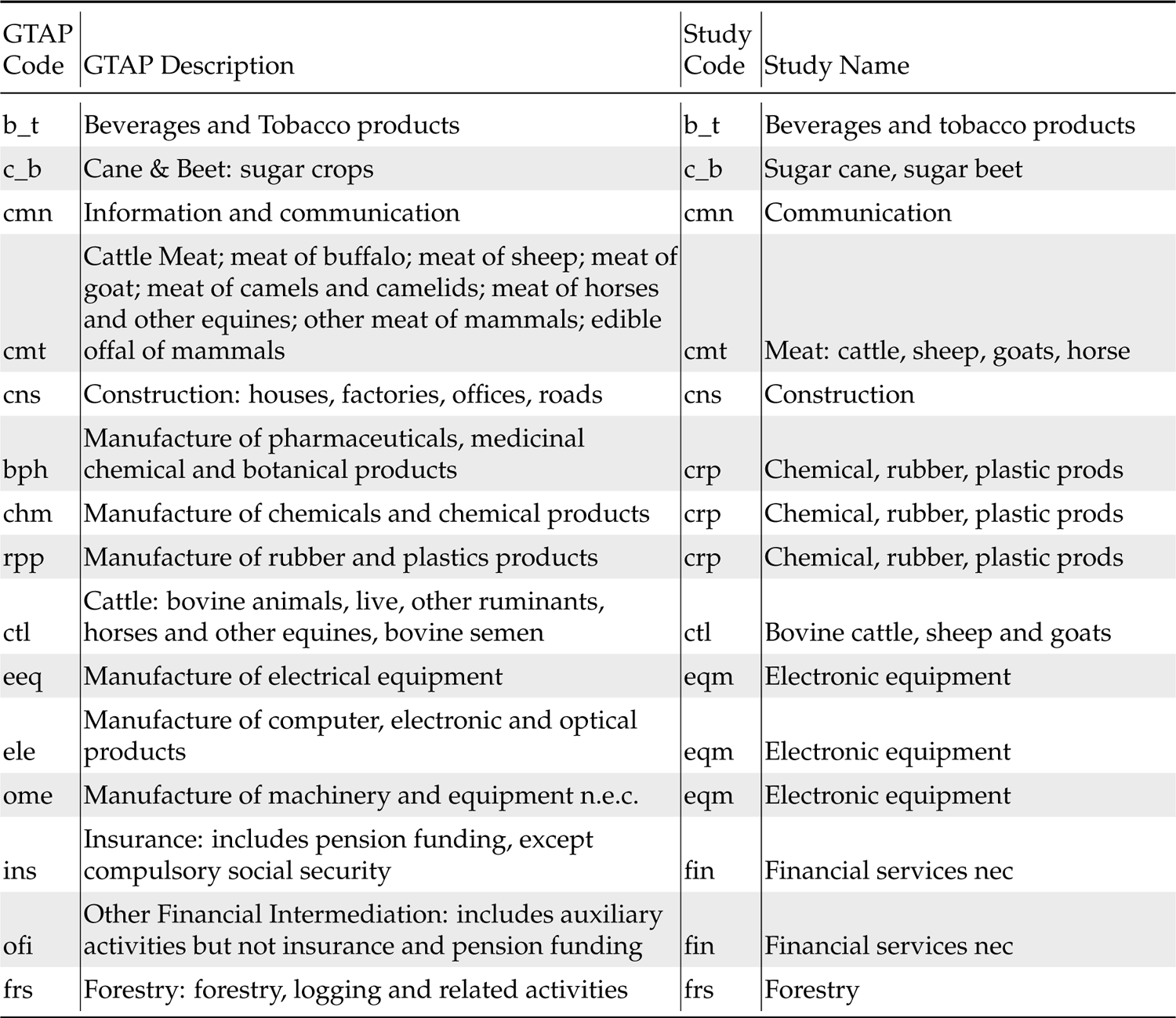
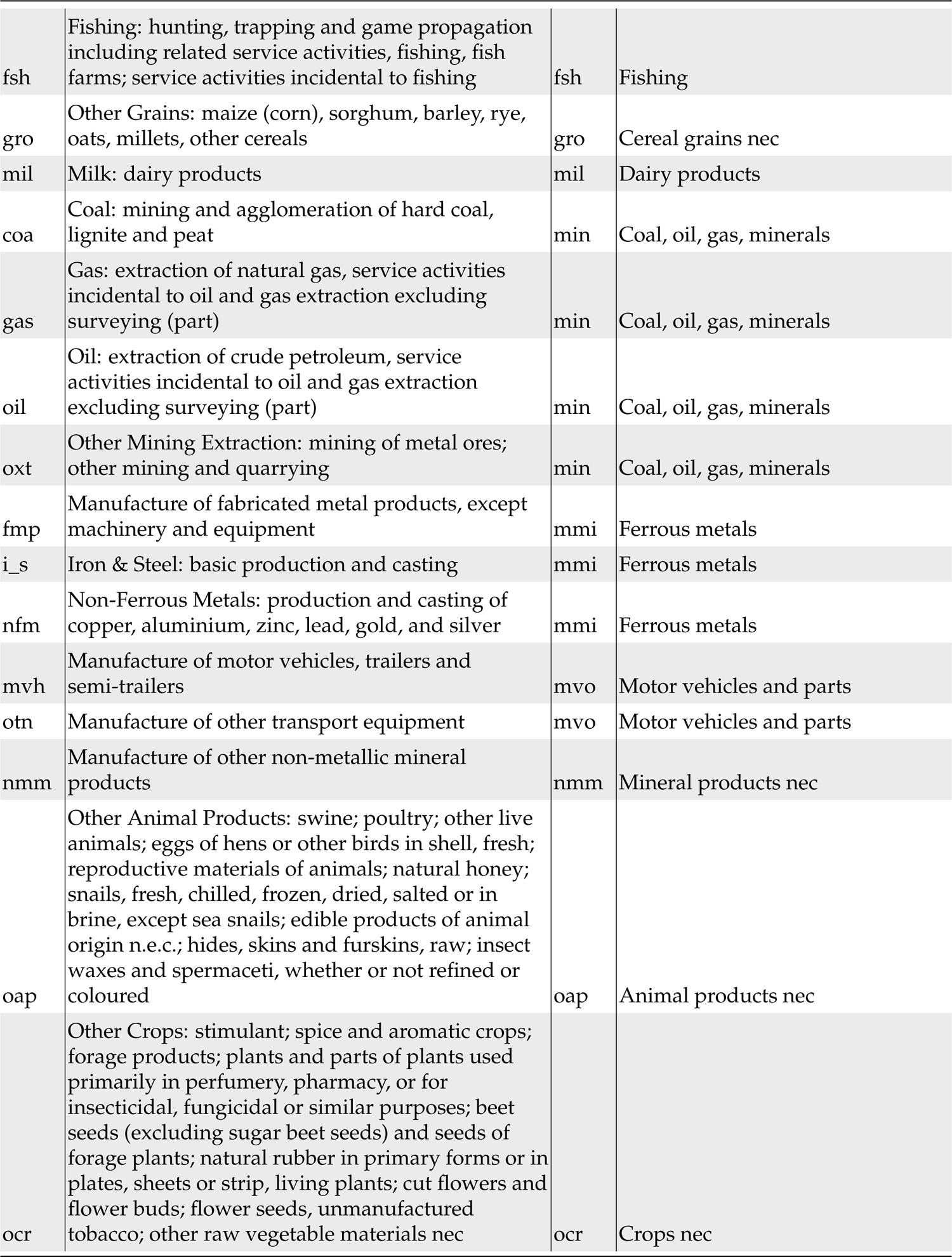
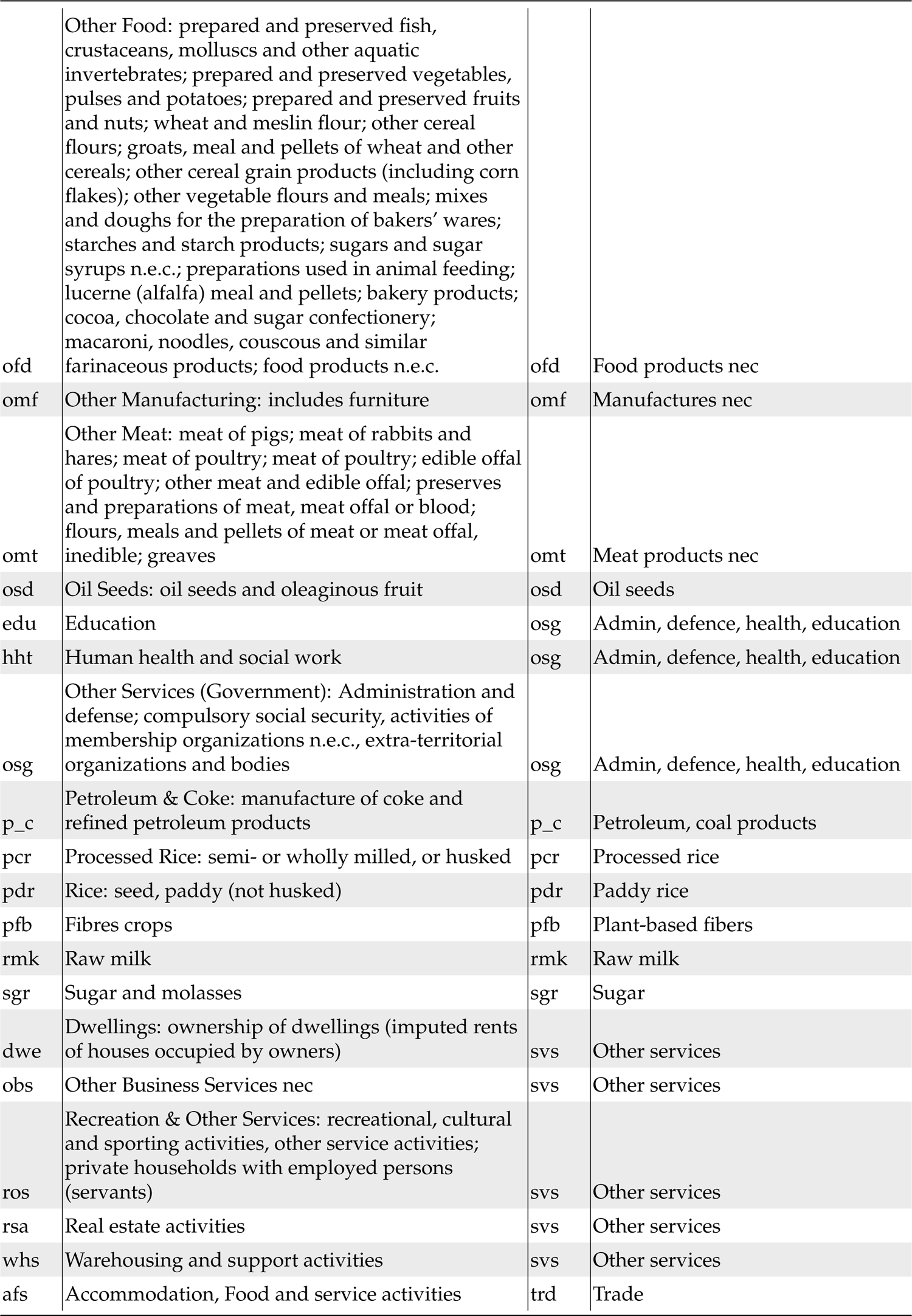
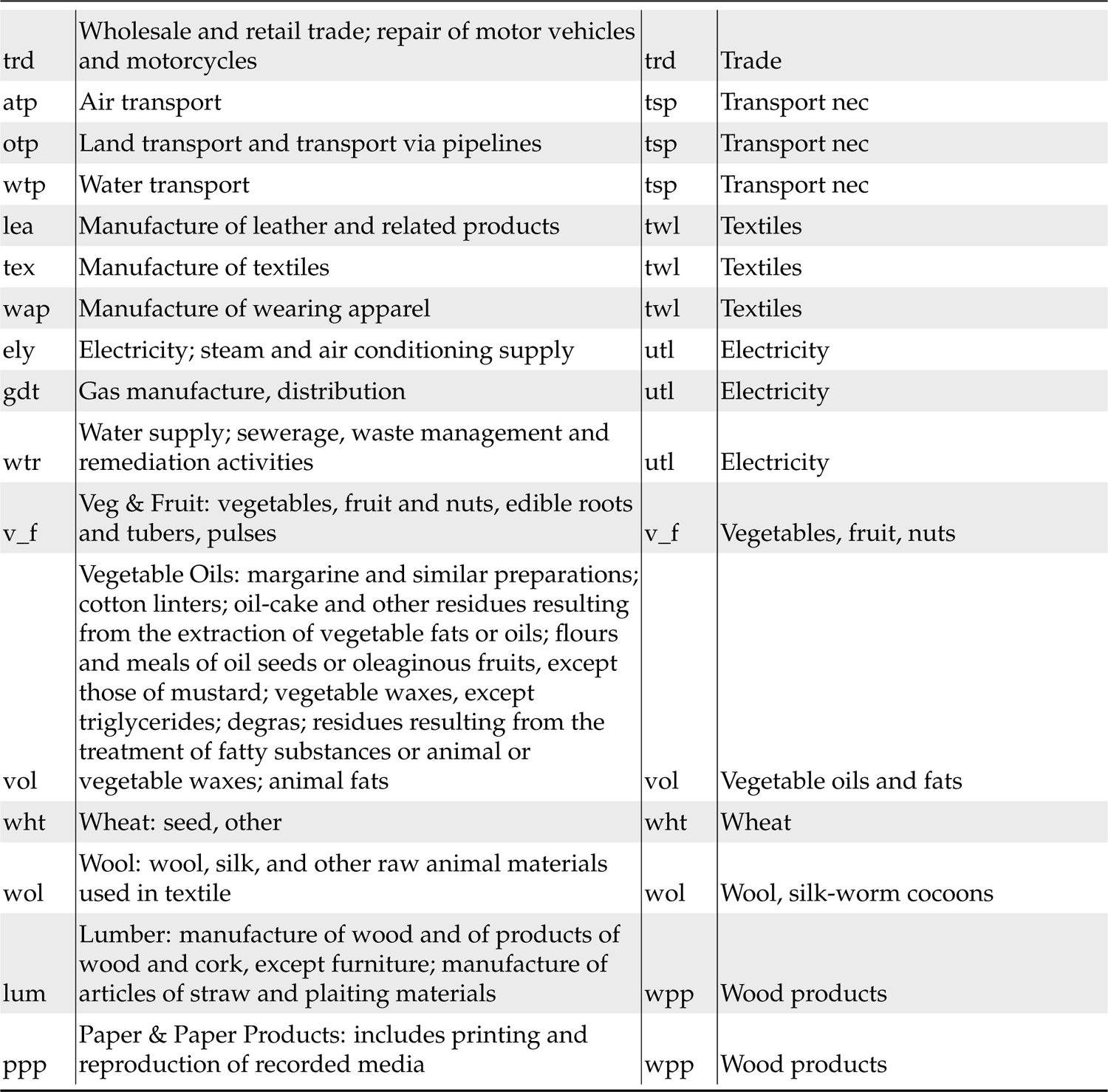
Sectoral mappings: GTAP sectorial codes and descriptions and how they correspond aggregated commodity sectors used in this study (i.e. Study codes).

### 4.1. Simulating climate change impacts

Climate change damages encompass nearly all aspects of the economy and daily life. There are many uncertainties associated with the identification, quantification, and modelling of known impacts within the context of an economic model such as that used in this analysis. Given these challenges we adopt a conservative approach and implement three important broad impact channels from climate change (Roson & Sartori (2016) and Li & Kompas (2022)): 1) limited sea level rise (losses in arable land), 2) heat stress losses on agricultural productivity and labour productivity, and 3) crop productivity losses. State-change events (i.e., tipping points) which may drastically alter the economic landscape are not modelled within the current framework. Due to limitations in the quantification of the many vectors through which climate change damages may manifest into economic losses, we cautiously recommend that results (Chapter 5) be interpreted as “optimistic”.

#### 4.1.1. Climate change scenarios

The Representative Concentration Pathways (RCPs) present a goal-oriented normative scenario approach to envisioning potential emissions pathways consistent with a prescribed radiative forcing ranging from 2.6 to 8.5 W/m^2^ (Moss *et al*., 2010). RCP4.5 emissions peak at 650 ppm CO_2_ equivalent (Thomson *et al*., 2011) while RCP 8.5 represents a future characterized by high fossil fuel use and 1350 ppm CO_2_ equivalent (Riahi *et al*., 2011). RCP 4.5 and 8.5 temperature paths can be simulated by MAGICC model^2^ (Meinshausen *et al*., 2011a) up to 2100. MAGICC is what is known as a reduced complexity climate model (see Nicholls *et al*., 2020) and allows for less computationally demanding model runs compared to more complex Earth System Models. MAGICC’s default parameters are used within the current analysis (climate change parameters from CMPI3 and carbon cycle setting from C4MIP).

For temperature paths from 2100 and beyond, we follow Kompas & Ha (2021) and use a 2-step approximation process. First, we approximate the unadjusted temperature path using a two layers (atmosphere and lower ocean layer) temperature model following Glotter *et al*. (2014). We also use the mid year radiaive forcing data from Riahi *et al*. (2007); Meinshausen *et al*. (2011c,b) as inputs^3^. In the second step, we adjust the above approximated temperature path to resemble the middle temperature profile for RCP4.5 and RCP8.5 scenarios in (Meinshausen *et al*., 2011b; Nazarenko *et al*., 2015; Nauels *et al*., 2017). Long-term temperature paths for each RCP are mapped to each region within the study (Table 4.1). For consistency with Roson & Sartori (2016) results, we use linear extrapolation to map the point damage estimation to the RCP8.5 temperature path to calculate the shock values beyond 2100.

#### 4.1.2. Damage functions and shocks

All damages are calculated directly from Roson & Sartori (2016) with the exception of the agricultural productivity shocks found in Li & Kompas (2022) (see Figure 4.1). Damages are implemented through the use of “shocks” within the CGE model, meaning that trajectories are directly provided to the corresponding variables within the model (e.g., land supply). Losses at 2050 and 2100 (see Roson & Sartori (2016); Appendix Tables) are linearly interpolated over each RCP temperature paths and mapped to time steps within the model, resulting in a smooth trajectory and eventual plateau for each shock (far beyond 2100). Within the current analysis there is no “business-as-usual” counterfactual future with zero climate change impacts therefore all CGE model results can be interpreted as the change between data in the reference year and a climate-change enriched trajectory.

**Figure 4.1.:**
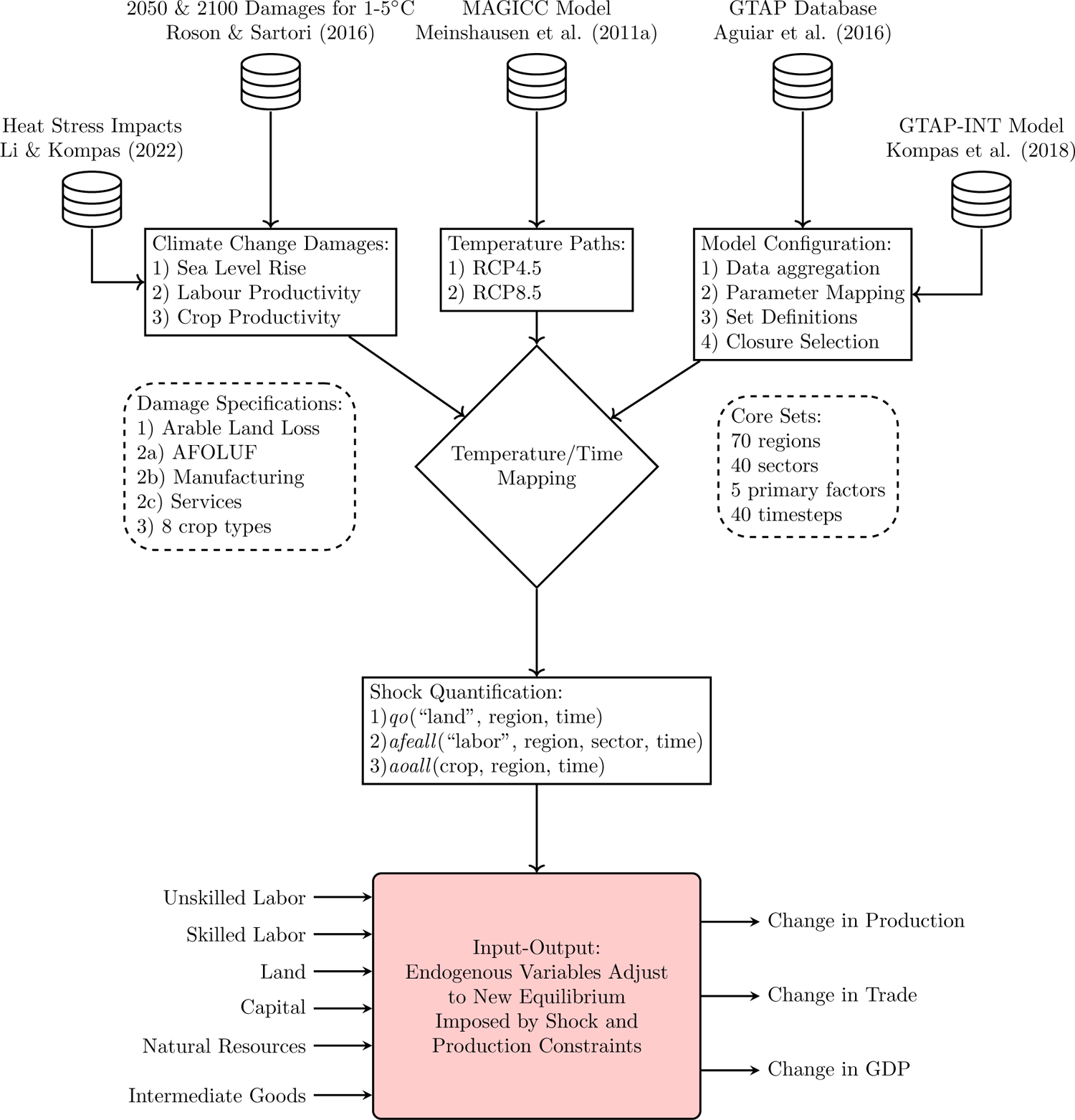
CGE modelling framework. AFOLUF represents agriculture, forestry, other land-use, and fishing.

### Sea level rise

Sea level rise impact on *arable land* in Australia is insignificant compared with other countries, with Roson & Sartori (2016) predicting only a 0.005% loss of arable land in Australia under the 5°C increase scenario by 2100. This estimate corresponds with the relatively low amount of agricultural land situated in areas susceptible to sea level rise. Loss of arable land due to sea level rise is simulated directly through shocks to land supply (variable *qo*) which is indexed to the land primary factor for each region at all time steps. Within the current analysis we do not examine the impact of sea level rise or storm surge on other industries such as manufacturing nor are the costs of protecting urban areas taken into consideration. These can be considerable. Similarly, impacts on coastal biodiversity, losses in residential, commercial, and industrial lands and ecosystem services are also not considered.

### Labour productivity impacts

In addition to directly targeting crop productivity, the current model simulates labour productivity losses in all model sectors. This is accomplished by reducing the efficiency of both skilled and unskilled labour as primary factors via the skilled and unskilled labour elements within the *afeall* variable, which represents primary factor augmenting technological change. For labour productivity with Australia, Roson & Sartori (2016) predict a loss of 5.55% at 5 °C for agricultural sectors and near zero losses within manufacturing and service sectors (1-5°C). Dedicated shocks are delivered to services, manufacturing, and agriculture, forestry, and fishing sectors (Figure 4.2), with land-use based services in regions most susceptible to climate change experiencing the greatest losses.

**Figure 4.2.:**
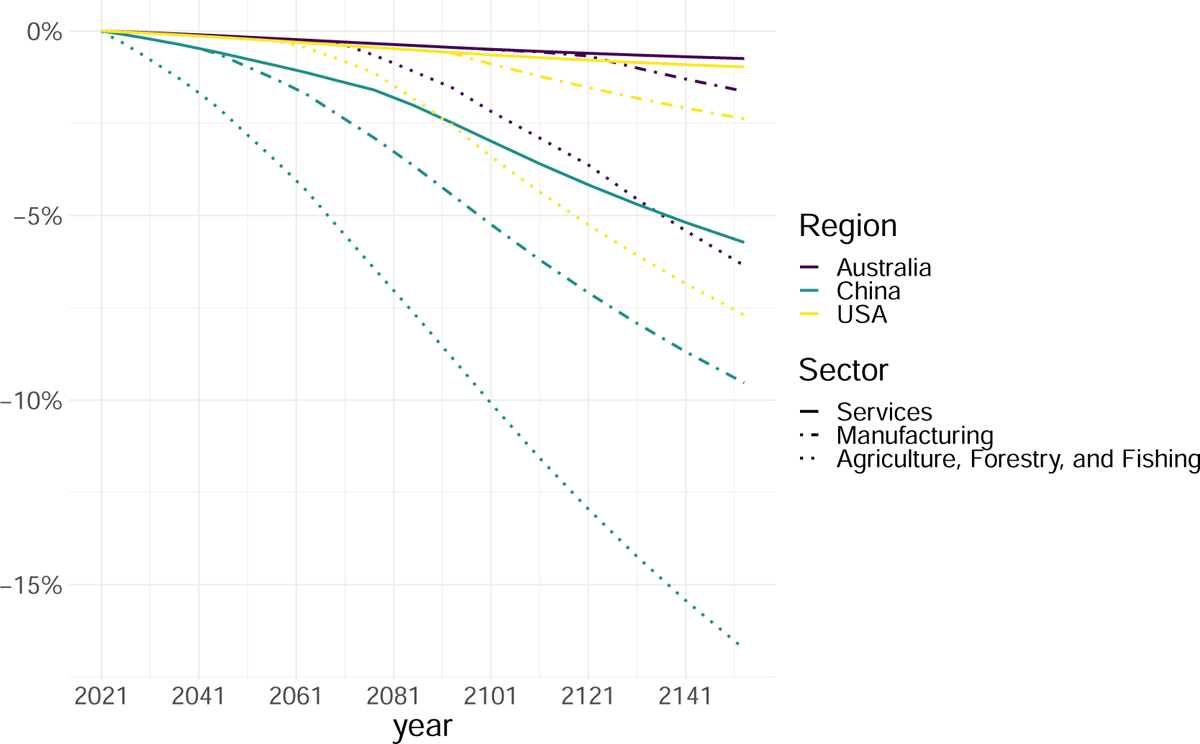
Sectoral labour productivity shocks for select regions under RCP8.5

Overall, with these limited damage functions from global warming, the impacts to Australia are small or very limited. However, impacts throughout the world are not small, with losses in national GDP ranging from marginally positive to −23% or more, depending on time horizon. It is these changes in global economies, with considerable heterogeneous impacts, that generate the substantial changes in the pattern of global imports and exports.

### Crop productivity impacts

The current study implements shocks to agricultural productivity (i.e., crop yield) which are produced through the use of temperature-response functions calibrated with national average impacts of climate change. The model distinguishes between 8 crop classes within the GTAP databases with dedicated shocks to wheat, rice, cereals and grains, oil seeds, and other crops.

In this paper, we use the estimation of crop productivity loss under five climate change scenarios from Li & Kompas (2022). Crop productivity shocks are carried out on the *aoall* variable which represents output augmenting technical change, indexed across all crop classes by region and time step. Specifically, we performed a meta-analysis of yield responses to temperature, precipitation and CO_2_ using the Consultative Group for International Agricultural Research (CGIAR) dataset coupled with mixed-effect models to account for correlation bias from multi-estimate studies. We fitted five models to future projections of temperature and precipitation from the sixth Coupled Model Intercomparison Project (CMIP6) ensemble for three Shared Socioeconomic Pathways (SSPs) and Representation Concentration Pathways (RCPs) on a 0.5*^◦^* grid. We systematically evaluated uncertainty attributable to model choice, sampling, missing data and the CMIP6 ensemble. We found larger yield losses for maize and rice in many regions than previously estimated in the literature, implying larger direct effects of climate change on agricultural productivity.

The global production-weighted mean percentage yield losses for maize at RCP8.5 is −12.9% with a range of −15 to −10%. For rice the global mean is −2.7% [−9, 1], for soy −17% [−40, 3] and for wheat −32% [−64, −6]. Mean results and ranges for RCP4.5 for maize, rice, soy and wheat are −11.6% [−13, −9], −1.1 [−4, 0.8], 1.1% [−10, 12], and −11% [−28, 1]. The upper range of damages occur in Africa, Southeast Asia and South Asia, where heat stress is largest.

## 5. Predicted global damages from climate change

Table 5.1 shows the economic impact of temperature increase on GDP under the RCP4.5 and RCP8.5 scenarios. Here it should be noted once again that these damages are limited to changes to labour and crop productivity as well as loss of arable coastal land. Western Africa, South Asia, and South East Asia are the most heavily impacted areas with losses as large as 20%+ GDP observed by 2100 under the RCP8.5 scenario. In the case of the less severe temperature increase scenario (RCP4.5), GDP damages from climate change reach 6-7% in 2100. Australia fares relatively well^1^ in terms of the direct climate change impacts simulated within this analysis, however, many key exporters within South and Southeast Asia experience substantial losses toward the end of the century.

**Table 5.1.:**
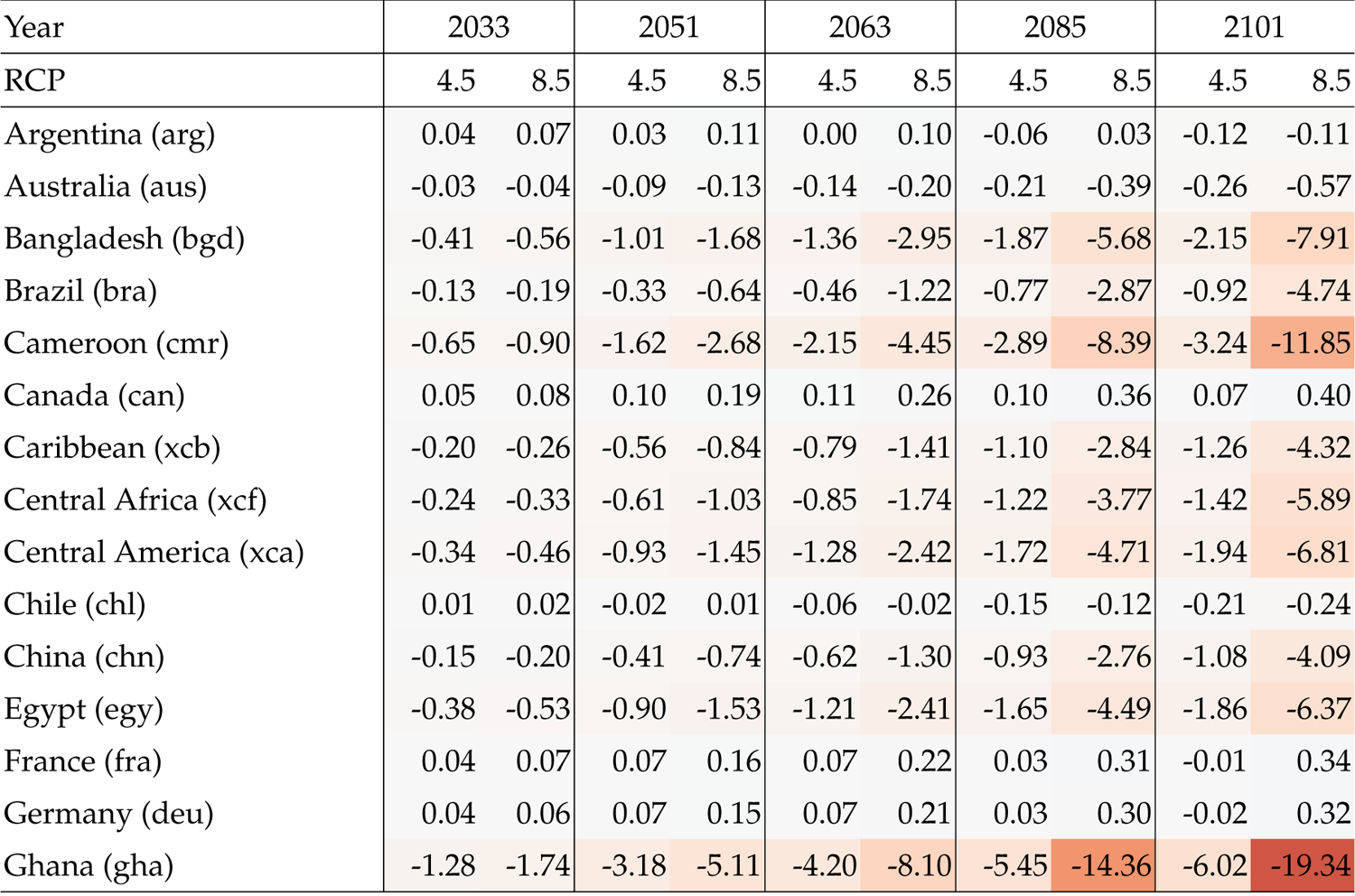
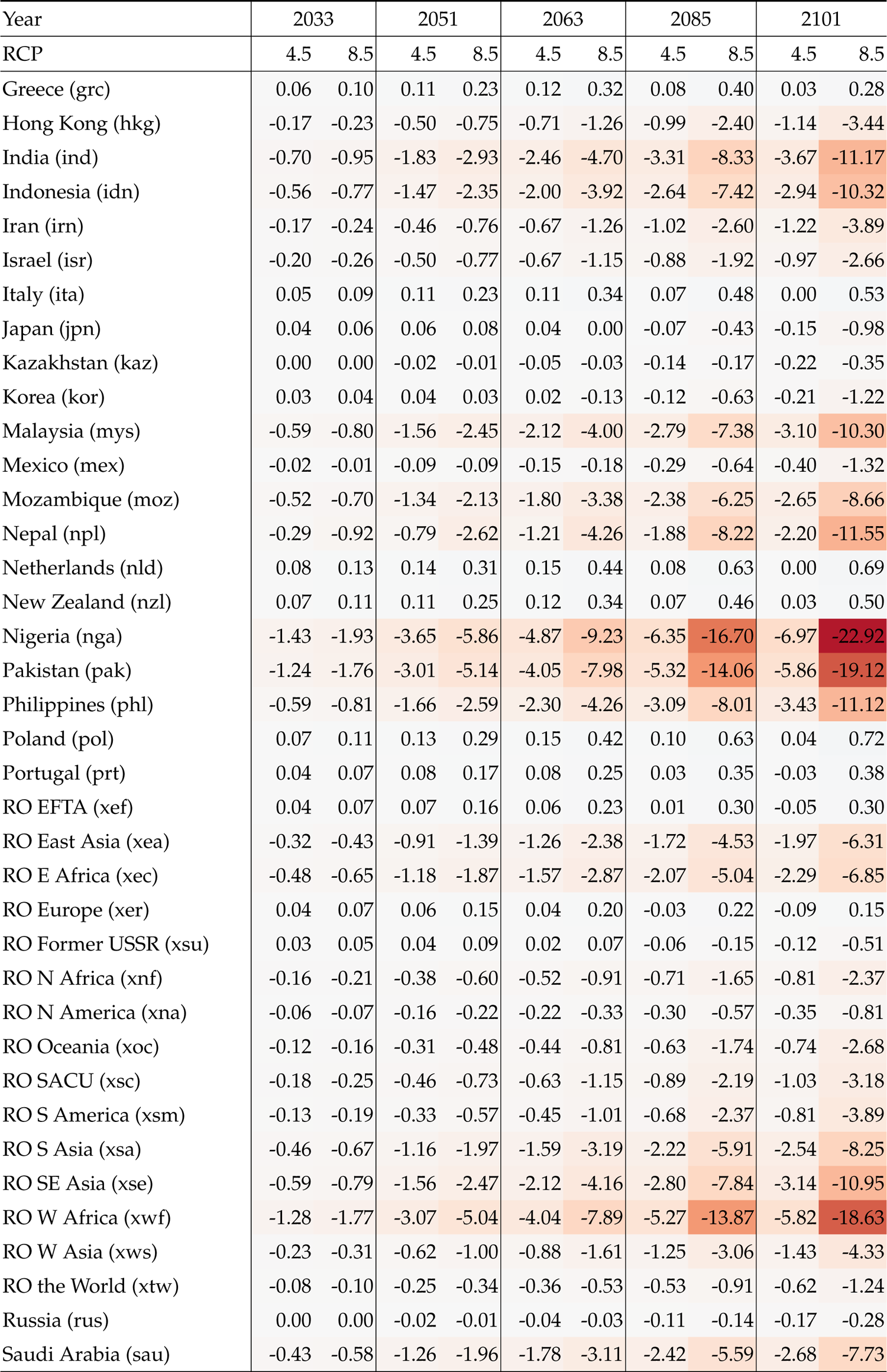
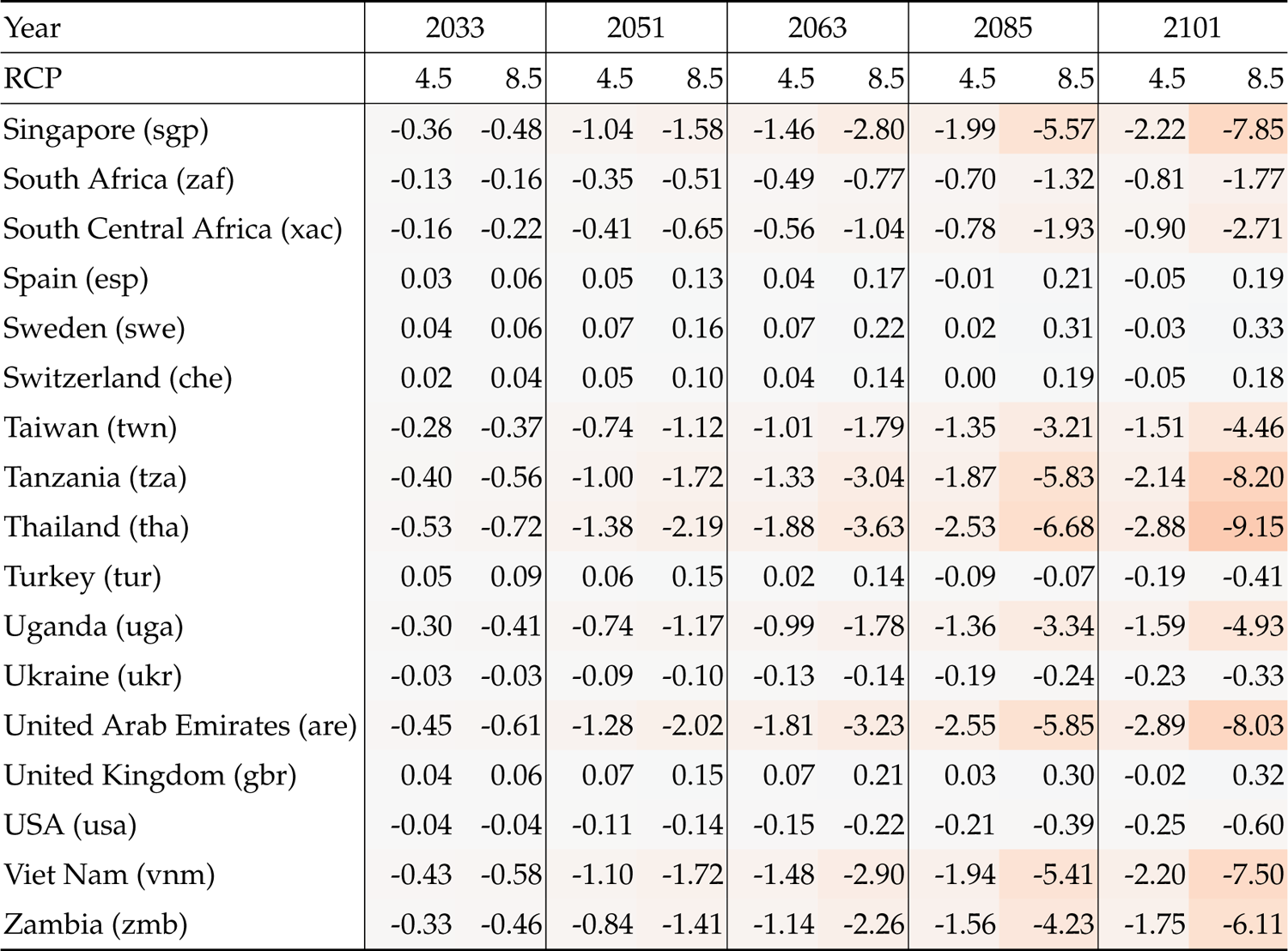
GDP impact due to crop and labour productivity losses (% change) for different regions under both RCP4.5 and RCP8.5. RO stands for “Rest of”. Darker shading indicates greater negative climate change impacts.

### 5.1. Australia’s trade future under climate change

Within this section we break down changes in Australia imports by macro industrial sector followed by a closer examination of individual sectors and their key exporters. Finally, we take a look at what is not currently included within the model and briefly touch upon some of the known sources of uncertainty in variables driving this analysis. Drops in imports are spatially concentrated in the same regions which directly experience climate change damages (i.e., South and Southeast Asia).

China remains the largest exporting nation despite undergoing a 3% drop in imports under RCP8.5 by 2100 while the general configuration of exporting nations remains the same with the exception that Thai exports are surpassed by South Korean exports around the 2070 time step.

Results from the current modelling exercise suggest that Australian imports will experience a wide range of impacts under climate change at a highly aggregated level between manufacturing, services, and agriculture as well as heterogeneous impacts within these sectors. In terms of overarching economic impacts, the largest absolute decreases in Australian imports were observed within manufacturing sectors where the total value of imports in 2101 fall below current levels for both RCP4.5 (−0.6%) and RCP8.5 (−1.9%). Similarly, service industries fall −0.5% and −1.5% by 2101 under RCP4.5 and RCP8.5 respectively. In contrast to manufacturing and service sectors, we see considerable variation in crop sectors imports in large part due to the varied mix of exporters for each crop type and differential impacts experienced. Turning to livestock, forestry and other non-agricultural land-use sectors we see smaller climate change impacts, primarily due to the largest exporters being situated outside of the global zones that experience the greatest damages from global warming.

In terms of Australian exports, the greatest absolute decreases occur with respect to China, India, and Southeast Asia while Pakistan experiences the greatest relative drop at −23%. The greatest absolute drops correspond to the Coal, oil, gas, and minerals sector (`min`) and wheat (`wht`).

In order to provide greater transparency into model results, an interactive dashboard was constructed which allows for a closer examination of trade data by region, sector, and time-step (Figure 5.1). The left side of the dashboard responds to user inputs over exporter(s), importer, commodity, scenario, and target year. For any number of select exporters, the line plot (top middle pane) shows the change in traded value to the select importer over time by each climate scenario. The bar plot (top right pane) responds to the selection of importer, commodity, and year – providing insight into the top exporters at each time step. Finally, the world map (bottom pane) uses a heatmap calculated on the square root of the absolute deviation between the selected year and 2014 by commodity. Although absolute change should be interpreted with some caution due to the unpredictable nature of currencies over such a long span of time, both absolute (in 2014 USD) and relative deviations can be accessed by clicking on a region of interest.

**Figure 5.1.:**
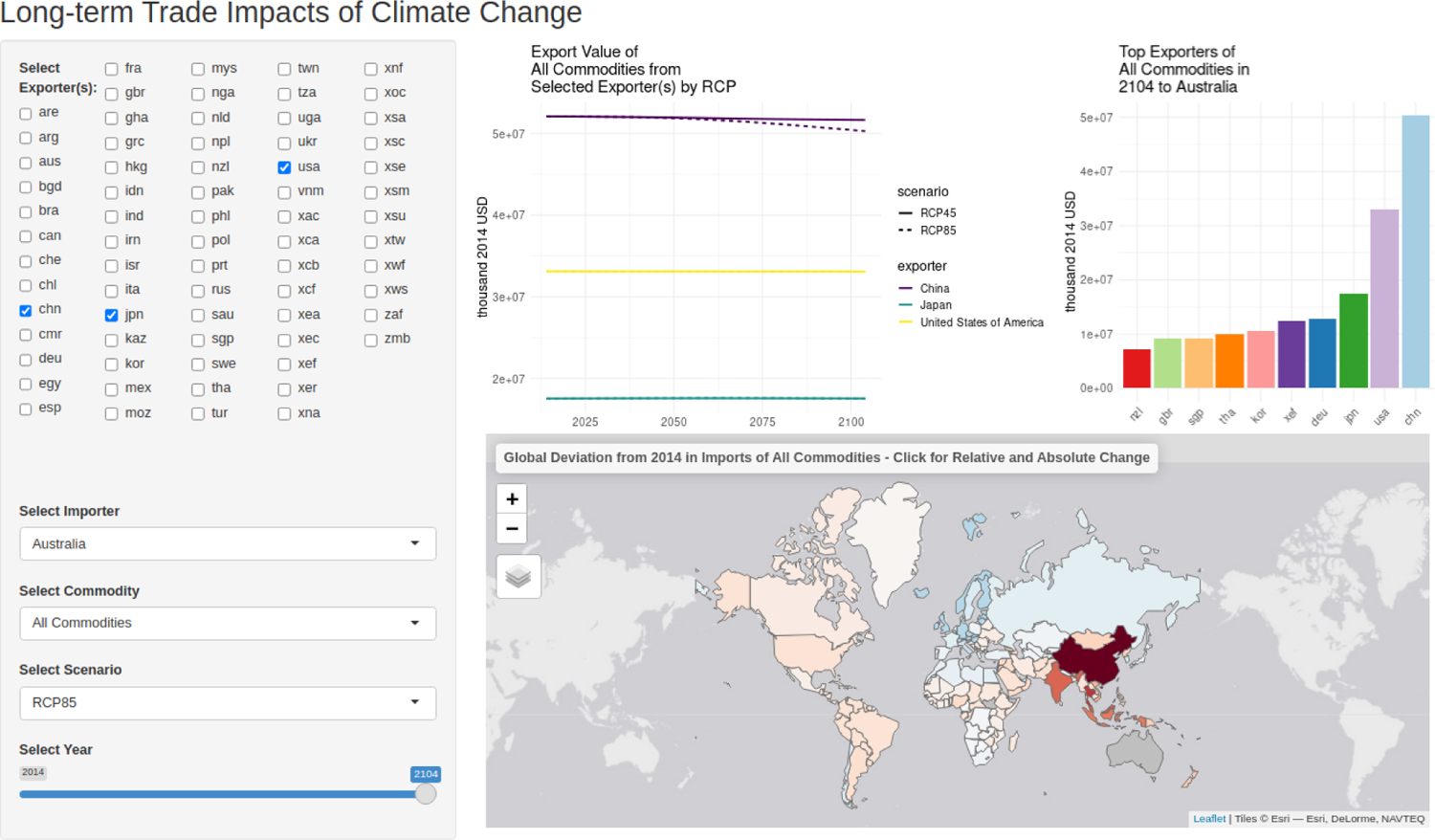
Interactive dashboard screenshot from: https://apps.cebra.unimelb.edu.au/trade-dashboard/.

#### 5.1.1. Manufacturing imports

Manufacturing imports from East, South, and Southeast Asia experience the greatest drops due to climate change impacts in this region (Figure 5.2). Within manufacturing, *Motor vehicles and parts* and *Electronic equipment* are the most heavily impacted, with China, Thailand, and Malaysia bearing the brunt of labour productivity losses. Although 68% of exporter/export combinations experience a loss, *Electronic equipment* from the USA and Germany see small increases throughout the model run. As the largest exporter of manufacturing goods and a key region affected by climate change, China sees the largest decline in exports, however most of South and Southeast Asia fares worse in terms of relative losses with India and Indonesia seeing over 10% losses. Conversely, net gains are made across several European regions, particularly Germany and the UK.

**Figure 5.2.:**
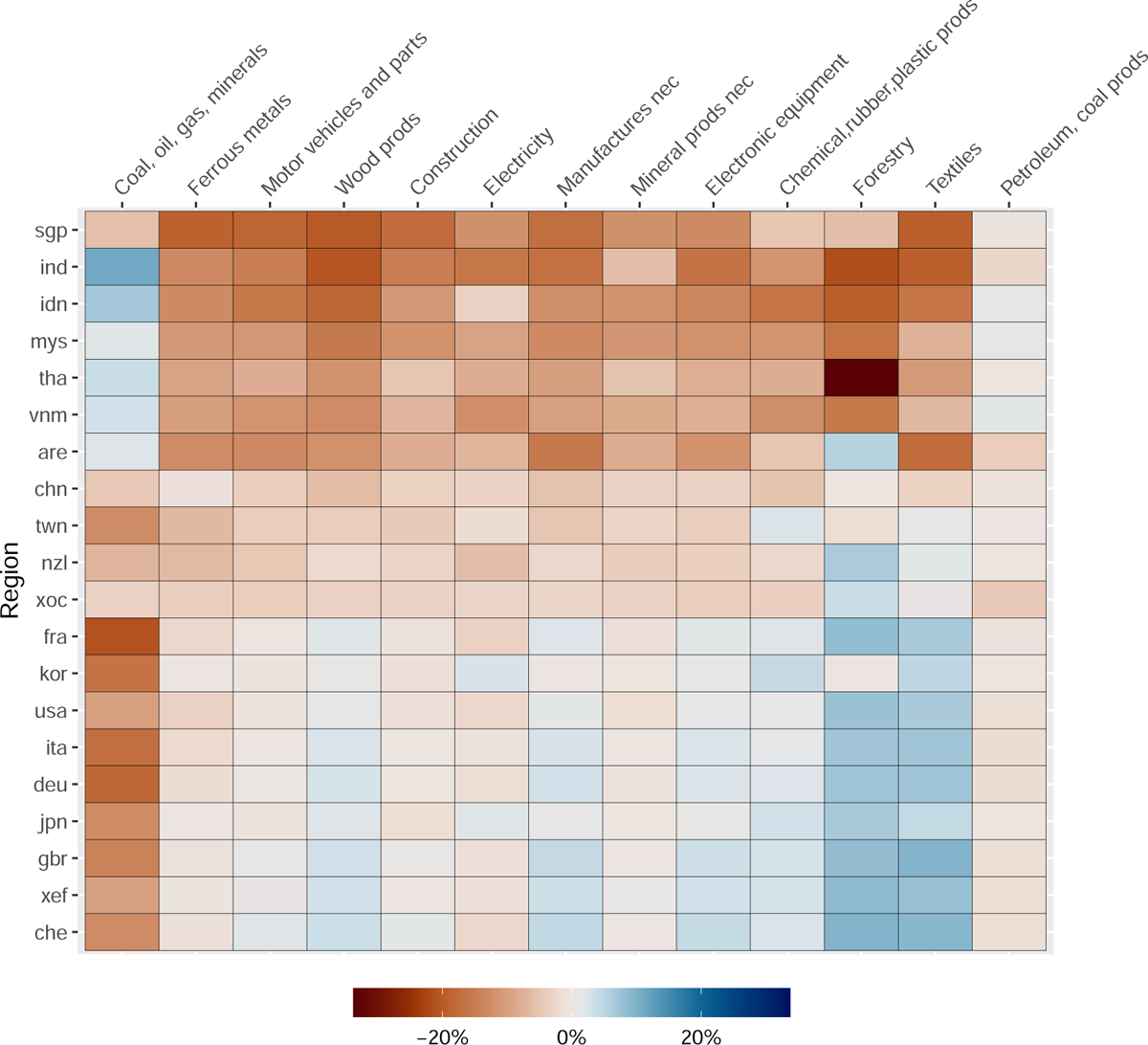
Change in import dynamic among the top 20 exporters of manufacturing goods to Australia under RCP85 in 2104. Axis ordering based on the (unweighted) mean change in value across all regions/sectors.

#### 5.1.2. Agricultural and other land-use intensive imports

In contrast to manufacturing imports which on the whole decline due to heat stress impacts on labour productivity, change in agricultural imports and in particular crop sectors exhibit a greater range both in terms of exporting regions and imports. The largest absolute increase among food- and agriculture-related sectors is seen within *Vegetables, fruit, nuts* with large increases in exports from OECD regions and China (Figure 5.3). All other sectors within this category show either negligible or net decreases in imports with *Other Food* (consisting of a variety of prepared and processed foods) experiencing the largest absolute losses, chiefly borne by Thailand and China. In relative terms we see increases for much of Western Europe while South and Southeast Asia experience losses. Given the large number of commodities modelled and wide range of variables considered, we recommend that users access the interactive dashboard to gain greater insight into the nature of these changes through 2104.

**Figure 5.3.:**
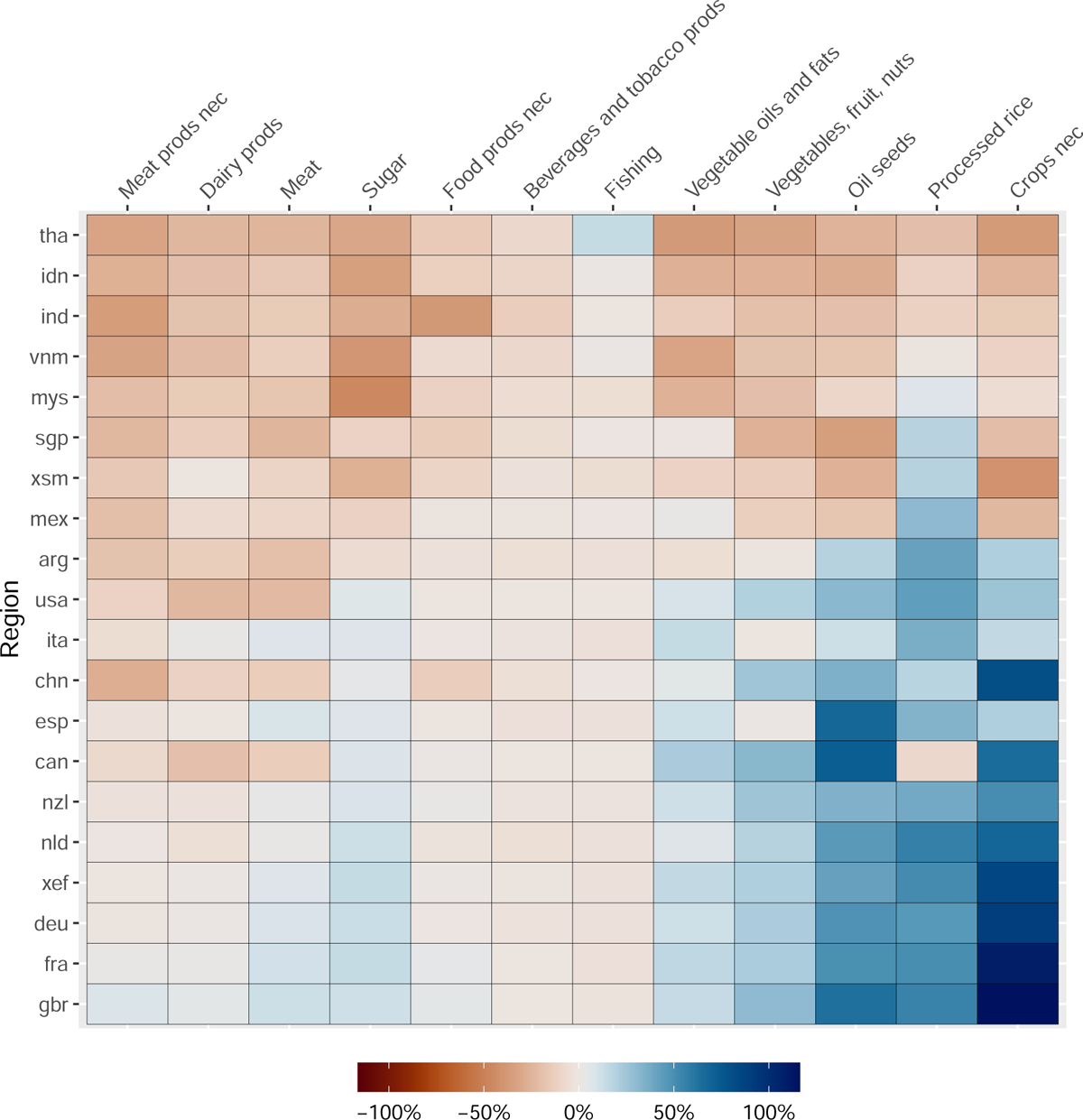
Relative change in import dynamic among the top 20 exporters of food and agricultural goods to Australia under RCP85 in 2104. Food and agricultural goods here are limited to the top 12 sectors by imported value. Axis ordering based on the (unweighted) mean change in value across all regions/sectors.

## 6. Modelling contamination rates, propagule pressure & establishment exposure

To determine the implications of changing climate and trade flows on region-specific propagule pressure and establishment exposure of hitch-hiking threats, we have extended an existing model developed by the Centre of Excellence for Biosecurity Risk Analysis (CEBRA; Camac *et al*., 2021a) by integrating it with spatio-temporal projected changes in: 1) international trade flows (derived from GTAP simulations, see Chapters 4 & 5); 2) threat climate suitability; and 3) human population, under different future climate and climatic change scenarios.

In the following sections, we briefly outline the data obtained from the Australian Department of Agriculture Fisheries and Forestry (DAFF), how these data were used to estimate trading partner by commodity contamination likelihoods, and ultimately, region-specific temporal dynamics in threat propagule pressure and establishment exposure. We illustrate the implementation of this model using the five priority hitch-hiking plant pests selected by the Department of Agriculture, Fisheries and Forestry: brown marmorated stink bug (*Halyomorpha halys*), spongy moth (*Lymantria dispar*), Asian honey bee (*Apis cerana*), giant African snail (*Lissachatina fulica*) and Khapra beetle (*Trogoderma granarium*). These five threats were selected as they are each deemed priority plant pests and all are known hitchhikers. Moreover, they were selected as each is an exemplar of a different hitch-hiking functional group defined by the Department of Agriculture, Fisheries and Forestry. Specifically:

- **Overwintering** – brown marmorated stink bug (BMSB; *Halyomorpha halys*): Pests that have biological attributes that result in them aggregating in large numbers over winter.
- **Egg laying** – Spongy moth (*Lymantria dispar*): Pests that commonly lay eggs on structures or exported goods.
- **Nesting** – Asian honey bee (*Apis cerana*): Pests that commonly build nests on structures or exported goods.
- **Sheltering** – Giant African snail (*Lissachatina fulica*): Pests that commonly take shelter on structures or exported goods.
- **Internal storage** – Khapra beetle (*Trogoderma granarium*): Pests that commonly occurs and persists in internal environments such as shipping containers and their associated goods.

### 6.1. Interception & import data

To estimate threat-specific commodity contamination likelihoods for different regions, the Australian department of Agriculture Fisheries and Forestry (DAFF) supplied CE-BRA with all interception records available in the Agriculture Import Management System (AIMS) database between 2010-01-01 and 2022-06-02. Each record contained a incident ID, entry ID and date, the imported commodity type (i.e., 6-digit Harmonised System code, also known as a HS code) the contaminant was detected on, and the country the imported item was believed to originated from^1^. Where possible, biological contaminants were identified to species. However, if this was not possible, identification was made to the lowest possible taxonomic levels (i.e., genus, family or order).

Interception data were then processed by removing 1) duplicate records (i.e. those with same taxon ID, entry ID, date and commodity type); 2) records prior to 2014 (the earliest projected year in the GTAP modelling) and after 2021; 3) records that could not be attributed to a HS commodity code; and 4) records that contained an unknown country of origin or were attributable to a country with no known established population. Data were then subset to records containing the five threats of interest: brown marmorated stink bug (*Halyomorpha halys*), spongy moth (*Lymantria dispar*), Asian honey bee (*Apis cerana*), giant African snail (*Lissachatina fulica*) and Khapra beetle (*Trogoderma granarium*). Note, as spongy moth was not always identified to species-level, as such, we also included all genus records (coded as “(Lymantria)” in the interception database).

The department of Agriculture Fisheries and Forestry also provided CEBRA with annual summary statistics on the number of imported lines inspected (derived from AIMs data) and the total number of lines imported into Australia (derived from Integrated Cargo System - ICS) for each 4-digit HS commodity code and country of origin for the same period as the interception dataset. These data are critical to quantify: 1) contamination rates (i.e., the number of lines contaminated relative to the total number of lines inspected), and 2) the total number of commodity lines imported into Australia for each country of origin by commodity and year combination – acknowledging that most imports have less than 100% inspection rates.

To predict changes in contamination rates over time and under different climate change and climate scenarios, we first standardise all HS codes to a single HS version (HS version 5), we then mapped these codes in both datasets to the 31 commonly traded commodity sectors^2^ used in the GTAP inter-temporal trade flow model (Table 4.2). This was achieved using a HS (version 5) to GTAP 10 sector concordance table.^3^ Country codes were also mapped onto the 70 GTAP global regions used in this study (Table 4.1). Statistics such as the number of interceptions of each threat, the number of line inspections, and the number of line imports were then aggregated (i.e. summed) for GTAP region, GTAP commodity sector, and year. The interception and import datasets were then merged into a single dataset containing: numbers of threat-specific interceptions, number of line inspections and total number of lines imported into Australia for each GTAP sector by region by year combination.

Out of the five exemplar threats, brown marmorated stink bug was the most intercepted threat between 2014 and 2021, this was followed by giant African snail, Khapra beetle, Asian honey bee and lastly, spongy moth (Table 6.1).

**Table 6.1.:** Summary of threat interceptions as well as the number of GTAP regions and commodity sectors they were detected at Australian borders between 2014 and 2021.

| Threat | Interceptions | Regions | Commodities |
| --- | --- | --- | --- |
| Halyomorpha halys | 1407 | 15 | 13 |
| Lissachatina fulica | 107 | 14 | 11 |
| Trogoderma granarium | 52 | 4 | 4 |
| Apis cerana | 17 | 7 | 5 |
| Lymantria dispar | 5 | 2 | 3 |

### 6.2. Estimating climate suitability

For most plant pests, climate is likely to be the major abiotic barrier to establishment upon arrival, especially at large spatial scales (Thuiller *et al*., 2005; Araújo & Rozenfeld, 2014; Higgins & Richardson, 2014). The geographic distribution of suitable climate can be estimated using a wide variety of approaches including climate matching algorithms (e.g. CLIMATCH, CLIMEX’s climate matching algorithm; Kriticos *et al*., 2015; DAFF, 2020), environmental convex hulls and Range Bagging (e.g. Drake, 2015), correlative species distribution models (e.g. Maxent Phillips *et al*., 2006), physiological models (e.g. NicheMapper; Kearney & Porter, 2020), semi-mechanistic models (e.g. CLIMEX; Kriticos *et al*., 2015), or when data are poor, expert-derived suitability maps (e.g. Martin *et al*., 2015). Many tools exist, and there are diverse opinions on how to use them, but there remains no strong evidence of a single *best* approach for predicting an invasive species’ potential distribution (Barry *et al*., 2015; Elith, 2017). As a consequence, our work flow is agnostic as to how abiotic suitability is estimated. We strongly recommend users consult Camac *et al*. (2020) to obtain practical guidance on how to robustly estimate a species’ climatic suitability.

Here, we estimated the geographic distribution of threat-specific climate suitability using a recently proposed method known as range bagging (Drake, 2015). Range bagging is an algorithm that estimates the environmental limits of a species’ habitat by calculating convex hulls around environmental conditions at occurrence locations.

Environments that fall within the hull are defined as suitable, whereas those that fall outside are considered unsuitable. This process is then repeated using random subsets of both occurrence records as well as available environmental covariates (e.g., annual rainfall, mean annual temperature, etc.). The number of covariates included in a replicate is defined by the user. Suitability is then defined for each raster cell as the proportion of replicates that define the location as suitable. For example, a suitability score of 0.1 would indicate only 10% of the estimated convex hulls ensembled deemed that location suitable. By contrast, a score of 0.9 would indicate that 90% of estimated convex hulls deemed that location climatically suitable.

This approach has seen recent applications to invasion biology, and appears promising in the context of biosecurity. Part of the appeal is that no absences or background data are required – presence data are sufficient (Camac *et al*., 2020; Hill *et al*., 2022). This in turn removes a number of subjective decisions required in the modelling process and instead focuses solely on the data that we do have – presence locations. The method may also reduce inaccuracies that can arise from projecting to novel environmental conditions. This is because, unlike some other methods (e.g., Maxent), the method does not attempt to estimate continuous response curves, but rather defines convex hull boundaries in environmental space based on known occurrences, whereby everything within the hull is considered suitable and everything outside it is deemed unsuitable. Another major advantage of range bagging is that it can readily be used to deal with uncertainty in covariate selection. This is done by specifying low dimensionality (e.g., 2-dimensions) and allowing the algorithm to randomly select from among a suite of possible covariates – effectively resulting in an ensemble of hundreds of competing models.

Here, we used the range bagging algorithm with dimensionality set to 2 (meaning only two covariates are fitted at a time), the number of bootstrapped replicate models set to 100 and the proportion of occurrence records used per model set at 0.5. We allowed the algorithm to sample from all 19 WorldClim 2.1 (Fick & Hijmans, 2017) average historical BIOCLIM variables (i.e., BIO01 to BIO19; averages based on global weather station data between 1970 and 2000) derived from the published 5 arc-minute (approximately 10 km resolution) raster layers. We used ensembles of “simple” two-dimensional models in order to minimise biases associated with model over-fitting and collinearity, and thus, maximise the models transferability into novel environments (Camac *et al*., 2020). Work by Breiner *et al*. (2017) has found that using ensembles of small models, each with only two variables, often outperforms standard SDM methods.

We estimated the climate suitability for four of our five threats: brown marmorated stink bug, Asian honey bee, giant African snail and spongy moth. We did not estimate climate suitability for Khapra beetle, because it is a pest stored grain and foodstuff (there have been no documented cases of infestations in farms) which are mostly stored in locations not subject to ambient climatic conditions used in most climate suitability models. Moreover, the species is highly tolerant of a wide range of environmental conditions, capable of entering diapause for significant periods (6+ years) when conditions are adverse (Burges, 1963). As such, for the purposes of estimating contamination and exposure rates, we assumed that climate was not a significant barrier to entry and establishment for this species.

Occurrence records for Asian honey bee, giant African snail and spongy moth, were sourced from the Global Biodiversity Facility^4^. For brown marmorated stink bug, we used an expert compiled list of occurrence records collated by Kriticos *et al*. (2017). Prior to running the range bagging algorithms we first cleaned these occurrence records by using cleaning routines available in the R package CoordinateCleaner (Zizka *et al*., 2019). These routines are useful for flagging and removing records that are likely to have erroneous or dubious GIS coordinates. For example, in large databases such as GBIF, if a record does not have accurate GIS coordinates, data providers have sometimes used coordinates associated with a country centroid, capital city, or biodiversity institute. Generally such rough coordinates are too coarse when estimating climatic suitability for a species, which involves mapping occurrence records onto fine-resolution grided climatic data (Zizka *et al*., 2019).

Here, we removed records that:

1. had equal latitude and longitudes or within 0.5-degrees radius of coordinates 0,0^5^;
2. were within a 5 km radius of a capital city;
3. were within a 10 km radius of the centroid of a country or province;
4. were within 1-degree radius around the GBIF headquarters in Copenhagen, Denmark;
5. were within 100 m radius around known biodiversity institutions; or
6. were located in the ocean.

We also removed duplicate records and thinned occurrence records to one point per 5 arc-minutes (the spatial resolution of the WorldClim 2 climate data). Following this, we removed all occurrence records that were outside known countries of establishment as verified by threat-specific CABI distributional datasheets. This screening ensured that the remaining occurrences were most likely to be from established populations, and thus, suitable for inclusion in the range bagging analysis. The estimated current global climate suitability of the four threats can be found in Figure 6.1.

**Figure 6.1.:**
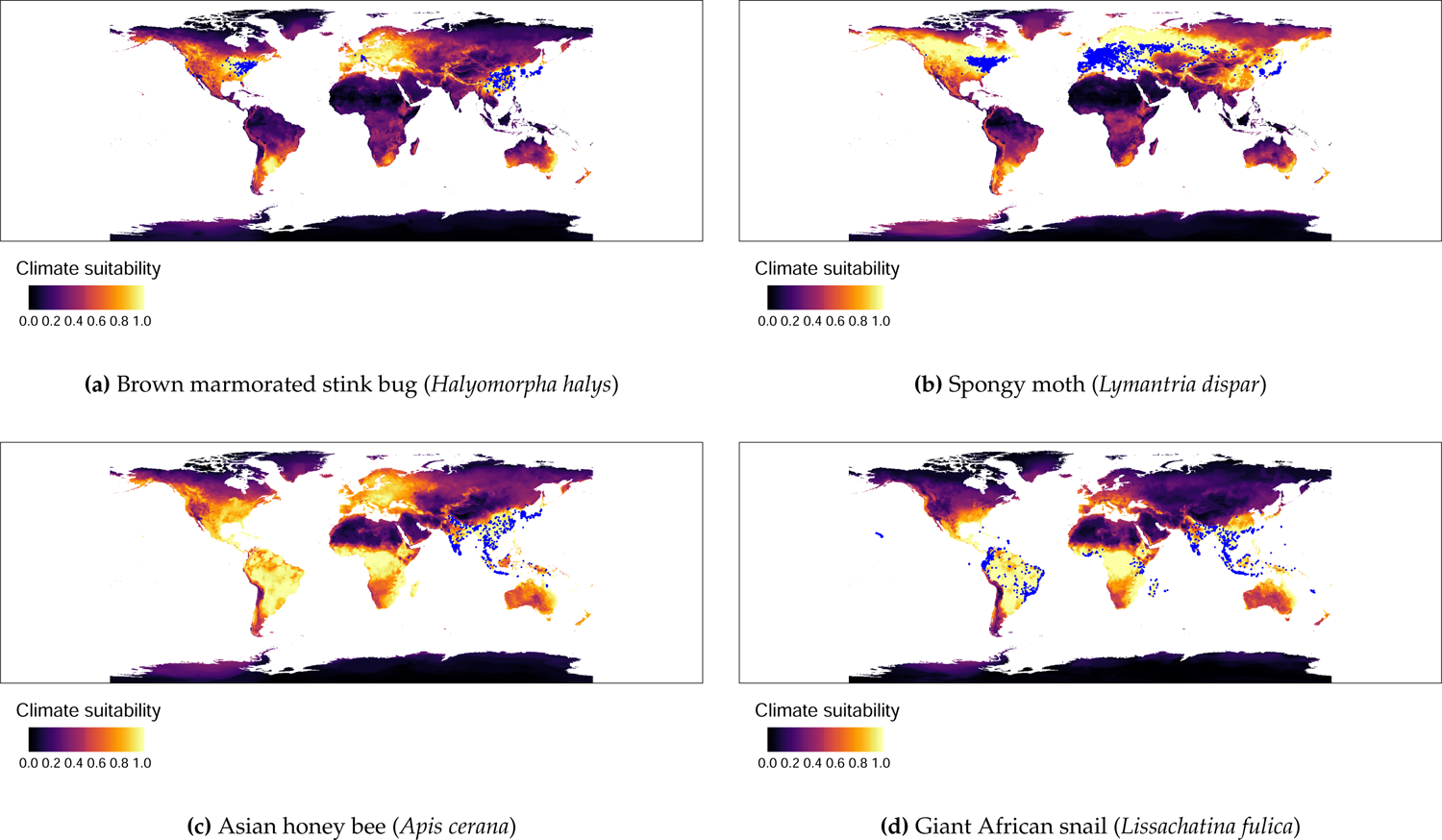
Mean global climate suitability for four exemplar hitchhiker threats for 2023. Suitability is the proportion of ensembled convex hulls that identify a location as climatically suitable across bootstrapped combinations of environmental variables. Blue dots signify known threat locations. Lighter colours (i.e., yellower) signify higher climatic suitability.

**Figure 6.2.:**
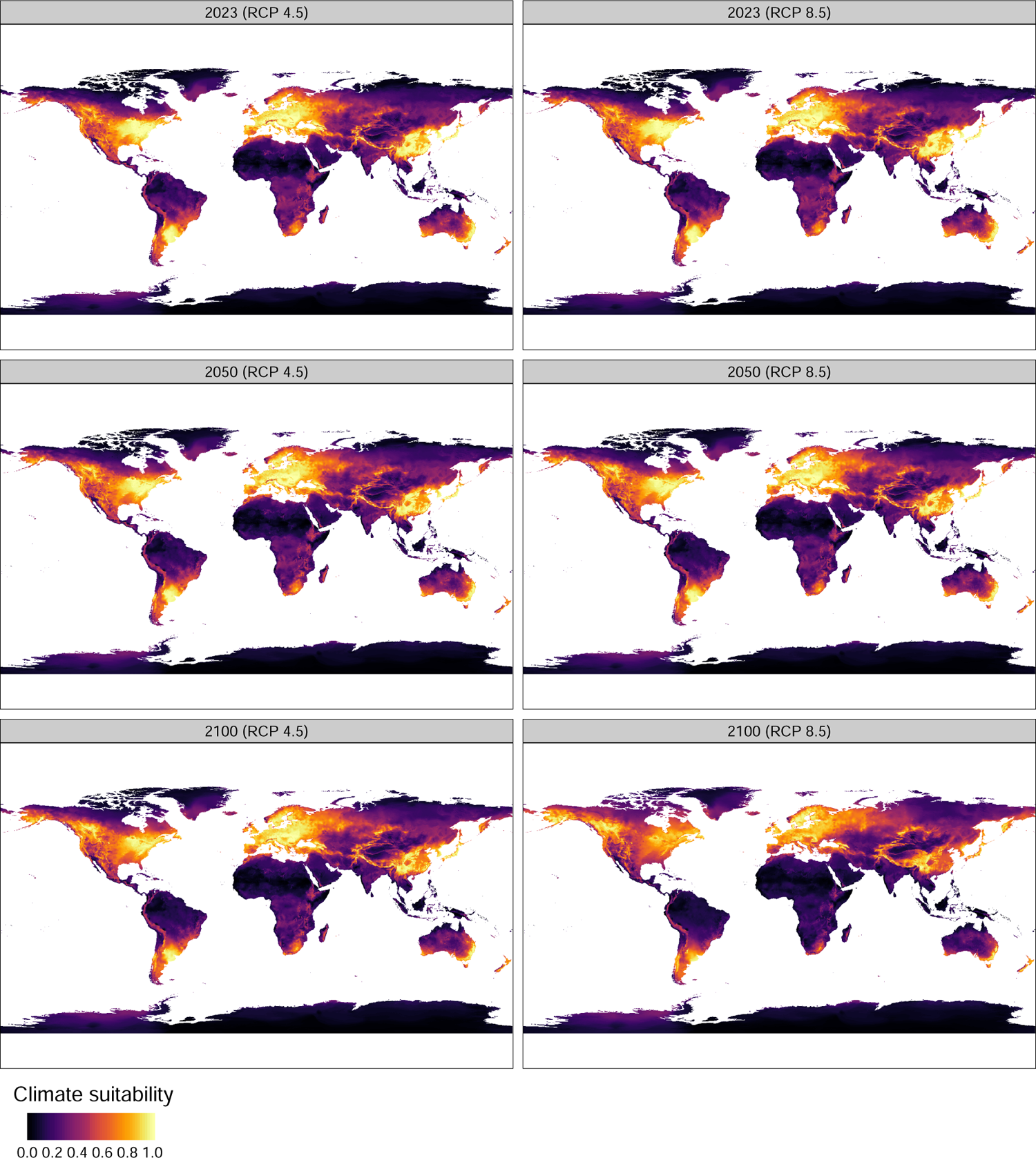
Brown marmorated stink bug (*Halyomorpha halys*) mean climate suitability at 2023, 2050 and 2100 for RCP 4.5 and RCP 8.5. Suitability is the proportion of ensembled convex hulls that identify a location as climatically suitable across bootstrapped combinations of environmental variables. Lighter colours (i.e., yellower) signify higher climatic suitability.

**Figure 6.3.:**
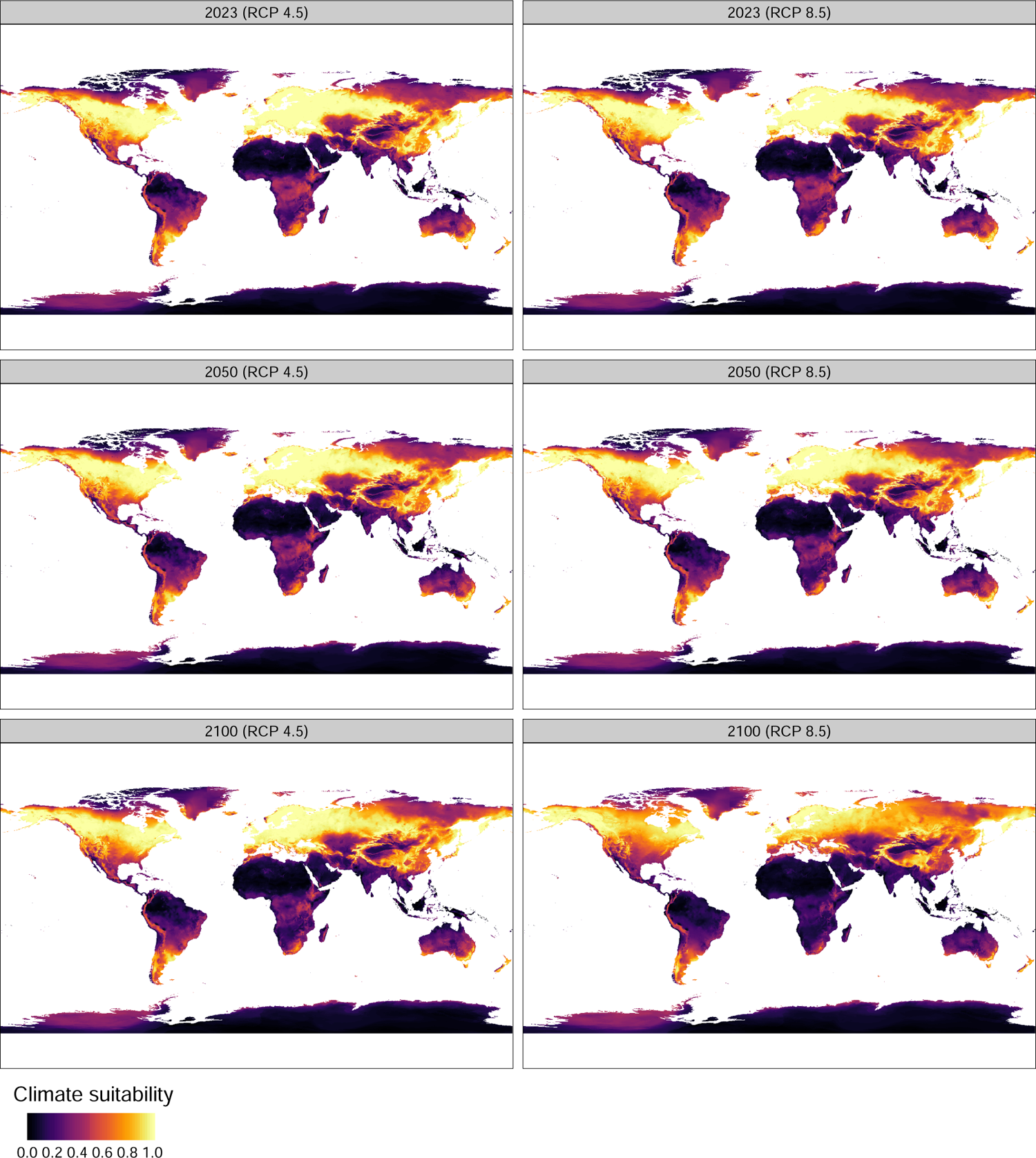
Spongy moth (*Lymantria dispar*) mean climate suitability at 2023, 2050 and 2100 for RCP 4.5 and RCP 8.5. Suitability is the proportion of ensembled convex hulls that identify a location as climatically suitable across bootstrapped combinations of environmental variables. Lighter colours (i.e., yellower) signify higher climatic suitability.

**Figure 6.4.:**
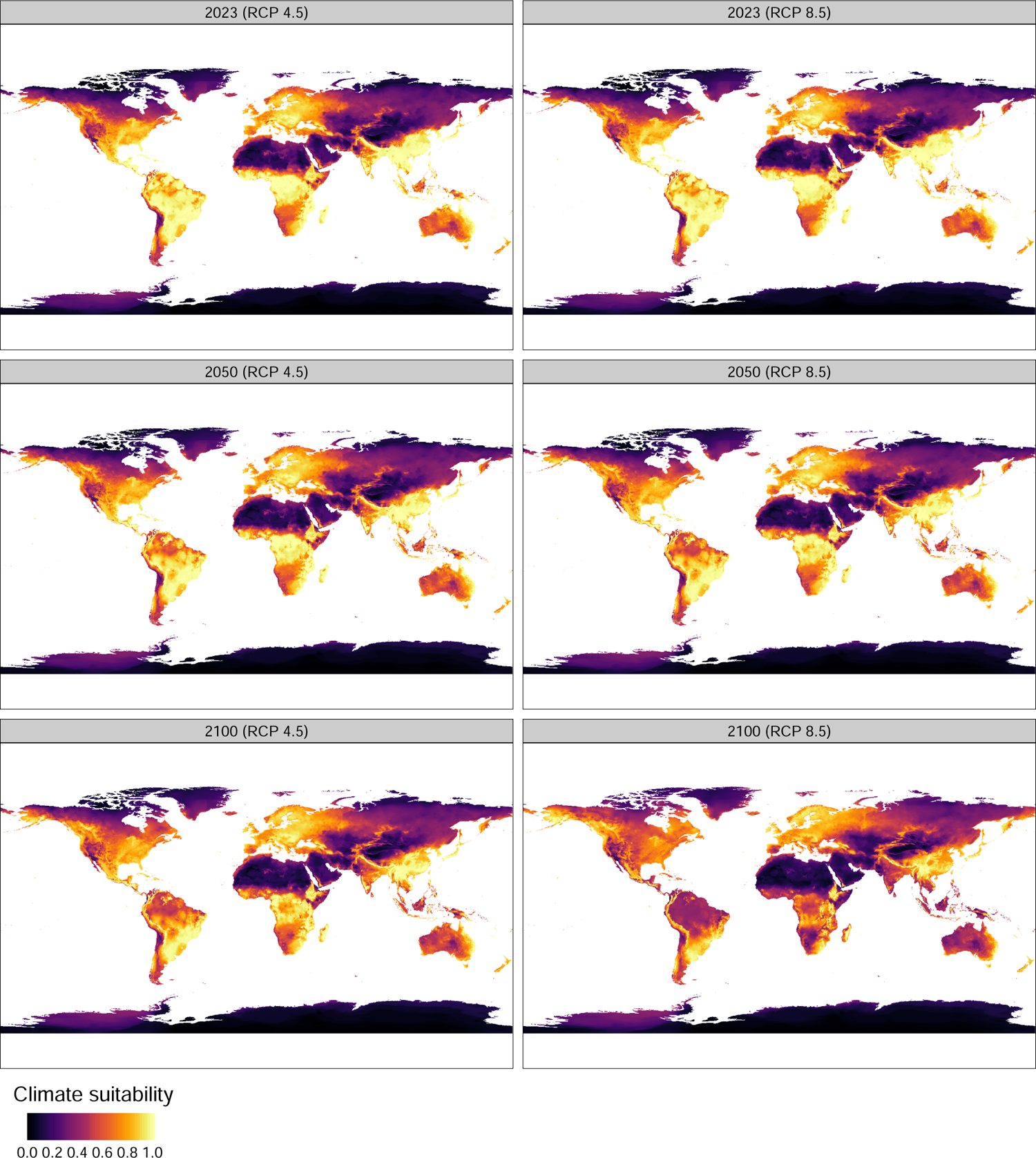
Asian honey bee (*Apis cerana*) mean climate suitability at 2023, 2050 and 2100 for RCP 4.5 and RCP 8.5. Suitability is the proportion of ensembled convex hulls that identify a location as climatically suitable across bootstrapped combinations of environmental variables. Lighter colours (i.e., yellower) signify higher climatic suitability.

**Figure 6.5.:**
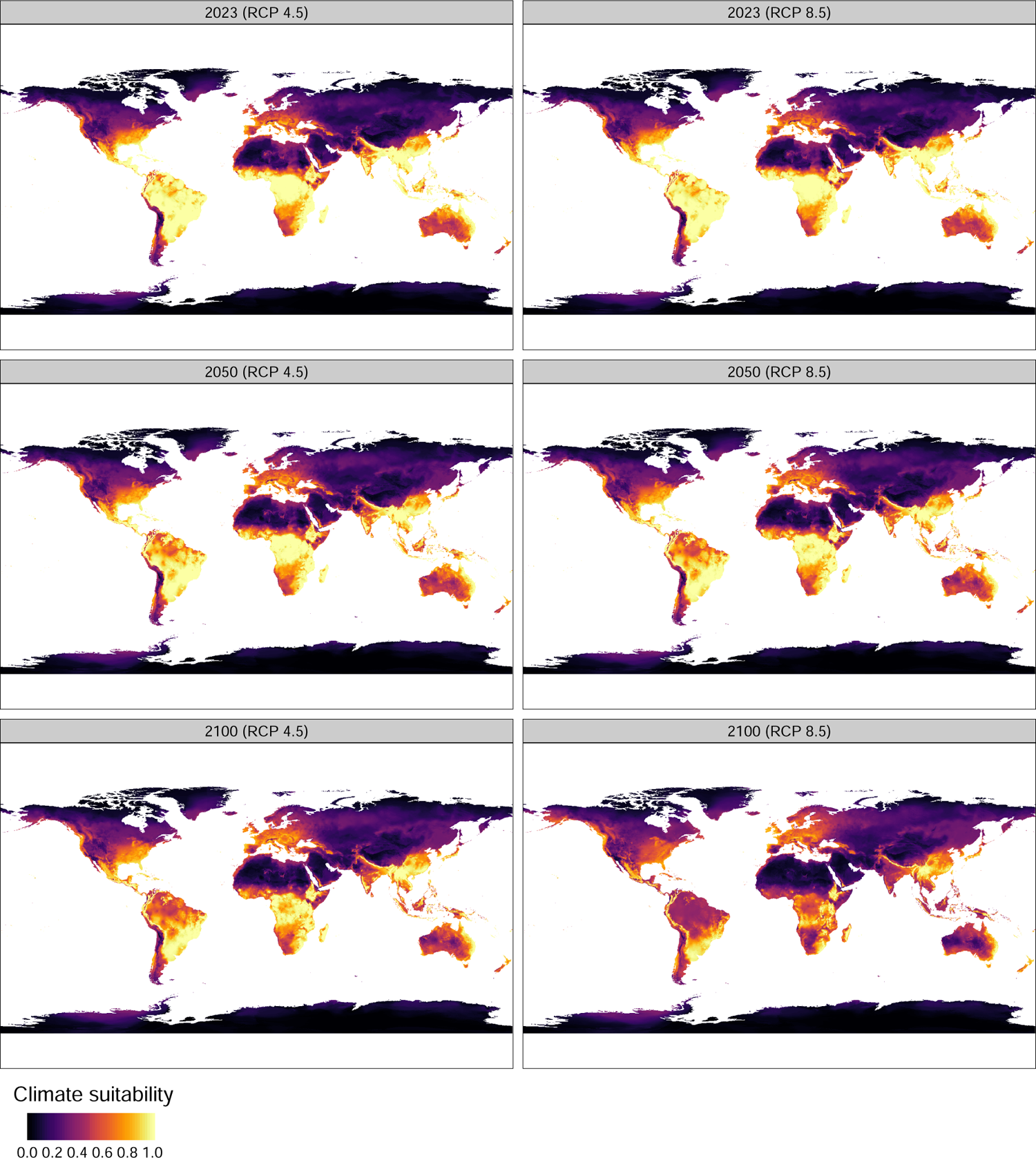
Giant African snail (*Lissachatina fulica*) mean climate suitability at 2023, 2050 and 2100 for RCP 4.5 and RCP 8.5. Suitability is the proportion of ensembled convex hulls that identify a location as climatically suitable across bootstrapped combinations of environmental variables. Lighter colours (i.e., yellower) signify higher climatic suitability.

#### Projecting future climate suitability

To predict annual changes in climate suitability for each threat under different climate change scenarios, we used downscaled (5 arc-minute) and bias-corrected BIOCLIM variable projections derived from 23 CMPIP6 global climate models (GCMs) and Shared Socio-economic Pathways SSP2-45 and SSP5-85, provided by WorldClim v2.1.^6^ Each GCM contained projected averaged estimates of the 19 standard BIOCLIM variables for four 20-year future time horizons (2021-2040, 2041-2060, 2061-2080, 2081-2100). We used the midpoint years of the historical and future projected time horizons (i.e., 1985, 2030, 2050, 2070, 2090) as a basis for piecewise linear interpolation to derive GCM- and SSP-specific annual multiband rasters of the 19 BIOCLIM variables for each year from 2014 to 2100.

Using the fitted range bag models described in section 6.2, coupled with GCM and SSP specific annualized BIOCLIM variables, we predicted climate suitability for each year from 2014 to 2100. This resulted in 3956 (i.e., 23 GCM *×* 86 year *×* 2 SSPs), 5 arc-minute rasters of predicted climate suitability for each threat. These annual GCM-specific rasters were then ensembled across GCMs for each year to provide annual mean estimates of climate suitability (see Figures 6.2 to 6.5) for both SSP2-45 and SSP5-85 climate scenarios ^7^. These annual ensembled rasters were then reprojected to Moillwede equal area projection such that each cell was approximately 10 km^2^. This was done to reduce latitudinal biases associated with unequal cell areas when quantifying the annual proportion of suitable climate (i.e., mean climate suitability) in a country or GTAP region.

### 6.3. Contamination model

The original model, developed in Camac *et al*. (2021a), estimated the expected number of contamination events hitting a country’s border using a Bayesian Generalised Additive Mixed Model (GAMM). In this model, the observed number of interceptions, *y_ijt_*, originating from each infected country, *i*, commodity type, *j*, and year, *t*, combination was modeled as a function of the expected value of inspected imported goods (in USD) with random intercept terms for country of origin, commodity type, and year. Dependent on the level of over-dispersion present in the interception data, these counts were modeled as a random realisation from either a Poisson or negative binomial distribution.

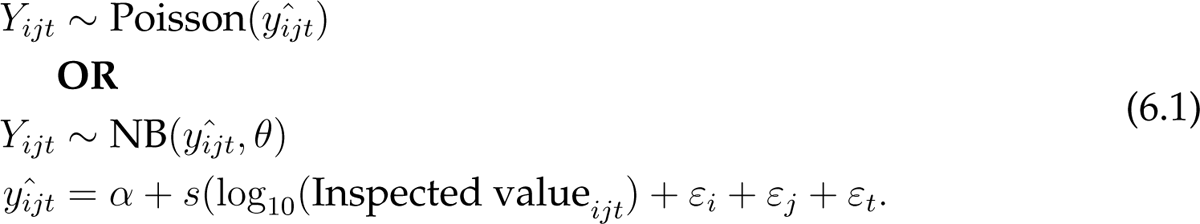

#### New model formulation

In this project we make several refinements to the original model. First, to better encapsulate the underlying sampling process, we have shifted to a binomial regression model. This has two distinct advantages: 1) It more appropriately accounts for the sampling uncertainty used to derive contamination counts – that is, individual consignments are inspected and the outcome is binary (either contaminated or not); and 2) The model estimates a more useful metric – the likelihood an imported line from a given infected country is contaminated. In making this change, we’ve shifted away from using the expected import value of inspected goods (i.e. Inspected value*_ijt_*) as a non-linear smoothing term in the model.

Instead, we now model the likelihood that an imported line (*p_ijt_*) is contaminated with a threat as a function of infected GTAP exporter region (*ε_i_*) (Table 5.1), the type of commodity (*ε_j_*), and year (*ε_t_*) as random intercept terms in a aggregated binomial Generalised Linear Mixed Model (GLMM) with logit link. Furthermore, we included a fixed effect for the mean estimated climate suitability climsuit*_it_* of a GTAP region in a given year, *t*. For GTAP regions that are a aggregation of multiple countries (e.g., GTAP region (`xef`)), we estimated climsuit*_it_*as the sum of climate suitability in countries of known establishment, divided by the total number of raster cells in the GTAP region. This better represented the current potential distribution of the threat. By incorporating climate suitability effect, we can assess how contamination rates among infested regions vary as a function of the average climate suitability, and thus, how contamination rates may change under future climatic conditions as a species’ potential range may expand or contract.

As such our new model formulation is as follows:

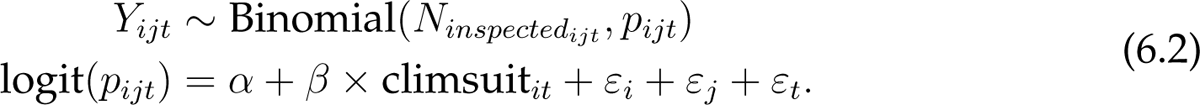

Here, our response variable, *Y_ijt_*, is the observed number of contaminated lines exported from GTAP region, *i*, of commodity type, *j* in year *t*. *N_inspectedijt_* is the associated number of lines inspected at the border. Like the original model, this formulation includes random intercept effects for country of origin, *ε_i_*, commodity type, *ε_j_* and year *ε_t_*. Random effects were included for five critical reasons: 1) they provide a convenient method for appropriately accounting for the non-independent structure of observations within years, country of origin, and tariff codes; 2) they allow us to estimate group-level effects without substantially increasing the degrees of freedom; 3) they allow estimation of group-levels with few observations through a method referred to as “partial pooling” (Gelman & Hill, 2007) whereby low observation groups are estimated closer to the mean across groups, but are also more uncertain relative to groups with many observations; 4) they provide a means for estimating the group-level variation that is otherwise not accounted for by fixed effect terms; and most importantly 5) they provide a means for making predictions to new group-levels not included in model fitting (e.g. high risk commodities not imported into Australia) (Gelman & Hill, 2007).

Models were fit using the R package brms (Bürkner, 2021) with a weakly informative Gaussian priors specified on the intercept and coefficient terms (i.e., *N* (0, 10)) and exponential priors on the variance terms (i.e., Exp(1)). We ran the models using 4 chains, sampling 2000 iterations for each. Chain convergence was assessed using the Brooks-Gelman-Rubin convergence diagnostic on the last 1000 iterations of each chain (the first 1000 being used as burn-in) (Brooks & Gelman, 1998). Posterior inferences were made using the last 1000 iterations of each chain (i.e., 4000 samples in total).

Based on realised border screening effort, we used posterior predictive checks to ensure the models were able to accurately predict observed numbers of contaminated lines entering Australia in aggregate as well as partitioned by infected exporter, commodity and year. Across all threat models, the posterior predictions encompased the observed values (For more details see: Appendices A to E).

#### Forecasting annual propagule pressure

The primary focus of this project was to forecast international trade flows under different climate scenarios and examine how these changes may impact propagule pressure and establishment exposure of hitch-hiking threats hitting Australia and its trading partners borders. The GTAP model provides users with proportional changes in trade flows, which can be converted to absolute changes in import/export value relative to a baseline. In this project, GTAP outputs were converted to absolute USD values using 2014 as the baseline year. Our new contamination model estimates the likelihood an imported line is contaminated with a threat. As such, to estimate propagule pressure (i.e., the number of infected lines hitting a countries border) we need to convert these import estimates of trade value to expected numbers of imported lines.

We approximated this by first extracting the Australian GTAP import values estimates of each commodity by infected exporter by year combination for 2014 to 2021 (i.e., Import value*_ijt_*). We then divided these values by their associated observed number of lines imported into Australia over the same period (i.e., *N_obsijt_*). This resulted in an approximate estimate of the value of a single imported line in each year of a particular commodity type coming from a particular exporter. We then calculated the median line values, line value*_ij_*, across years for each exporter by commodity combination. These median import values were then used to convert GTAP import values to expected annual numbers of imported lines (rounded up to the nearest integer), *N_ijkt_*, exported from infected region, *i*, of commodity type, *j*, into GTAP region, *k* for each year, *t*. These were then used multiplied by 1000 samples of the posterior estimate of *p_ijt_* to derive the expected number of contaminated lines associated with each *ijkt* combination. To predict contamination rates of unobserved random grouping variables not present within the fitted data (i.e. years *>* 2022 and commodity types from infected countries that do not export to Australia) we took 1000 random posterior draws, such that each posterior sample could be drawn from a different group level. This sampling meant that the posterior draws represented the variation across all existing levels, and thus, appropriately reflected the uncertainty in predicting unobserved groups.

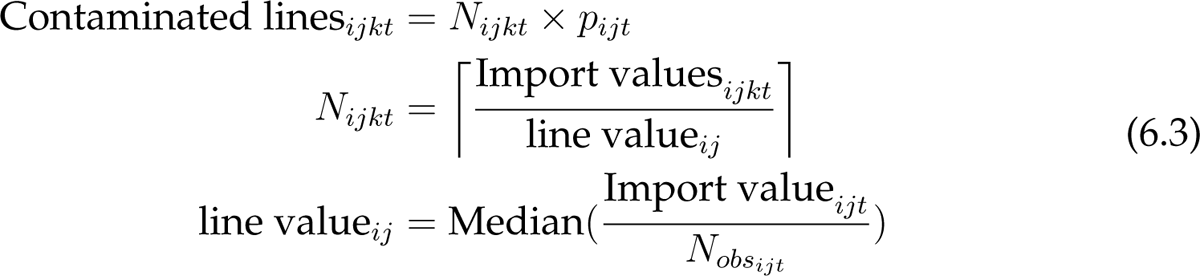

The total propagule pressure hitting a given GTAP region’s borders *k* in year *t*, can then be calculated by summing the expected contaminated lines across infected GTAP regions *i* and commodity type, *j*:

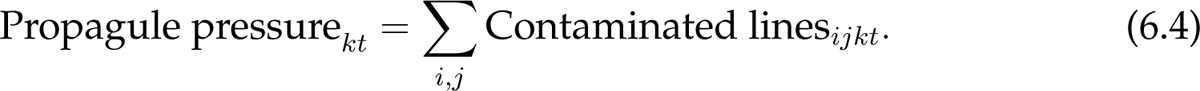

### 6.4. Estimating establishment exposure given propagule pressure

For threats where potential distributions are likely to be influenced by ambient climate conditions (i.e. all exemplar threats except for Khapra beetle), we estimate establishment exposure as the number of propagules arriving at a given location multiplied by the likelihood the location is climatically suitable for establishment. We do this by first assuming imported goods will most likely be unloaded in regions of high human population density (i.e., where demand is expected to be greatest). We also assume that these locations will be where a hitchhiking threat has the greatest opportunity to escape and potentially establish. There is evidence to support the use of such an assumption, with studies commonly finding a strong correlation between human population density and first detections of exotic threats even when correcting for potential survey bias (Dodd *et al*., 2015).

The original threat exposure model used a contemporary raster of human population (i.e., population counts in 2020) for distributing country-level establishments of propagule pressure. As our focus was to project changes in establishment exposure over time and under different climate scenarios, we replaced this the contemporary human population layer used in the original model with global annual raster projections of human population counts derived from work recently published by Olén & Lehsten (2022). Olén & Lehsten (2022) generated global 30 arc-second (approximately 1km^2^) rasterized population projections for 6 combinations of SSPs and RCPs, with annual predictions spanning from 2010 to 2100. We retrieved published predictions from Olén & Lehsten (2022) for both SSP2 RCP 4.5 and SSP5 RCP 8.5. These annual rasters were aggregated (i.e. summed) to 5 arc-minutes and then reprojected to Mollweide equal area projection so that they were in alignment with the climatic suitability layers.

Using these annual human count rasters, we distributed the mean (*±* 95% credible intervals) propagule pressure within each country, *k*, as a function of the proportion of human population present in each raster cell *l* for a given year, *t*:

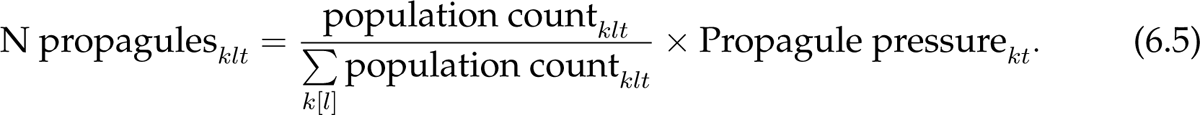

We then accounted for the climate suitability at those locations by weighting the expected number of propagules, N propagules*_klt_*, at each country-specific grid cell, *k*[*l*] in each year *t* by the annual estimated climatic suitability of that location:

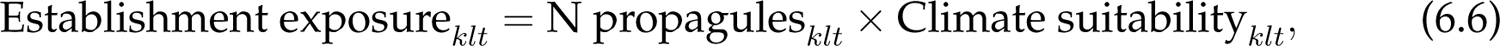

where climate suitability is bounded between zero and 1 (i.e. the output from range bagging).

Finally, we estimated the mean (*±* 95% credible) Establishment exposure*_k_ t*, for each GTAP region, *k* and year *t* by summing the grid cell estimates of establishment exposure, *N*_Establishment exposure_, for each region and year combination:

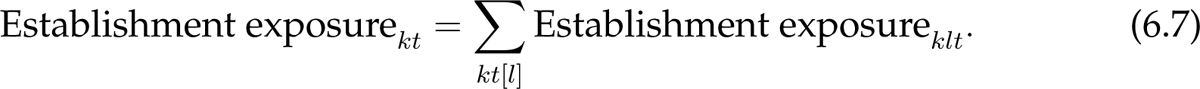

### 6.5. Fundamental model assumptions

The contamination model makes several pragmatic assumptions based on constraints associated with the available data. Specifically, the model assumes that:

1. **The border sampling within each country and commodity combination is random and independent.** Non-random sampling may bias contamination estimates by either over or under-estimation of contamination likelihoods. The validity of such an assumption is likely to vary significantly among commodity types with some being easier to undertake random and independent sampling than others.
2. **The contamination rates observed in Australia are representative of what is experienced by other countries.** This may not always be true, as other countries may impose pre-border interventions or none at all. Given Australia is a world-leader in biosecurity risk mitigation, implementing a range of pre-border interventions (including for the exemplars used in this study), we believe the derived contamination likelihoods are likely to be at the conservative end of the spectrum (i.e. contamination rates might be higher in locations with fewer biosecurity pre-border controls).
3. **Border screening detection rates are similar across commodity sectors.** In reality, this is unlikely to be true (Garrard *et al*., 2008; Wintle *et al*., 2012) as detection rates are known to vary substantially among species (Garrard *et al*., 2012; Martin, 2017), commodity types, points of entry, and the size and arrangement of consignments (which were not recorded). Imperfect detection rates (e.g. a pest is harder to find on certain commodity types) may result in underestimation of the true contamination rates for those commodity types. However, due to the lack of leakage survey data for each commodity type, estimating such rates is currently not practical without making additional model assumptions or undertaking expert elicitation surveys (Hemming *et al*., 2018; Courtney Jones *et al*., 2023)^8^.
4. **Country by commodity contamination rates only vary overtime as a function of changing climate suitability (*if included*).** Again this is unlikely to be true as source populations increase or contract for a variety of reasons. For example, countries with new incursions may have relatively low contamination rates as the area infested is low. However, as time goes by, area of infestation may increase, which may translate into higher commodity exposure to the pest, and thus, higher contamination rates. However, to account for such a process requires an indepth understanding of the area of infestation in each invaded country and how this may change over time – acknowledging that different countries are likely to have different patterns of spread and differing capabilities in managing and containing outbreaks.
5. **All contamination events have equal establishment viability.** This is unlikely to be true as the viability of any contamination event will depend on the level of contamination (i.e. number of individuals) and the transport survivability, factors that will vary substantially among exporting regions and commodity types. Currently we do not have the necessary data to approximate these viability rates.
6. **Estimates of numbers of imported lines are accurate.** Unfortunately, statistics on the number of commodity lines imported into different countries is generally not reported. Instead, most trading countries provide databases such as the United Nations COMTRADE database with estimates of the annual value of commodity-specific imports obtained from different exporters. Here, we used Australian data to approximate the median value of a imported line for each country by commodity combination. The assumption here is that these estimated median values are representative of the the true line values imported elsewhere.
7. **Estimates of propagule pressure assume no spread to unoccupied regions.** New incursions are inherently difficult to predict as they are stochastic and dependent on a wide range of factors (e.g. region biosecurity preparedness, pathway viability rates, within-region secondary movements) that ultimately influence the likelihood of establishment and subsequent spread. As such, the model reported here makes the pragmatic decision of only estimating contamination rates and subsequent outgoing propagule pressure for regions with known established threat populations. As a consequence, future propagule pressure may be underestimated if new incursions occur in unoccupied regions. If such an event occurs, one would need to rerun the model with an updated list of invaded regions. Note, a potential extension to this model is to simulate global spread (See Chapter 12).

### 6.6. Model outputs: User friendly interactive maps of pest pressure and exposure for all 70 regions

While the outputs reported in here are focused on Australia (see following chapters), the contamination model is capable of estimating trends in propagule pressure and establishment exposure for all 70 regions. To facilitate interrogation of model outputs, we have developed leaflet interactive maps for each of the five threats (Figure 6.6). Included in these maps^9^ are information about temporal trends in pest pressure, establishment exposure (if threat is influenced by climate suitability) and proportion of risk attributable to each infected exporters and commodity sector.

**Figure 6.6.:**
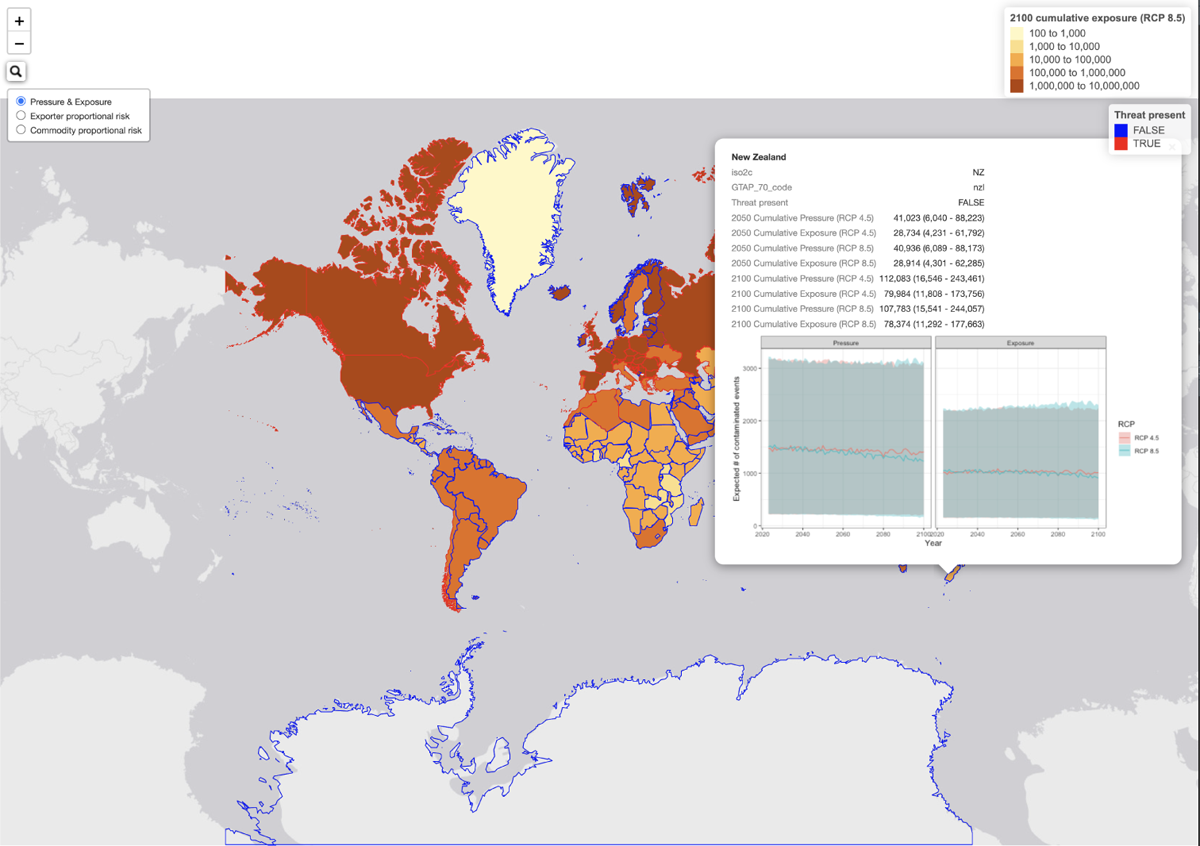
Example screenshot of interactive map showing predicted temporal trends in BMSB propagule pressure and establishment exposure in New Zealand.

## 7. Overwintering pest: Brown marmorated stink bug (*Halyomorpha halys*)

### 7.1. Likelihoods of contamination

The brown marmorated stink bug contamination model revealed that regions with higher mean climate suitability (i.e., climsuit*_it_*) tended to have higher likelihoods of contamination (confidence in direction > 70%; Figure 7.1a). The model also revealed that, even after accounting for exporter climate suitability (i.e. fixed effect term), there was considerable variation within and between random grouping variables (Figure 7.1b). Specifically, contamination rates were most variable among commodity sectors (Commodity SD = 1.89), followed by exporter region (Region SD = 1.48) and then year (Year SD = 1.15).

**Figure 7.1.:**
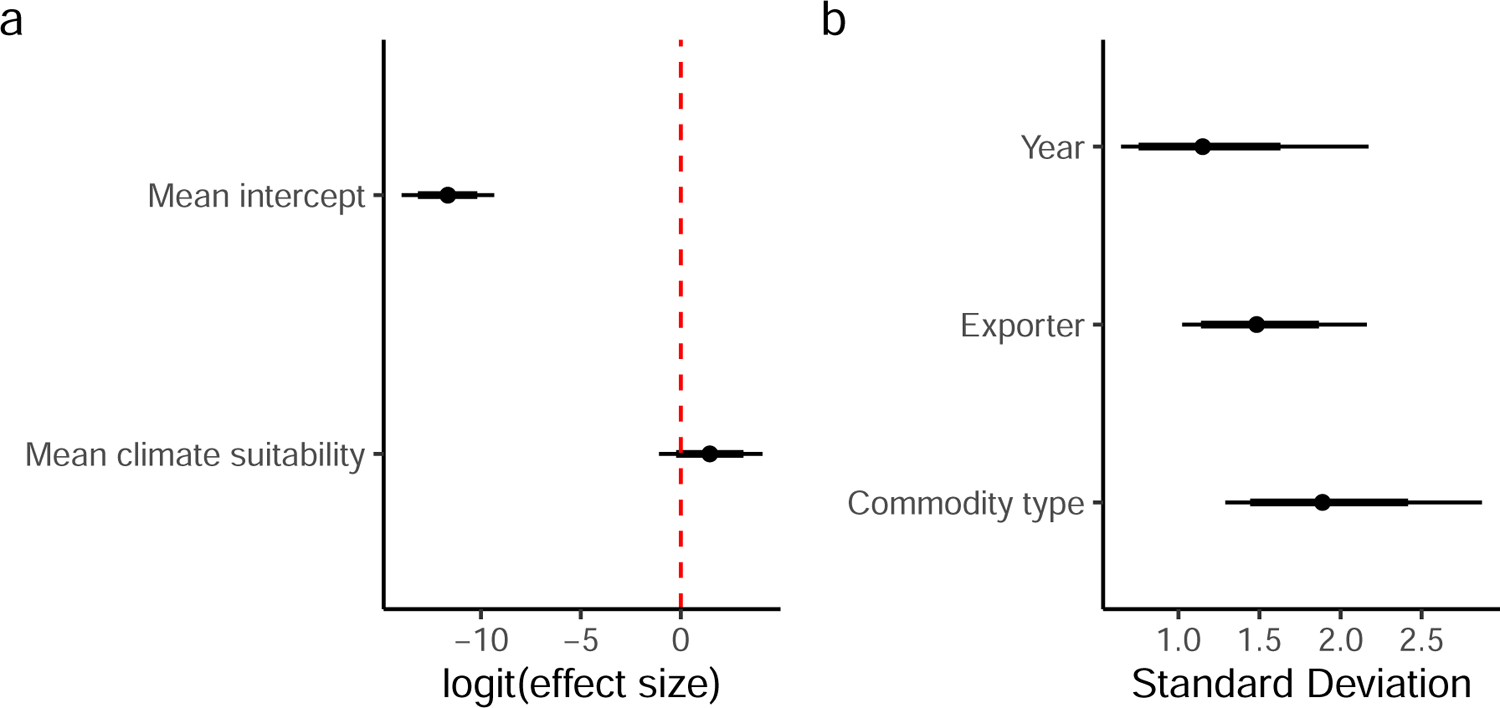
Brown marmorated stink bug (*Halyomorpha halys*) coefficient and group variance plots. a) Model coefficients; b) Random effect standard deviations (i.e. measure of variability within each grouping variable). Error bars reflect 95% (thin) and 80% (thick) credible intervals.

Examination of baseline likelihoods for infected exporters (Figure 7.2a) revealed that Italy (`ita`) had on average the highest contamination rates. This was followed by the United States of America (usa) and the GTAP aggregation “rest of Europe” (xer). By contrast, Taiwan (twn), Korea (kor), France (fra), and Portugal (prt) had some of the lowest infected region contamination rates among infected regions.

Baseline likelihoods for commodity sector (Figure 7.2b) also showed that BMSB contamination rates were highest in petroleum and coal products (p_c), mineral products (nmm), motor vehicles and associated parts (mvo), electronic equipment (eqm), and chemical, rubber and plastic products (crp). By contrast, sugar cane/beet (c_b), processed rice (pcr), food products (ofd), and beverages and tobacco products (b_t) exhibited some of the lowest estimated contamination rates among modelled commodity sectors.

**Figure 7.2.:**
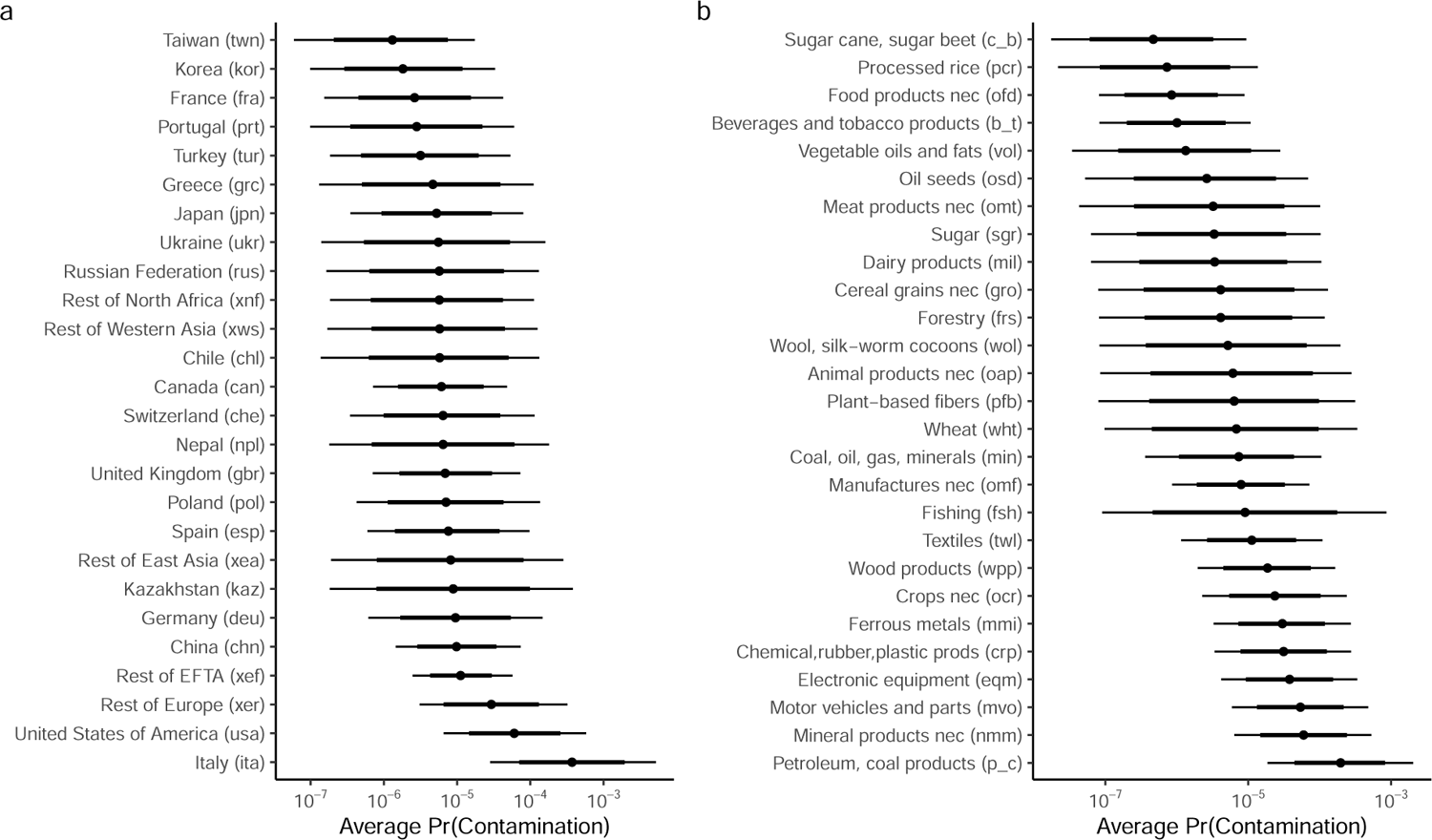
Brown marmorated stink bug (*Halyomorpha halys*) baseline contamination likelihoods associated with (a) infected exporter; and (b) commodity type. Error bars reflect 95% (thin) and 80% (thick) credible invervals. Note: baseline likelihoods are back-transformed (anti-logit) intercept terms in the model and that do not account for exporter mean climate suitability.

BMSB contamination rates also varied substantially between 2014 and 2021 (Figure 7.3). Specifically, average contamination rates prior to 2017 tended to be lower relative to post-2017. Such a temporal pattern may reflect a true increase in contaminated imports arriving at the Australian border in recent years. However, it may also be an artefact of changed border screening practices that have resulted in higher border detection rates for this species.

**Figure 7.3.:**
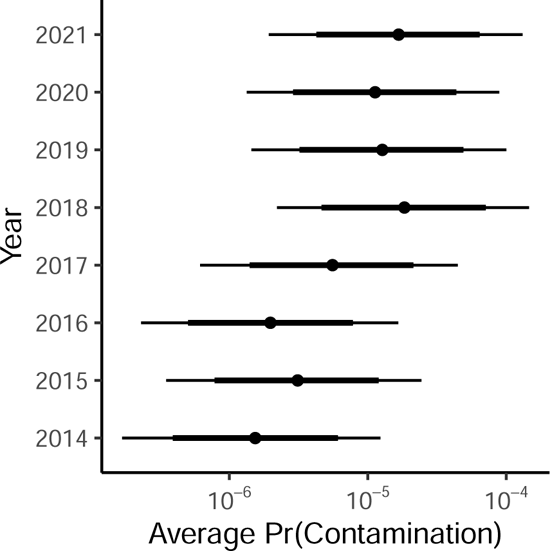
BMSB annual baseline contamination likelihoods. Error bars reflect 95% (thin) and 80% (thick) credible invervals. Note: baseline likelihoods are back-transformed (anti-logit) intercept terms in the model and that do not account for exporter mean climate suitability.

Imports from Italy (`ita`) accounted for 15 of the top 20 estimated contamination likelihoods for 2023 (Table 7.1). In particular, imports of petroleum and coal products (p_c) from Italy contained the highest estimated contamination likelihood with a 3.1% chance a single imported line was contaminated. This was followed by imports of mineral products (nmm; 0.91%), motor vehicles and associated parts (mvo; 0.83%), electronic equipment (eqm; 0.59%), and chemical, rubber and plastic products (crp; 0.48%). Other exporters that fell within the top 20 included the United States of America (usa) and Germany (deu), with imports of petroleum and coal products (p_c) containing an estimated BMSB contamination likelihood of 0.42% and 0.092% for each exporter, respectively.

**Table 7.1.:**
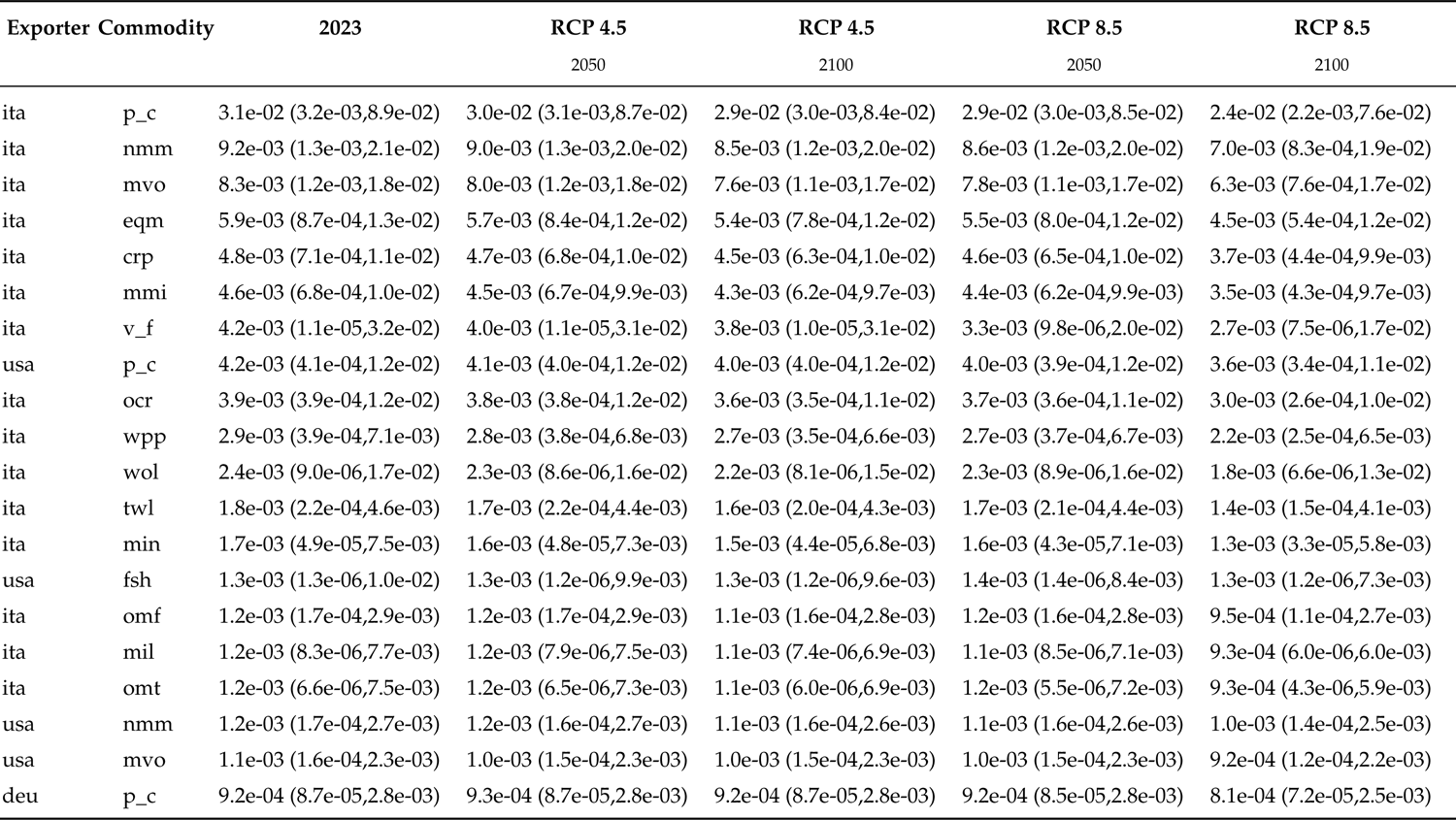
Brown marmorated stink bug top 20 commodity sector by region contamination likelihoods for commodities with 100 or more imported lines into Australia and how they are predicted to change under RCP 4.5 and RCP 8.5 by 2050 and 2100. Numbers in brackets signify 95% credible limits.

As the model accounted for predicted changes in BMSB climate suitability on infected exporter contamination rates, we also found that over time contamination rates would marginally decrease in regions whose BMSB climate suitability is expected to decline. For example, average contamination rates associated with p_c from Italy are expected to decrease from approximately 3.1% in 2023 to 2.9% by 2100 under RCP 4.5. The decline is even greater under the RCP 8.5 scenario, with contamination rates predicted to decline to 2.4% by 2100. These declines are driven by a decrease in the average BMSB climate suitability in Italy, which in 2023 is estimated to be 0.86 (0.84-0.88) and by 2100 is estimated to be 0.8 (0.74-0.86) under RCP 4.5 and 0.65 (0.57-0.75) under RCP 8.5, respectively.

Declines in contamination rates over time were exhibited by most infected regions. This is because mean climate suitability in most infected exporter regions are predicted to decline (Table 7.2). However, exceptions do exist. Regions such as Canada (can), Chile (chl), United Kingdom (gbr), and Russia (rus) are all predicted to increase in climate suitability between now and 2100. This increase in average climate suitability is expected to have the consequence of increasing contamination likelihoods as the threat’s distribution in these regions expands.Ultimately manifesting in higher risk of entry and establishment in importing countries such as Australia.

**Table 7.2.:**
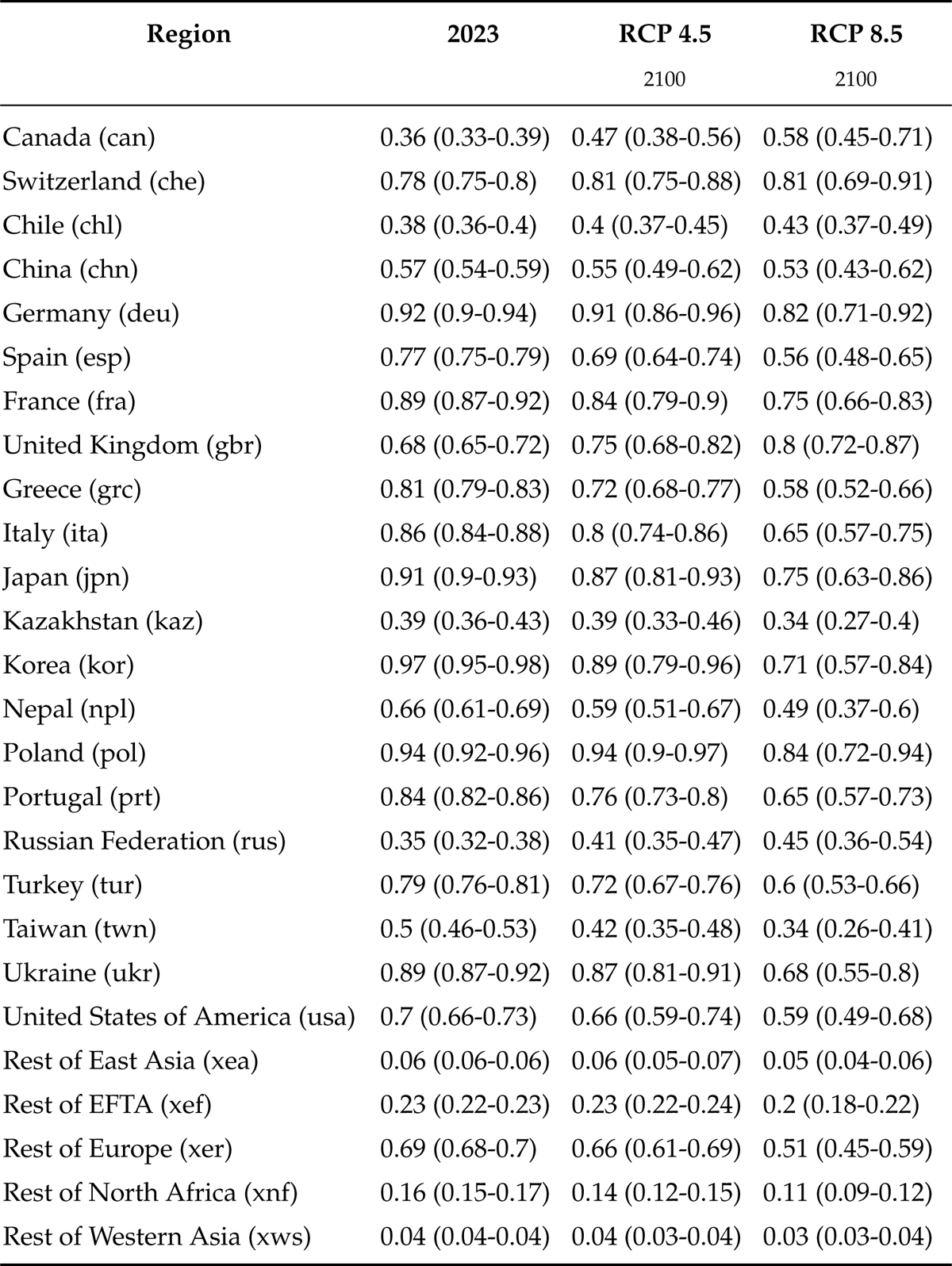
Brown marmorated stink bug mean climate suitability for each infected exporter region in 2023 and 2100 for RCPs 4.5 and 8.5. Mean estimates are based on the ensembled annual climate suitabilities derived from 23 GCMs. Numbers in parentheses are the 10th and 90th percentiles derived from ensembles annual models.

### 7.2. Australia: BMSB propagule pressure and establishment exposure

Annual brown marmorated stink bug propagule pressure arriving at Australian borders is expected to marginally decline from 8779 (95CI: 1298-19035)^1^ in 2023 to 8338 (95CI: 1234-18427) and 7181 (95CI: 974-18123) by 2100 for RCP 4.5 and RCP 8.5, respectively (Figure 7.4).

**Figure 7.4.:**
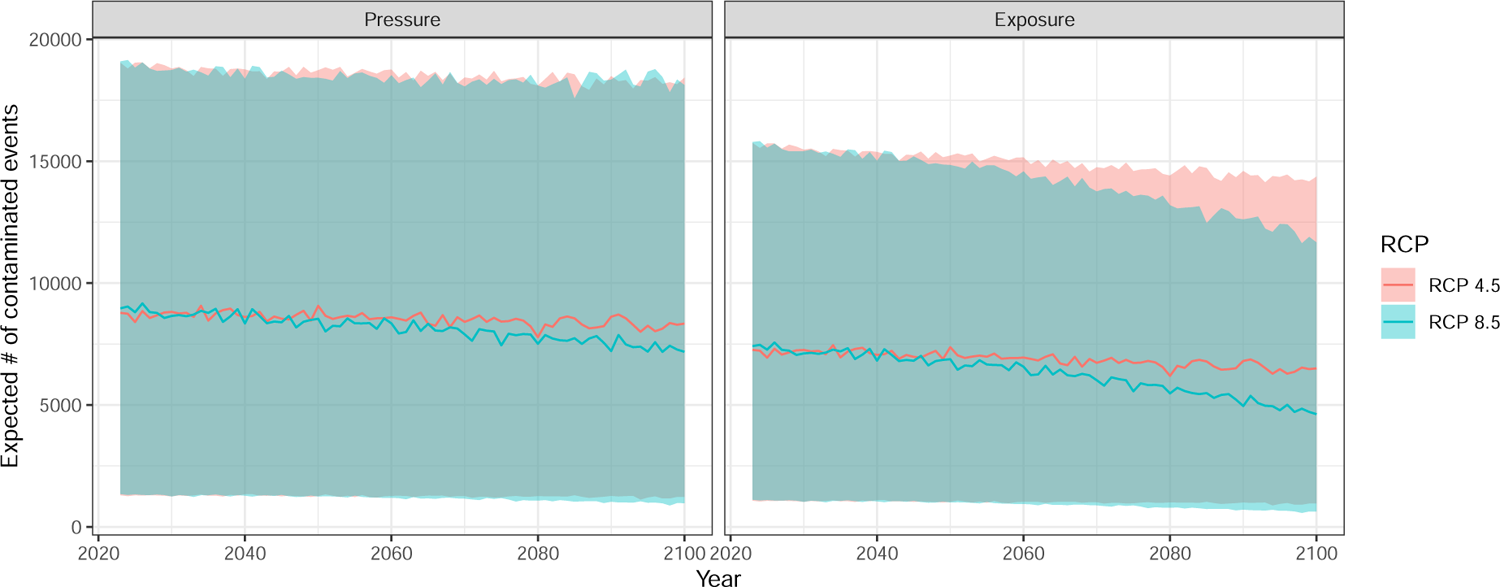
Annual mean (95% CI) brown marmorated stink bug (*Halyomorpha halys*) propagule pressure (left) and establishment exposure (right) hitting Australia under RCP 4.5 and 8.5. Note: Predictions include stochastic uncertainty associated with future years and for unobserved commodity and infected exporter combinations.

Estimated establishment exposure in Australia (i.e., propagule pressure conditioned on climate suitability in the importing region), is predicted to decline under RCP 8.5 but remain relatively stable under RCP 4.5. This is because under RCP 8.5 there is a more significant decline in BMSB suitable climatic conditions at locations where imported goods are expected to be unloaded (i.e., regions of highest human population density; Figure 6.2). As a consequence, the expected number of contaminated lines arriving at a climatically suitable locations within Australia will be lower.

### 7.3. Australia: BMSB risk attributable to different exporters & commodity sectors

Using the model outputs we can also examine the proportion of total propagule pressure attributable to each infected exporter (Figure 7.5) or commodity sector (Figure 7.6), and how these proportions may change over time under different climate scenarios. We found that in 2023, Italy (`ita`) is expected to account for 61% (58-63) of the total BMSB propagule pressure hitting Australian borders. However, by 2100, this proportion marginally decreases to 60% (56-63) and 57% (49-65) for RCP 4.5 and 8.5, respectively. By contrast, proportional risk attributable to regions such as the United states (usa), China (chn) and the United Kingdom (gbr) are all expected to increase, particularly under the RCP 8.5 scenario (Figure 7.5). These proportional changes are small and mostly attributable to changes in climate suitability in the exporting regions (i.e., increased climate suitability in United Kingdom, but declines in Italy) but also small changes in the relative contributions in imports into Australia from different exporting regions (e.g. decreases in imports of many commodities from China and associated increases from the United Kingdom and the USA; Figure 5.2). It is important to note, that the current CGE trade model incorporates only a limited number of climate related damage functions that have only small impacts on manufactured goods – the primary sector that this pest, and others included in this report, are most likely to hitchhike on. As such, magnitude of changes are should be interpretated as likely being underestimated, with a range of damages that could impact manufacturing (e.g. coastal inundation, physical capital depreciation) largely unaccounted for at present.

While there was some marginal changes in BMSB propagule pressure attributable to different exporters, proportional risk attributable to different commodity sectors was estimated to remain relatively stable under both RCP scenarios (Figure 7.6). Imports of electronic equipment (eqm) was estimated to be the largest contributor to BMSB propagule pressure, accounting for approximately 38% (35-41) of the total number of predicted BMSB contaminated lines entering Australia. This was followed by ferrous metals (mmi; 13% (11-14)), textiles (twl; 10% (6-14)), and manufactures nec (omf; 4% (3- 6)). It is important to note that while petroleum and coal products (p_c) were found to have relatively high contamination rates, its overall contribution to BMSB propagule pressure was small, contributing less than 1% – highlighting that Australia imports relatively few lines of this commodity from infected exporters.

**Figure 7.5.:**
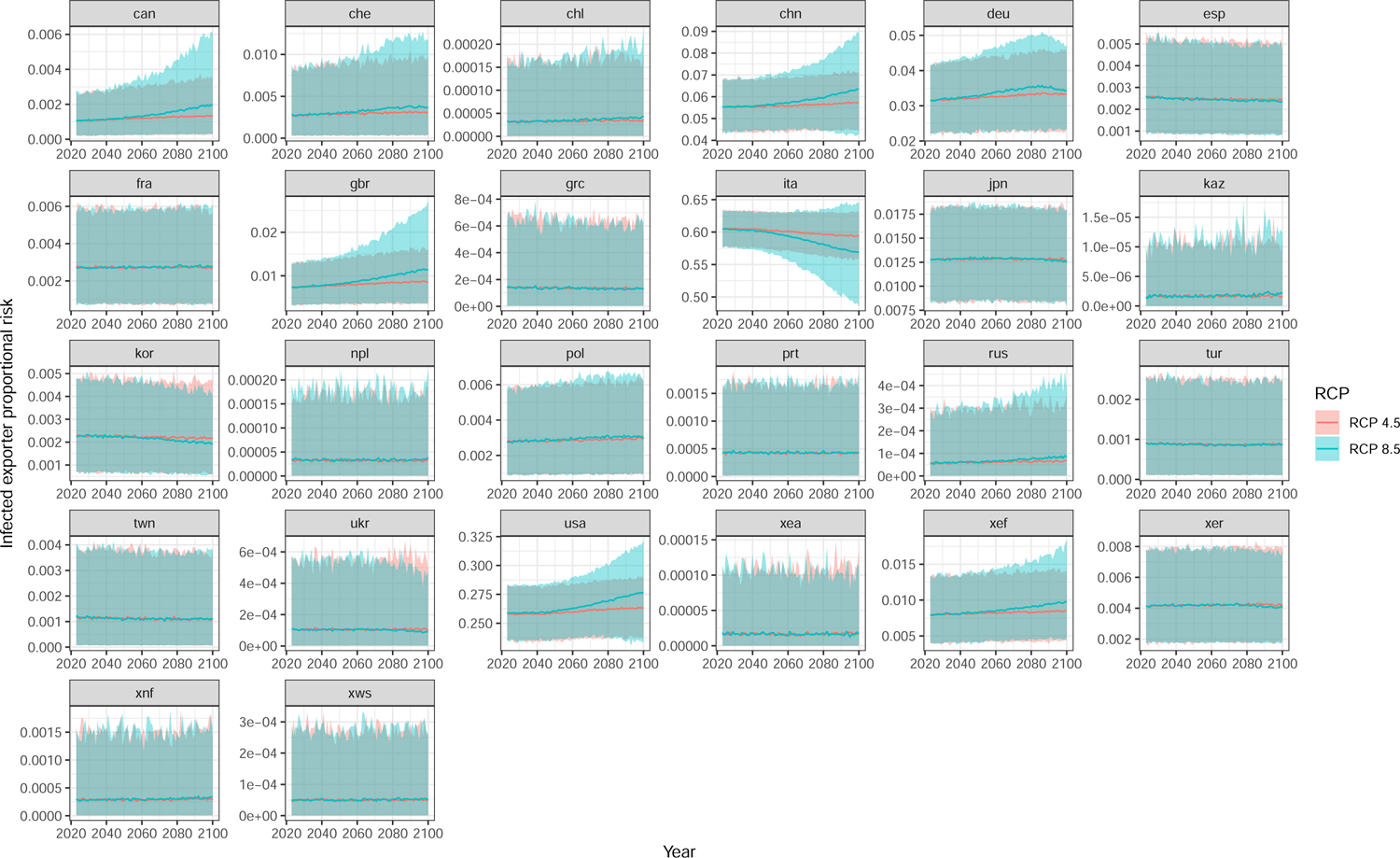
Annual mean (95% CI) exporter proportional contributions to brown marmorated stink bug (*Halyomorpha halys*) propagule pressure hitting Australia under RCP 4.5 and 8.5. Note: Predictions include stochastic uncertainty associated with future years and for unobserved commodity and infected exporter combinations.

**Figure 7.6.:**
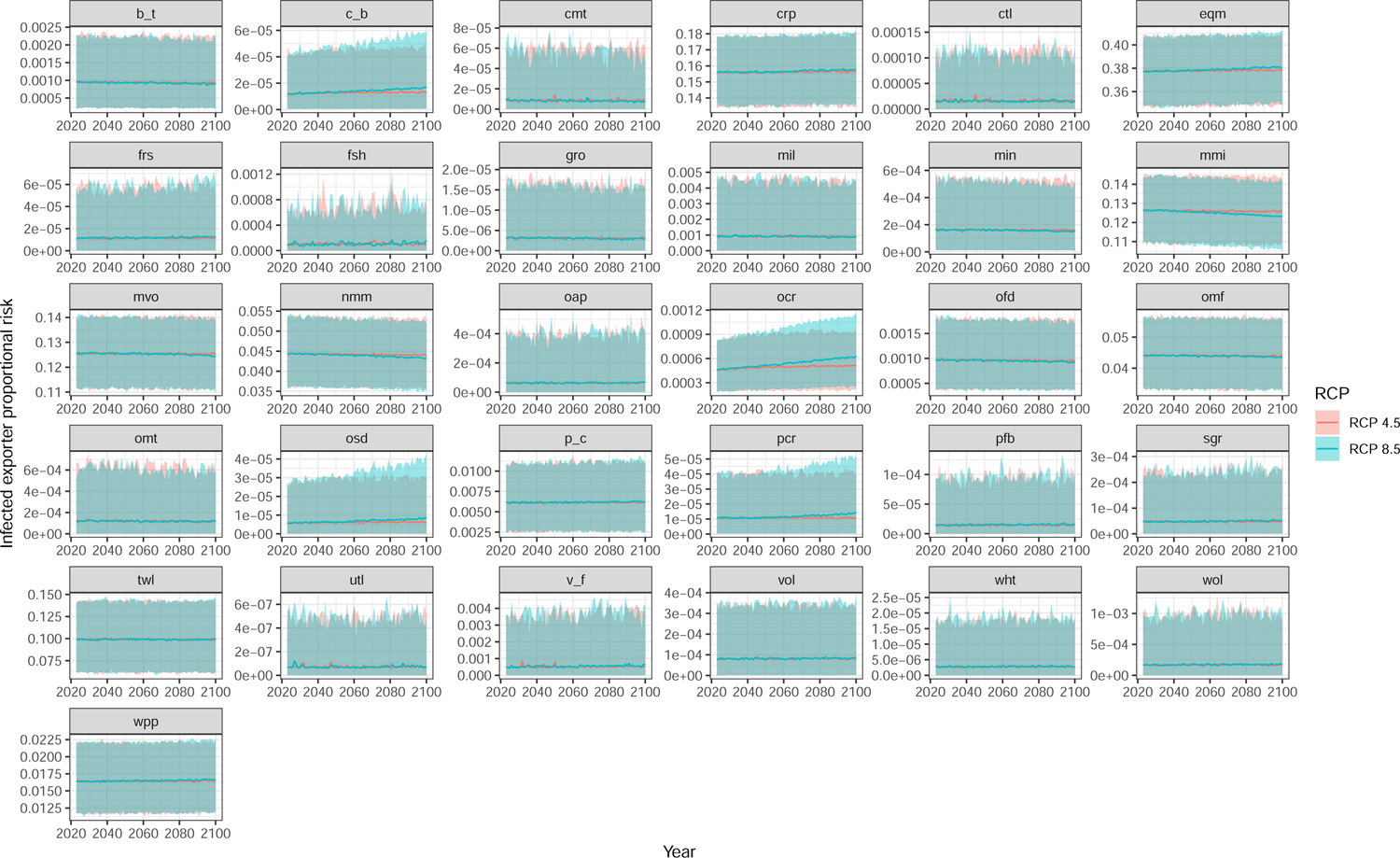
Annual mean (95% CI) commodity sector proportional contributions to brown marmorated stink bug (*Halyomorpha halys*) propagule pressure hitting Australia under RCP 4.5 and 8.5. Note: Predictions include stochastic uncertainty associated with future years and for unobserved commodity and infected exporter combinations.

## 8. Egg laying pest: Spongy moth (*Lymantria dispar*)

### 8.1. Likelihoods of contamination

The spongy moth (*Lymantria dispar*) contamination model revealed that regions with higher mean climate suitability (i.e., climsuit*_it_*) tended to have higher likelihoods of contamination. However, this effect was highly uncertain (confidence in direction > 50%; Figure 8.1a), likely due to the very low interceptions detected at the border (Table 6.1). The model also revealed that there was unexplained variation associated with random grouping variables (Figure 8.1b). Specifically, contamination rates were most variable (and uncertain) among years (Year SD = 1.13), followed by exporter region (Region SD = 0.93) and least variable among commodity sectors (Commodity SD = 0.51).

**Figure 8.1.:**
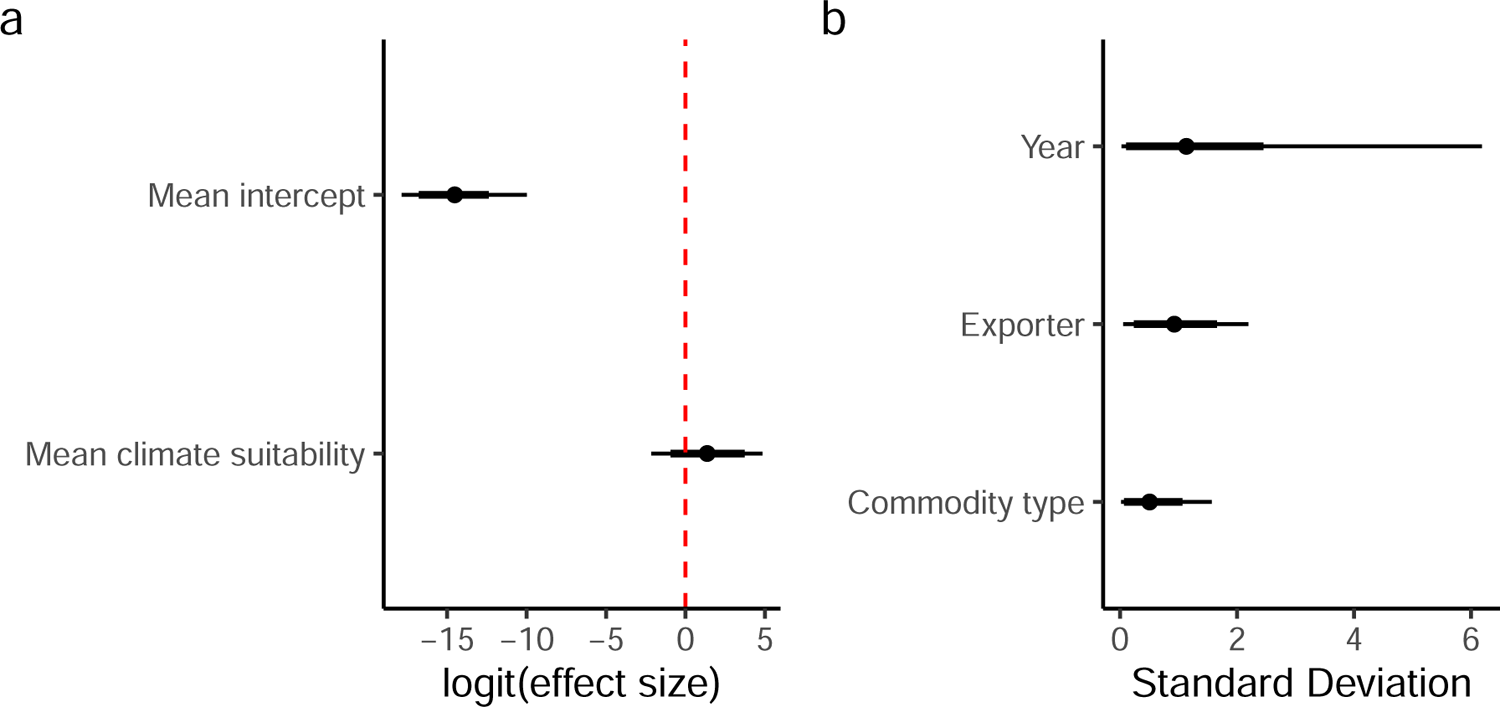
Spongy moth (*Lymantria dispar*) coefficient and group variance plots. a) Model coefficients; b) Random effect standard deviations (i.e. measure of variability within each grouping variable). Error bars reflect 95% (thin) and 80% (thick) credible intervals.

Examination of baseline likelihoods (Figure 8.2) revealed very low contamination rates (mean estimates all < 1e-05) across both exporters and commodity sectors. The model also found little difference in contamination likelihoods among exporters or commodity sectors. This is driven by the fact that there has so very few detections of this species at the Australian border, despite importing significant amounts of commodities from these regions.

**Figure 8.2.:**
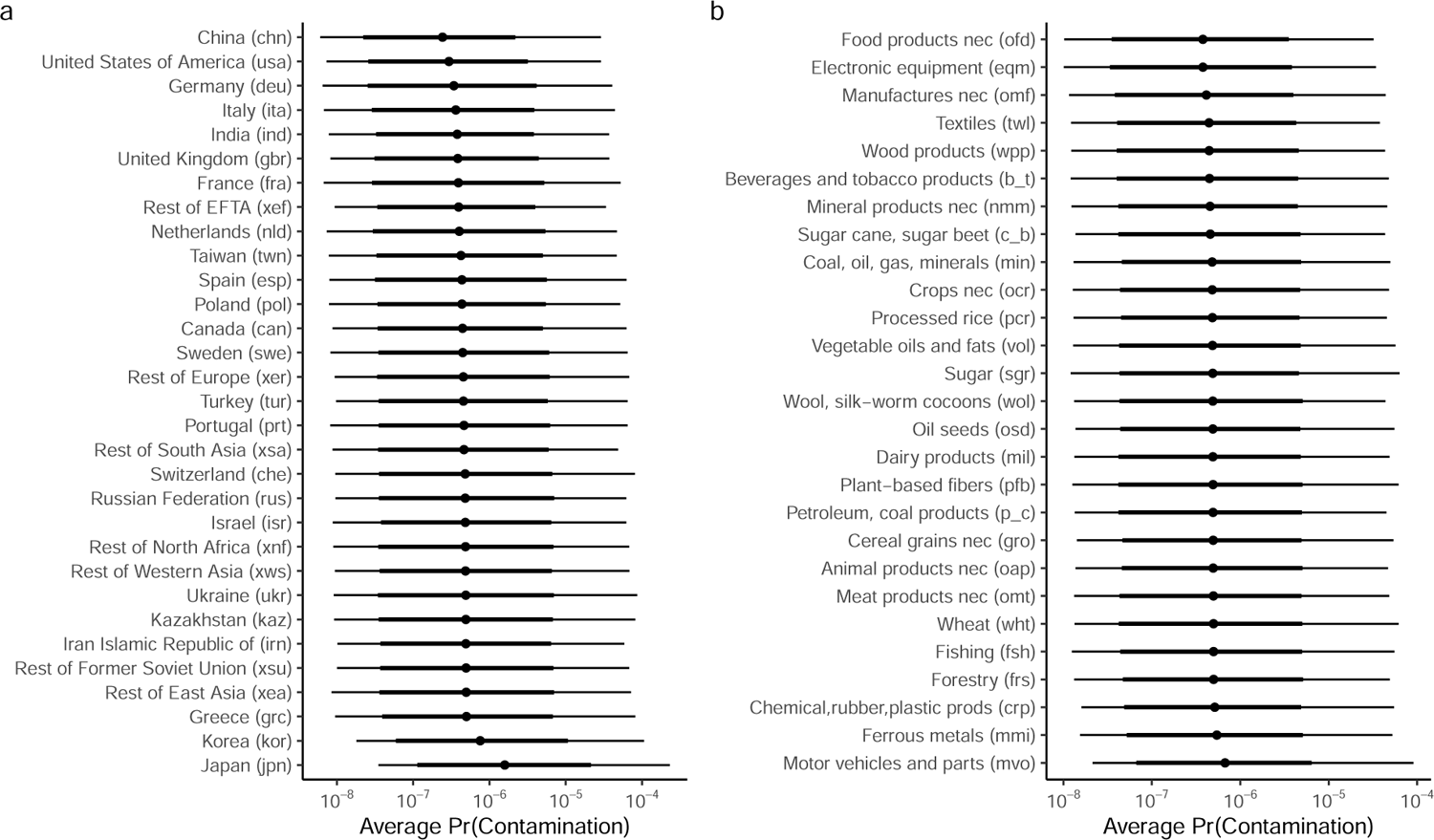
Spongy moth (*Lymantria dispar*) baseline contamination likelihoods associated with (a) infected exporter; and (b) commodity type. Error bars reflect 95% (thin) and 80% (thick) credible invervals. Note: baseline likelihoods are back-transformed (anti-logit) intercept terms in the model and that do not account for exporter mean climate suitability.

Spongy moth contamination rates also did not vary substantially between 2014 and 2021 (Figure 8.3), with contamination rates marginally higher in 2014, 2020, and 2021 – the years where border detections had occurred.

**Figure 8.3.:**
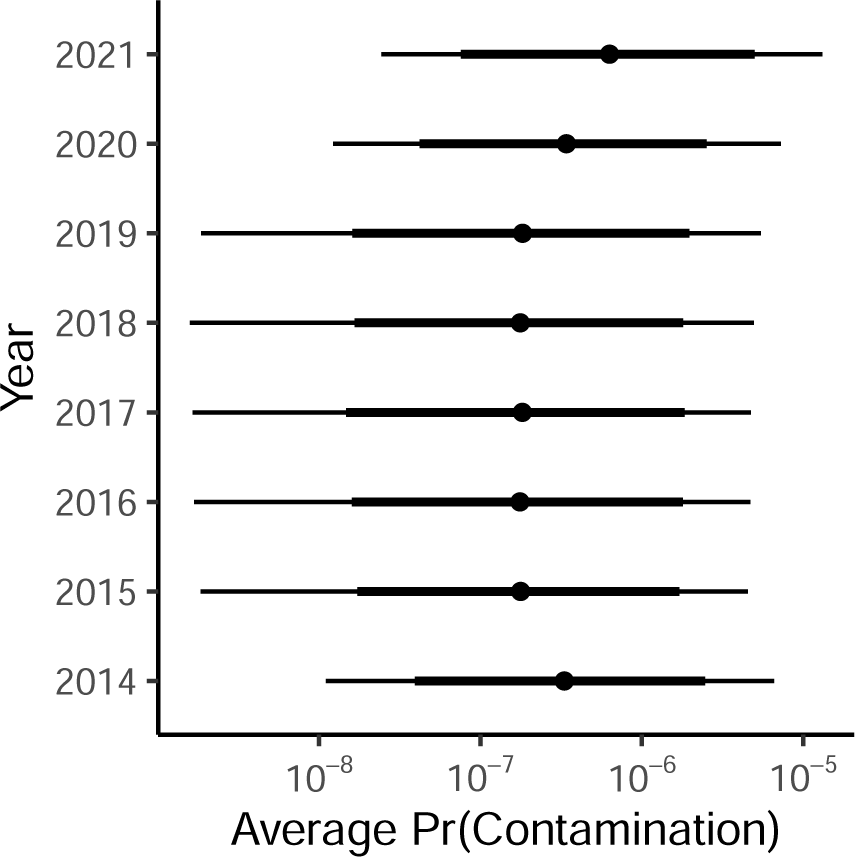
Spongy moth annual baseline contamination likelihoods. Error bars reflect 95% (thin) and 80% (thick) credible invervals. Note: baseline likelihoods are back-transformed (anti-logit) intercept terms in the model and that do not account for exporter mean climate suitability.

Imports from Japan (jpn) accounted for 19 of the top 20 estimated contamination likelihoods for 2023 (Table 8.1). However, it is important to note that all estimated contamination rates were exceptionally low, with imports of motor vehicles and associated parts (mvo) from Japan containing the highest estimated contamination rate at only 0.0007%. This might be because this species has been known to have egg masses on ships as opposed to traded commodities, and as such, have interceptions that are not attributed to a commodity sector.

While contamination rates were very low for this species, the model predicted further declines in contamination rates over time for most infected regions. This is because mean climate suitability in most infected exporter regions are expected to significantly or marginally decline (Table 8.2). For example, mean climate suitability for this species in Japan (jpn; the region with the highest predicted contamination rates) is expected to decline significantly from 0.91 (0.88-0.94) to 0.81 (0.73-0.88) for RCP 4.5 and 0.66 (0.54-0.77) for RCP 8.5 by 2100.

However, exceptions do exist. Canada (can) for example, is predicted to become more suitable for spongy moth with mean climate suitability increasing from 0.74 (0.71- 0.77) to 0.8 (0.74-0.87) for RCP 4.5 and 0.83 (0.71-0.91) for RCP 8.5 by 2100.

**Table 8.1.:**
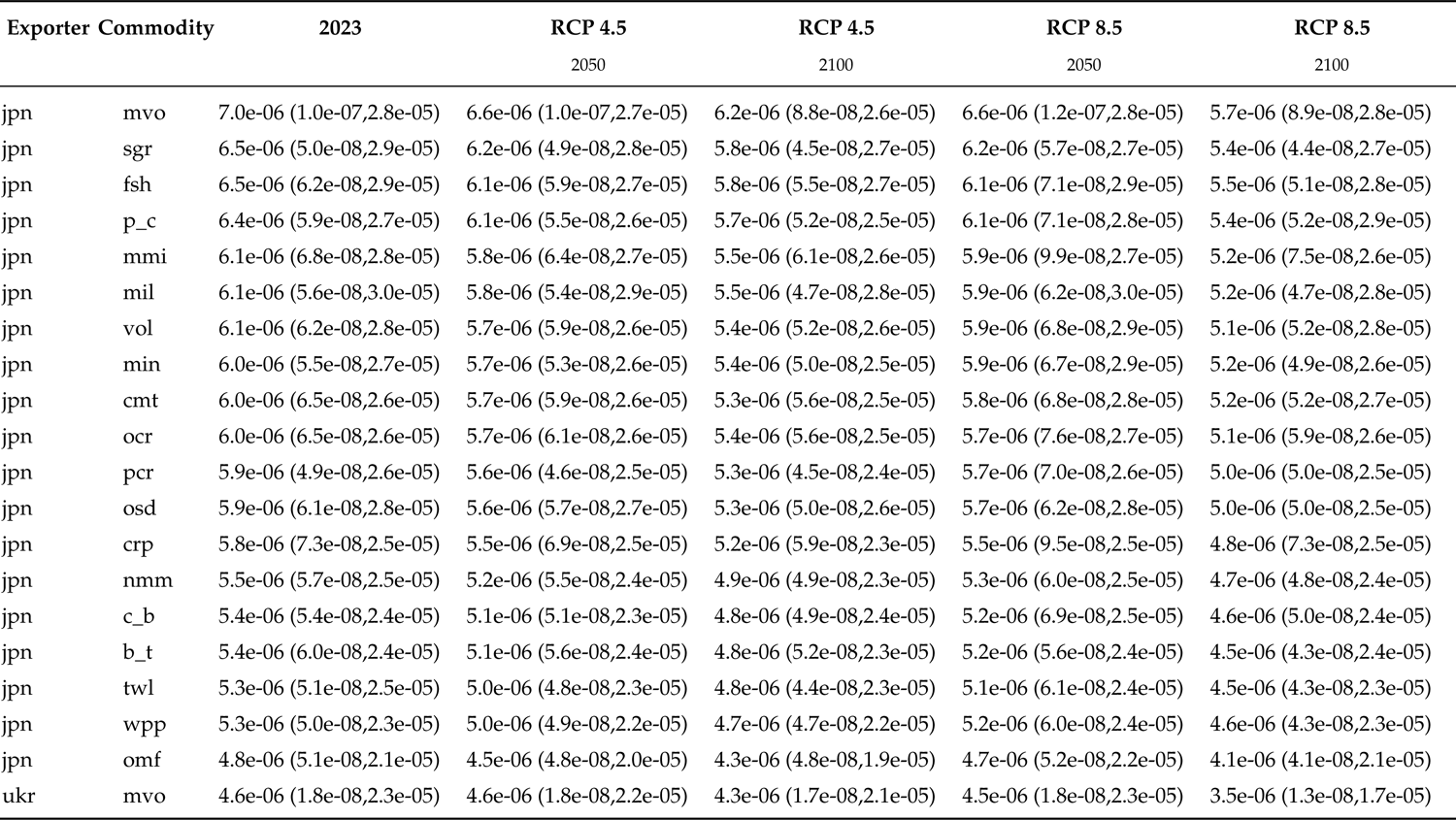
Spongy moth top 20 commodity sector by region contamination likelihoods for commodities with 100 or more imported lines into Australia and how they are predicted to change under RCP 4.5 and RCP 8.5 by 2050 and 2100. Numbers in brackets signify 95% credible limits.

**Table 8.2.:** Spongy moth mean climate suitability for each infected exporter region in 2023 and 2100 for RCPs 4.5 and 8.5. Mean estimates are based on the ensembled annual climate suitabilities derived from 23 GCMs. Numbers in parentheses are the 10th and 90th percentiles derived from ensembles annual models.

| Region | 2023 | RCP 4.5 | RCP 8.5 |
| --- | --- | --- | --- |
|  |  | 2100 | 2100 |
| Canada (can) | 0.74 (0.71-0.77) | 0.8 (0.74-0.87) | 0.83 (0.71-0.91) |
| Switzerland (che) | 0.95 (0.93-0.96) | 0.96 (0.92-0.98) | 0.9 (0.79-0.98) |
| China (chn) | 0.66 (0.62-0.68) | 0.62 (0.55-0.68) | 0.56 (0.48-0.64) |
| Germany (deu) | 1 (1-1) | 0.99 (0.97-1) | 0.9 (0.75-0.99) |
| Spain (esp) | 0.9 (0.87-0.92) | 0.77 (0.71-0.83) | 0.6 (0.5-0.7) |
| France (fra) | 0.98 (0.98-0.98) | 0.94 (0.91-0.97) | 0.79 (0.63-0.92) |
| United Kingdom (gbr) | 0.95 (0.93-0.96) | 0.95 (0.93-0.97) | 0.94 (0.89-0.97) |
| Greece (grc) | 0.92 (0.9-0.94) | 0.77 (0.7-0.84) | 0.57 (0.5-0.66) |
| India (ind) | 0.3 (0.27-0.34) | 0.25 (0.21-0.29) | 0.22 (0.17-0.26) |
| Iran Islamic Republic of (irn) | 0.44 (0.41-0.46) | 0.37 (0.33-0.41) | 0.3 (0.26-0.34) |
| Israel (isr) | 0.36 (0.34-0.38) | 0.27 (0.24-0.31) | 0.2 (0.18-0.22) |
| Italy (ita) | 0.96 (0.95-0.98) | 0.84 (0.77-0.92) | 0.64 (0.55-0.74) |
| Japan (jpn) | 0.91 (0.88-0.94) | 0.81 (0.73-0.88) | 0.66 (0.54-0.77) |
| Kazakhstan (kaz) | 0.66 (0.64-0.69) | 0.6 (0.54-0.65) | 0.49 (0.43-0.55) |
| Korea (kor) | 0.93 (0.88-0.96) | 0.77 (0.64-0.89) | 0.6 (0.45-0.75) |
| Netherlands (nld) | 1 (1-1) | 0.99 (0.97-1) | 0.94 (0.85-1) |
| Poland (pol) | 1 (1-1) | 0.99 (0.98-1) | 0.9 (0.75-0.99) |
| Portugal (prt) | 0.95 (0.94-0.97) | 0.83 (0.76-0.9) | 0.63 (0.52-0.76) |
| Russian Federation (rus) | 0.72 (0.7-0.75) | 0.77 (0.71-0.83) | 0.77 (0.66-0.87) |
| Sweden (swe) | 1 (0.99-1) | 1 (0.99-1) | 0.99 (0.97-1) |
| Turkey (tur) | 0.9 (0.89-0.91) | 0.81 (0.75-0.87) | 0.63 (0.54-0.72) |
| Taiwan (twn) | 0.38 (0.36-0.4) | 0.34 (0.31-0.37) | 0.29 (0.25-0.33) |
| Ukraine (ukr) | 1 (0.99-1) | 0.96 (0.91-0.99) | 0.75 (0.62-0.87) |
| United States of America (usa) | 0.8 (0.77-0.83) | 0.73 (0.67-0.79) | 0.63 (0.55-0.71) |
| Rest of East Asia (xea) | 0.59 (0.55-0.62) | 0.57 (0.51-0.64) | 0.52 (0.45-0.59) |
| Rest of EFTA (xef) | 0.35 (0.35-0.35) | 0.34 (0.33-0.35) | 0.31 (0.27-0.34) |
| Rest of Europe (xer) | 0.88 (0.88-0.88) | 0.84 (0.8-0.87) | 0.67 (0.57-0.77) |
| Rest of North Africa (xnf) | 0.16 (0.15-0.17) | 0.13 (0.11-0.14) | 0.1 (0.09-0.12) |
| Rest of South Asia (xsa) | 0.43 (0.41-0.44) | 0.4 (0.37-0.43) | 0.35 (0.31-0.4) |
| Rest of Former Soviet Union (xsu) | 0.48 (0.46-0.5) | 0.46 (0.42-0.49) | 0.42 (0.38-0.46) |
| Rest of Western Asia (xws) | 0.15 (0.14-0.15) | 0.13 (0.12-0.14) | 0.11 (0.09-0.12) |

### 8.2. Australia: Spongy moth propagule pressure and establishment exposure

Annual spongy moth propagule pressure arriving at Australia’s border is expected to remain fairly stable, with only a marginal decline detected for RCP 8.5 where propagule pressure declines from 32 (95CI: 1-106) in 2023 to 29 (95CI: 1-108) by 2100 (Figure 8.4). This relatively stable propagule pressure is mostly attributable to little variability in contamination rates among infected exporters or commodity sectors.

**Figure 8.4.:**
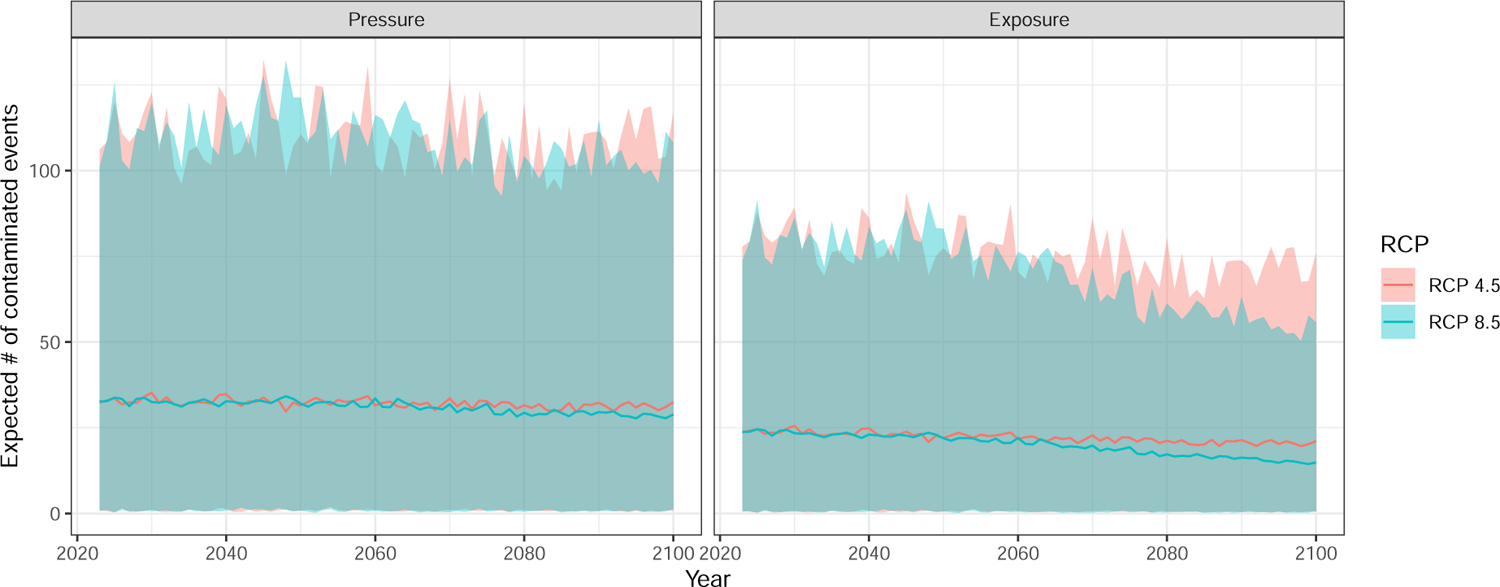
Annual mean (95% CI) spongy moth (*Lymantria dispar*) propagule pressure (left) and establishment exposure (right) hitting Australia under RCP 4.5 and 8.5. Note: Predictions include stochastic uncertainty associated with future years and for unobserved commodity and infected exporter combinations.

Estimated establishment exposure in Australia is expected to decline under RCP 8.5 but remain relatively stable under RCP 4.5. This is because under RCP 8.5 there is a significant decline in spongy moth suitable climatic conditions at locations where imported goods are expected to be unloaded (i.e., regions of highest human population density; Figure 6.3). As a consequence, the number of contaminated lines expected to arrive at a climatically suitable location within Australia will be lower.

### 8.3. Australia: Spongy moth risk attributable to different exporters & commodity sectors

Our model indicates that approximately 46% of the spongy moth propagule pressure hitting Australian borders can be attributed to just three exporters: Japan (jpn), China (chn), and Germany (deu;). We also see that for the most part there is little change in infected exporter contributions over time under both RCPs (Figure 8.5).

Similarly, risk attributable to different commodity sectors was also expected to remain relatively stable under both RCP scenarios (Figure 8.6). Imports of electronic equipment (eqm) was predicted to be the largest contributor to spongy moth propagule pressure, accounting for approximately 25% (4-43) of the total number of predicted spongy moth contaminated lines entering Australia. This was followed by Chemical, rubber and plastic products (crp; 16% (5-36)); ferrous metals (mmi; 13% (5-31)) and textiles (twl; 11% (2-26)).

**Figure 8.5.:**
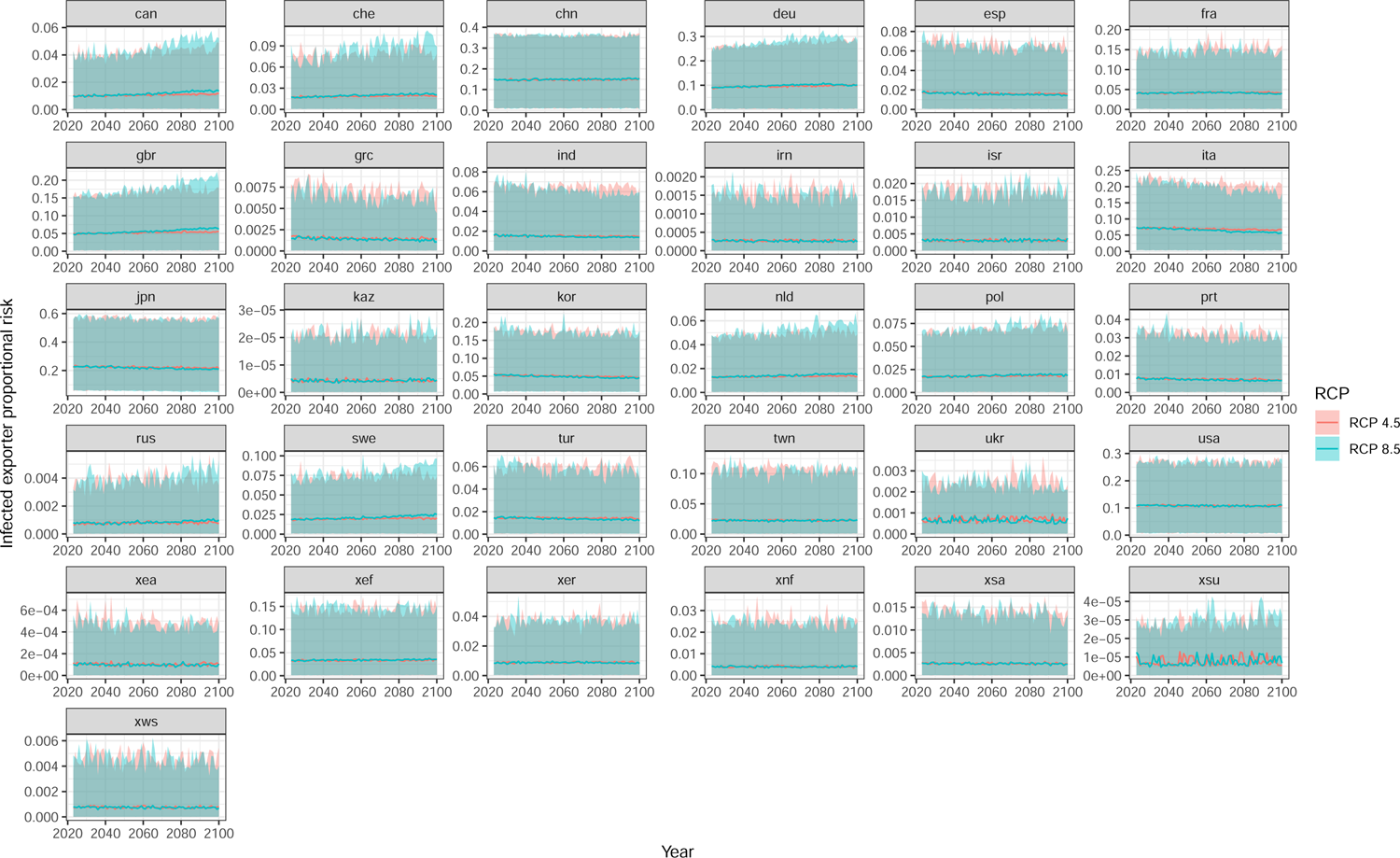
Annual mean (95% CI) exporter proportional contributions to spongy moth (*Lymantria dispar*) propagule pressure hitting Australia under RCP 4.5 and 8.5. Note: Predictions include stochastic uncertainty associated with future years and for unobserved commodity and infected exporter combinations.

**Figure 8.6.:**
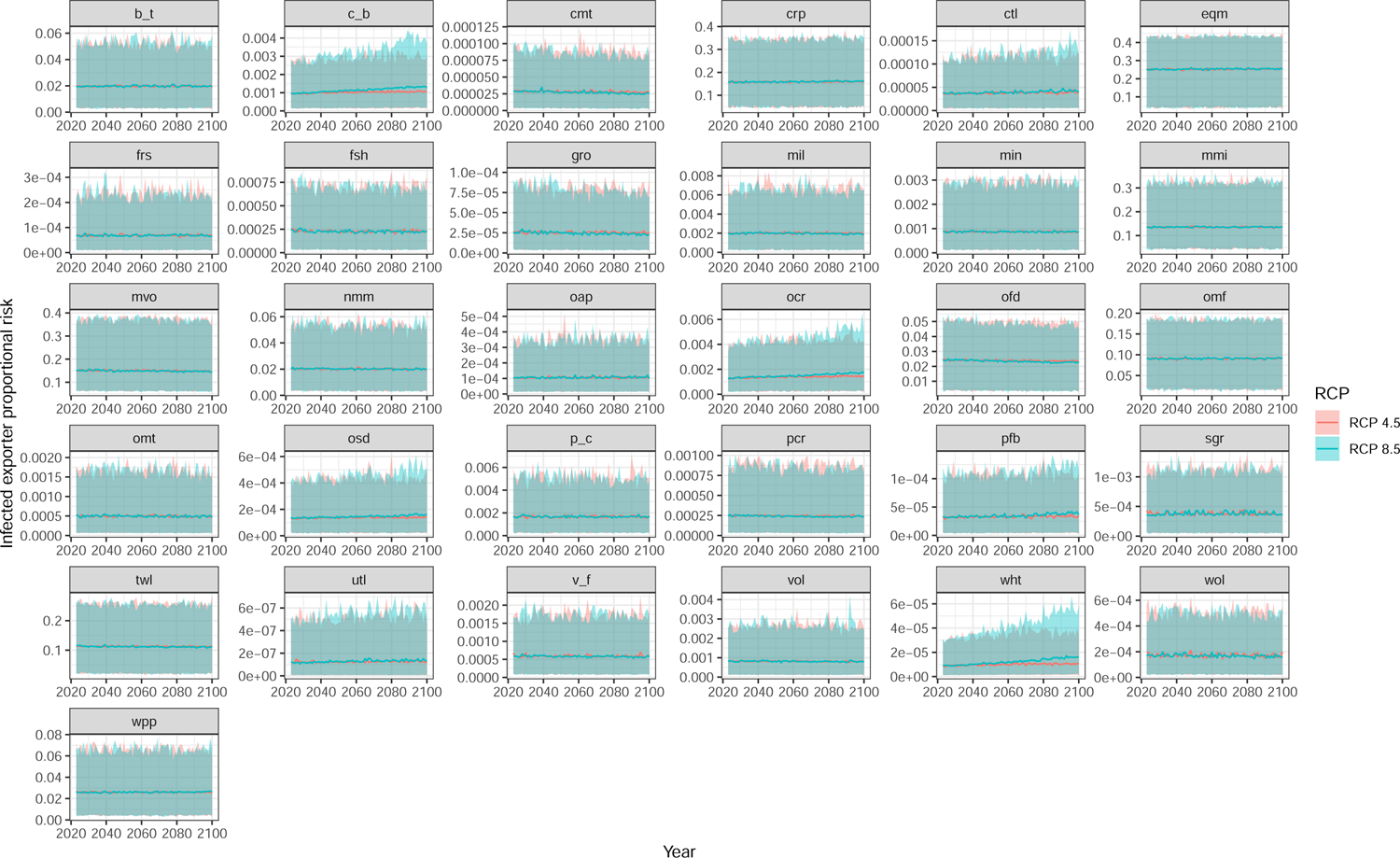
Annual mean (95% CI) commodity sector proportional contributions to spongy moth (*Lymantria dispar*) propagule pressure hitting Australia under RCP 4.5 and 8.5. Note: Predictions include stochastic uncertainty associated with future years and for unobserved commodity and infected exporter combinations.

## 9. Nesting pest: Asian honey bee (*Apis cerana*)

### 9.1. Likelihoods of contamination

The Asian honey bee contamination model revealed that there was neglible effect of exporter mean climate suitability (i.e., climsuit*_it_*) on contamination rates (Figure 9.1a). This is likely because the species has a relatively wide potential distribution of suitable climate in infected regions that commonly export goods to Australia (Figure 6.1) coupled with the fact that there has been few interceptions detected at the border (Table 6.1). The model revealed that contamination rates were most variable among commodity sector (Commodity SD = 1.56) and exporter region (Region SD = 1.55) and less variable among years (Year SD = 0.45).

**Figure 9.1.:**
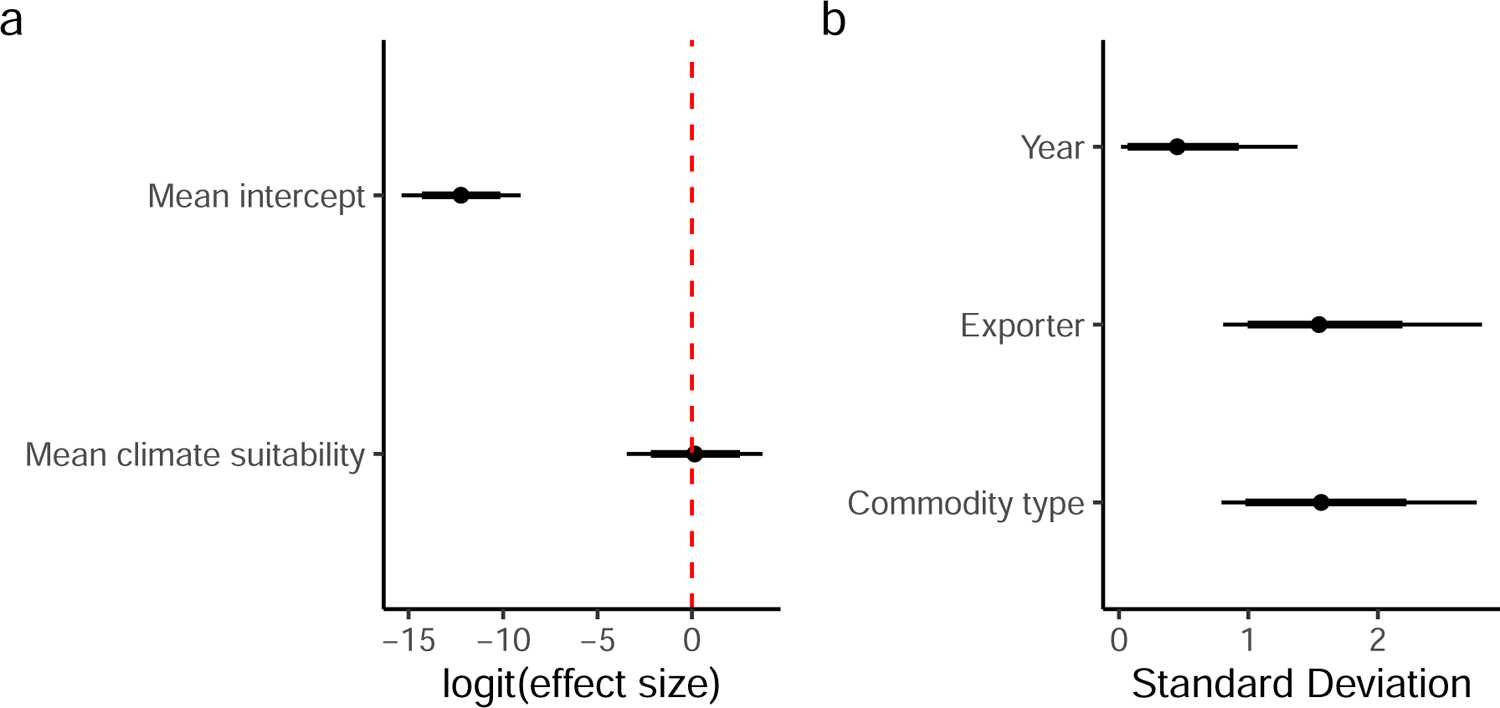
Asian honey bee (*Apis cerana*) coefficient and group variance plots. a) Model coefficients; b) Random effect standard deviations (i.e. measure of variability within each grouping variable). Error bars reflect 95% (thin) and 80% (thick) credible intervals.

Examination of baseline likelihoods (Figure 9.2) revealed very low contamination rates (mean estimates all < 1e-04) across both exporters and commodity sectors. The model also predicted little variation in contamination likelihoods among most exporters with Malaysia (mys) and Rest of Oceania (xoc) expected to have marginally higher baseline contamination rates relative to other exporters. Similarly, there was also little variation among commodity sectors, with electronic equipment (eqm); motor vehicles and associated parts (mvo); and Chemical, rubber and plastic products (crp) exhibiting marginally higher contamination likelihoods. Much like spongy moth, these low contamination rates is largely driven by the fact that there were few detections of this species at the Australian border, despite importing significant amounts of commodities from these regions.

**Figure 9.2.:**
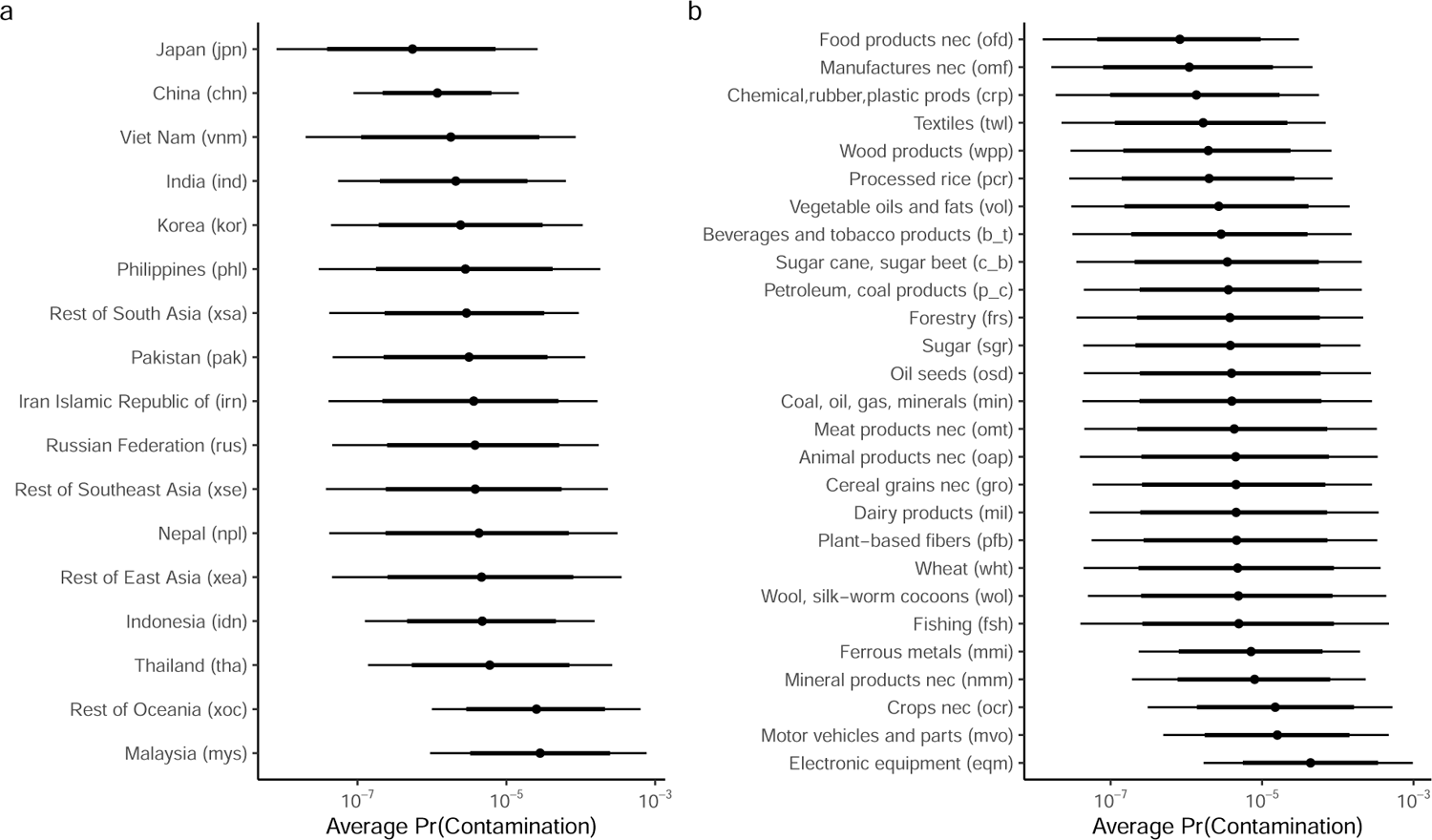
Asian honey bee (*Apis cerana*) baseline contamination likelihoods associated with (a) infected exporter; and (b) commodity type. Error bars reflect 95% (thin) and 80% (thick) credible invervals. Note: baseline likelihoods are back- transformed (anti-logit) intercept terms in the model and that do not account for exporter mean climate suitability.

**Figure 9.3.:**
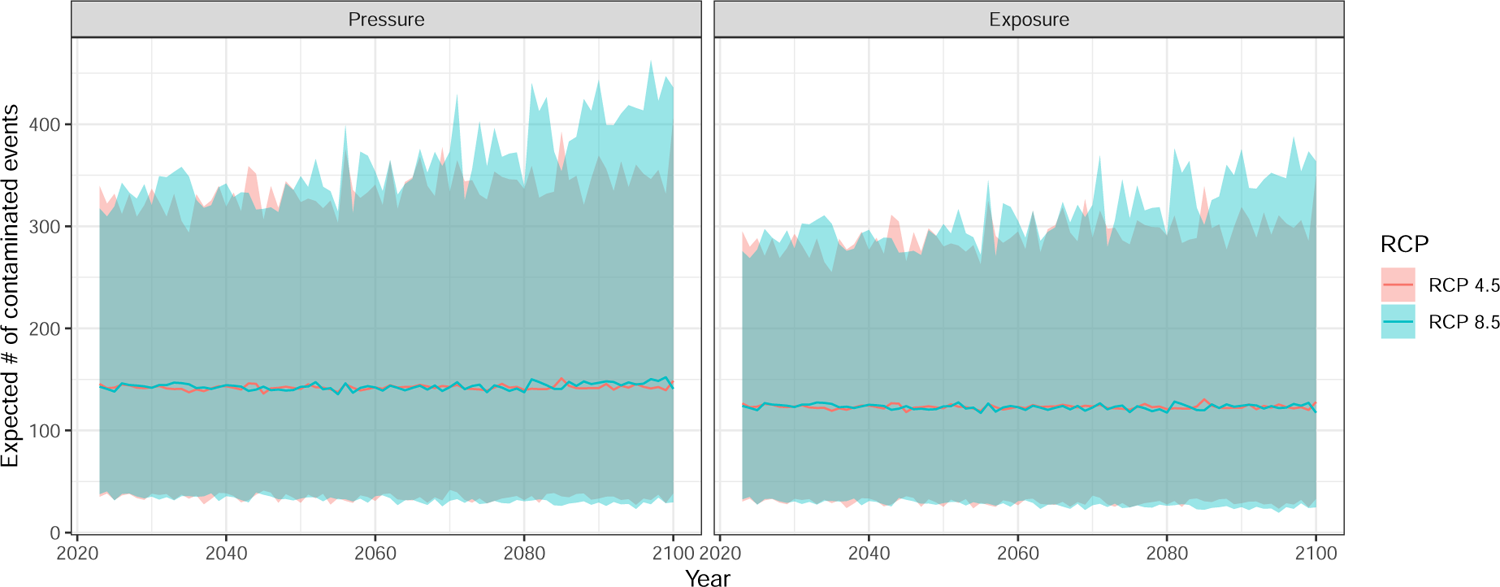
Annual mean (95% CI) Asian honey bee (*Apis cerana*) propagule pressure (left) and establishment exposure (right) hitting Australia under RCP 4.5 and 8.5. Note: Predictions include stochastic uncertainty associated with future years and for unobserved commodity and infected exporter combinations.

Imports of electronic equipment (eqm) from South and East Asia (xea, xse), Malaysia (mys), Rest of Oceania (xoc), Russia (rus), Iran (irn), and Thailand (tha) accounted for 7 of the top 20 estimated contamination likelihoods for 2023 (Table 9.1). However, in all cases, estimated contamination rates were low (means < 0.4%). As climate suitability in infected regions was not found to have significant impacts on contamination rates, minimal change was predicted over time under both climate scenarios.

**Table 9.1.:** Asian honey bee top 20 commodity sector by region contamination likelihoods for commodities with 100 or more imported lines into Australia and how they are predicted to change under RCP 4.5 and RCP 8.5 by 2050 and 2100. Numbers in brackets signify 95% credible limits.

| Exporter Commodity |  | 2023 | RCP 4.5 | RCP 4.5 | RCP 8.5 | RCP 8.5 |
| --- | --- | --- | --- | --- | --- | --- |
|  |  |  | 2050 | 2100 | 2050 | 2100 |
| xea | eqm | 3.4e-04 (3.9e-07,2.7e-03) | 3.4e-04 (4.0e-07,2.7e-03) | 3.4e-04 (4.0e-07,2.7e-03) | 3.3e-04 (3.8e-07,2.7e-03) | 3.3e-04 (3.8e-07,2.7e-03) |
| mys | eqm | 3.3e-04 (5.6e-05,9.0e-04) | 3.3e-04 (5.3e-05,9.7e-04) | 3.5e-04 (4.6e-05,1.1e-03) | 3.4e-04 (4.8e-05,1.1e-03) | 4.0e-04 (3.2e-05,1.6e-03) |
| xoc | eqm | 3.0e-04 (4.4e-05,8.6e-04) | 3.0e-04 (4.3e-05,9.0e-04) | 3.1e-04 (4.0e-05,9.8e-04) | 3.1e-04 (3.9e-05,9.6e-04) | 3.4e-04 (3.2e-05,1.2e-03) |
| mys | ocr | 2.0e-04 (5.7e-06,1.1e-03) | 2.0e-04 (5.4e-06,1.1e-03) | 2.1e-04 (5.0e-06,1.2e-03) | 2.1e-04 (5.0e-06,1.2e-03) | 2.4e-04 (3.9e-06,1.4e-03) |
| xoc | ocr | 1.8e-04 (5.0e-06,9.4e-04) | 1.8e-04 (4.8e-06,9.3e-04) | 1.8e-04 (4.7e-06,9.6e-04) | 1.8e-04 (5.0e-06,9.3e-04) | 2.0e-04 (4.2e-06,1.0e-03) |
| mys | fsh | 1.8e-04 (7.0e-07,1.0e-03) | 1.8e-04 (6.9e-07,1.0e-03) | 1.9e-04 (6.4e-07,1.0e-03) | 1.9e-04 (6.9e-07,1.1e-03) | 2.2e-04 (5.8e-07,1.3e-03) |
| xoc | fsh | 1.8e-04 (5.4e-07,9.9e-04) | 1.8e-04 (5.2e-07,9.8e-04) | 1.8e-04 (5.1e-07,9.6e-04) | 1.9e-04 (5.3e-07,1.1e-03) | 2.1e-04 (4.7e-07,1.2e-03) |
| mys | mvo | 1.5e-04 (1.3e-05,5.4e-04) | 1.5e-04 (1.2e-05,5.8e-04) | 1.5e-04 (1.1e-05,6.4e-04) | 1.5e-04 (1.1e-05,6.0e-04) | 1.8e-04 (7.6e-06,8.8e-04) |
| xoc | mvo | 1.3e-04 (1.1e-05,4.6e-04) | 1.3e-04 (1.1e-05,4.7e-04) | 1.3e-04 (9.9e-06,5.0e-04) | 1.3e-04 (9.3e-06,5.0e-04) | 1.5e-04 (7.5e-06,6.5e-04) |
| xoc | v_f | 1.2e-04 (4.6e-07,5.5e-04) | 1.2e-04 (4.6e-07,5.6e-04) | 1.2e-04 (4.3e-07,5.7e-04) | 8.9e-05 (4.9e-07,5.2e-04) | 9.7e-05 (4.1e-07,6.6e-04) |
| rus | eqm | 1.2e-04 (8.5e-07,7.9e-04) | 1.2e-04 (8.6e-07,7.6e-04) | 1.2e-04 (9.1e-07,7.4e-04) | 1.2e-04 (9.5e-07,7.7e-04) | 1.1e-04 (1.1e-06,7.6e-04) |
| irn | eqm | 1.1e-04 (6.5e-07,7.6e-04) | 1.2e-04 (6.3e-07,7.8e-04) | 1.2e-04 (6.2e-07,8.0e-04) | 1.2e-04 (5.9e-07,8.9e-04) | 1.3e-04 (5.0e-07,9.0e-04) |
| xse | eqm | 1.1e-04 (1.1e-06,6.9e-04) | 1.1e-04 (1.2e-06,6.9e-04) | 1.1e-04 (1.2e-06,6.7e-04) | 1.1e-04 (1.0e-06,7.0e-04) | 1.2e-04 (9.0e-07,6.9e-04) |
| xoc | oap | 9.6e-05 (5.2e-07,5.8e-04) | 9.6e-05 (5.0e-07,5.8e-04) | 9.7e-05 (4.6e-07,6.0e-04) | 9.9e-05 (4.7e-07,6.7e-04) | 1.1e-04 (3.9e-07,7.5e-04) |
| mys | nmm | 8.9e-05 (4.0e-06,4.1e-04) | 9.0e-05 (3.7e-06,4.2e-04) | 9.3e-05 (3.4e-06,4.5e-04) | 9.3e-05 (3.3e-06,4.3e-04) | 1.1e-04 (2.4e-06,5.8e-04) |
| npl | ocr | 8.5e-05 (1.6e-07,6.3e-04) | 8.5e-05 (1.6e-07,6.2e-04) | 8.5e-05 (1.6e-07,6.1e-04) | 8.7e-05 (1.8e-07,6.5e-04) | 8.5e-05 (1.8e-07,6.2e-04) |
| mys | min | 8.5e-05 (6.0e-07,5.7e-04) | 8.4e-05 (5.4e-07,5.8e-04) | 8.7e-05 (4.8e-07,6.2e-04) | 8.8e-05 (5.1e-07,5.9e-04) | 1.0e-04 (4.0e-07,7.2e-04) |
| xoc | nmm | 8.2e-05 (3.0e-06,3.8e-04) | 8.2e-05 (3.1e-06,4.0e-04) | 8.4e-05 (3.0e-06,4.0e-04) | 8.5e-05 (2.8e-06,4.1e-04) | 9.3e-05 (2.4e-06,4.8e-04) |
| xoc | min | 8.2e-05 (4.1e-07,5.4e-04) | 8.2e-05 (4.2e-07,5.4e-04) | 8.3e-05 (4.2e-07,5.6e-04) | 8.5e-05 (3.9e-07,5.6e-04) | 9.1e-05 (3.6e-07,6.5e-04) |
| tha | eqm | 8.1e-05 (7.9e-06,2.7e-04) | 8.0e-05 (7.5e-06,2.6e-04) | 8.2e-05 (7.3e-06,2.8e-04) | 8.2e-05 (8.3e-06,2.8e-04) | 9.8e-05 (5.5e-06,4.6e-04) |

However, it is important to note, that climate suitability for this species is expected to decline in most regions over time under both climate scenarios (Figure 6.4), with potential consequences on the long-term extent and population dynamics in infected regions (Table 9.2).

**Table 9.2.:** Asian honey bee mean climate suitability for each infected exporter region in 2023 and 2100 for RCPs 4.5 and 8.5. Mean estimates are based on the ensembled annual climate suitabilities derived from 23 GCMs. Numbers in parentheses are the 10th and 90th percentiles derived from ensembles annual models. Note: Australia has been included in this table due to a local contained population established near Cairns.

| Region | 2023 | RCP 4.5 | RCP 8.5 |
| --- | --- | --- | --- |
|  |  | 2100 | 2100 |
| Australia (aus) | 0.72 (0.69-0.75) | 0.63 (0.56-0.7) | 0.53 (0.45-0.6) |
| China (chn) | 0.56 (0.54-0.57) | 0.58 (0.53-0.63) | 0.6 (0.53-0.67) |
| Indonesia (idn) | 0.8 (0.75-0.85) | 0.63 (0.55-0.72) | 0.5 (0.43-0.58) |
| India (ind) | 0.82 (0.77-0.86) | 0.69 (0.6-0.77) | 0.55 (0.45-0.64) |
| Iran Islamic Republic of (irn) | 0.39 (0.37-0.41) | 0.35 (0.31-0.39) | 0.3 (0.26-0.35) |
| Japan (jpn) | 0.95 (0.94-0.96) | 0.94 (0.91-0.97) | 0.9 (0.81-0.96) |
| Korea (kor) | 0.97 (0.96-0.98) | 0.94 (0.9-0.97) | 0.85 (0.77-0.93) |
| Malaysia (mys) | 0.81 (0.74-0.87) | 0.62 (0.53-0.72) | 0.46 (0.38-0.56) |
| Nepal (npl) | 0.92 (0.9-0.93) | 0.9 (0.85-0.95) | 0.85 (0.77-0.93) |
| Pakistan (pak) | 0.51 (0.48-0.55) | 0.47 (0.42-0.52) | 0.43 (0.37-0.49) |
| Philippines (phl) | 0.92 (0.89-0.94) | 0.75 (0.68-0.84) | 0.57 (0.5-0.65) |
| Russian Federation (rus) | 0.4 (0.37-0.42) | 0.46 (0.4-0.51) | 0.5 (0.42-0.59) |
| Thailand (tha) | 0.96 (0.92-0.98) | 0.78 (0.65-0.89) | 0.57 (0.43-0.72) |
| Viet Nam (vnm) | 0.96 (0.94-0.97) | 0.87 (0.82-0.92) | 0.73 (0.62-0.82) |
| Rest of East Asia (xea) | 0.06 (0.06-0.06) | 0.06 (0.06-0.06) | 0.06 (0.05-0.06) |
| Rest of Oceania (xoc) | 0.77 (0.74-0.81) | 0.64 (0.56-0.73) | 0.51 (0.43-0.6) |
| Rest of South Asia (xsa) | 0.38 (0.35-0.4) | 0.36 (0.33-0.41) | 0.34 (0.3-0.39) |
| Rest of Southeast Asia (xse) | 0.96 (0.94-0.97) | 0.86 (0.78-0.93) | 0.69 (0.56-0.81) |

### 9.2. Australia: Asian honey bee propagule pressure and establishment exposure

Assuming the local population is effectively managed and contained, the average annual Asian honey bee propagule pressure arriving at Australia’s border is expected to remain stable at approximately 141 (95CI: 30-436). This relatively stable propagule pressure is mostly attributable to low variability in contamination rates among infected exporters and commodity sectors, and thus, little influence of changing trade flows. Moreover, as the model estimated negligable effects climate suitability on contamination likelihoods, rates were not affected by increases or decreases in the potential distribution of the species in infected regions.

Estimated establishment exposure in Australia, is also expected to remain stable under both RCPs – suggesting that the decline in suitable climate within Australia from approximately 0.72 to 0.53 (Table 9.2) is mostly occurring in locations where imported goods are unlikely to be unloaded (e.g. regions with low population density; Figure 6.3).

### 9.3. Australia: Asian honey bee risk attributable to different exporters & commodity sectors

The model indicates that Malaysia (mys) currently accounts for approxmately 39% (15- 65) of the total Asian honey bee propagule pressure hitting Australian borders (Figure 9.4). Other significant contributors were China (chn; 23% (7-44)) and Thailand (tha; 13% (2-32)). Exporter attributions to risk were found to remain fairly stable over time under both RCPs. However, the model is predicting marginal mean increases in propagules from China and Korea and slight declines in Malaysia, especially under RCP 8.5.

Similarly, risk attributable to different commodity sectors was also expected to remain relatively stable under both RCP scenarios (Figure 9.5). Imports of electronic equipment (eqm) were estimated to be the most significant contributor to Asian honey bee propagule pressure, accounting for approximately 76% (57-90) of the total number of predicted Asian honey bee contaminated lines entering Australia.

**Figure 9.4.:**
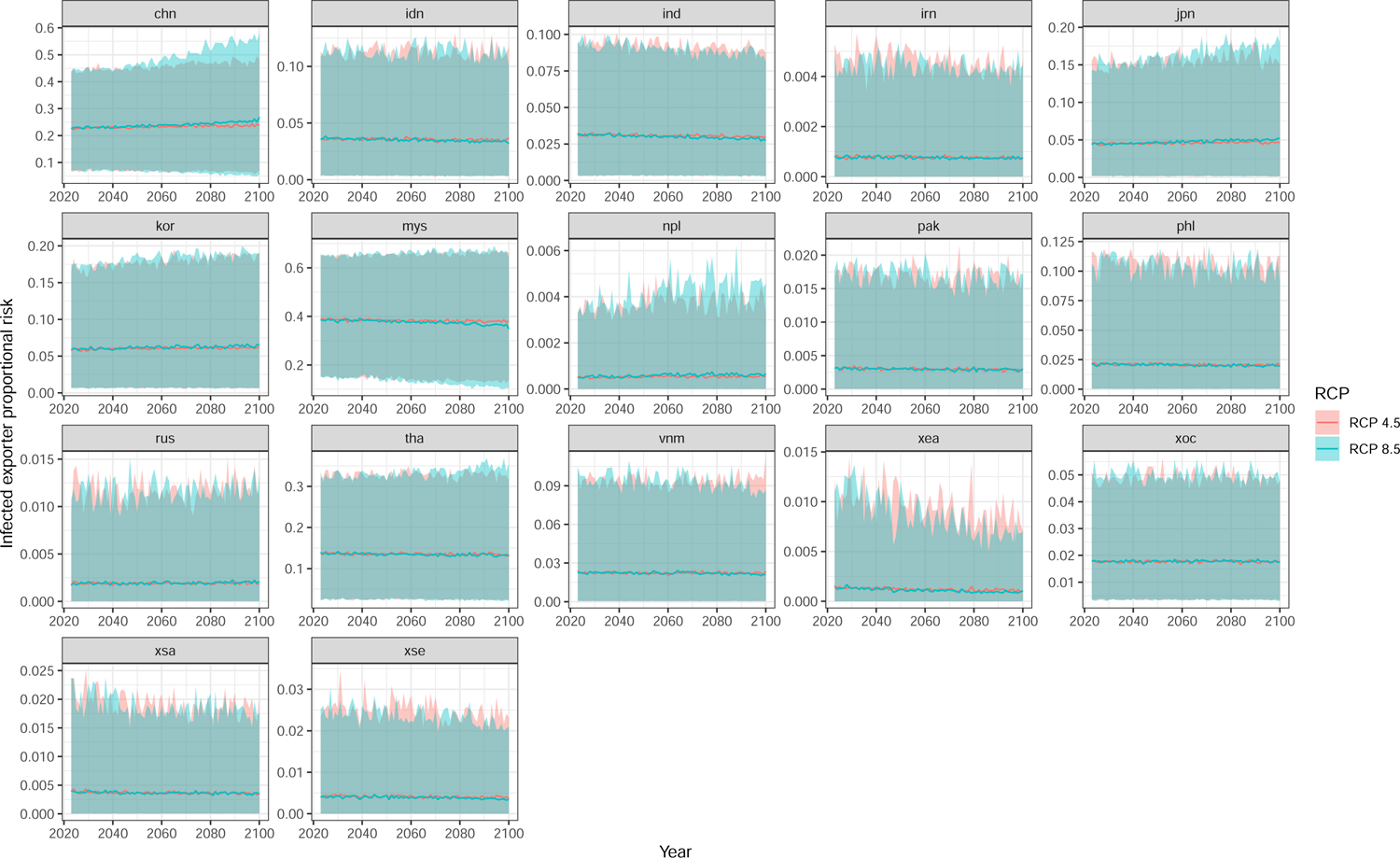
Annual mean (95% CI) exporter proportional contributions to Asian honey bee (*Apis cerana*) propagule pressure hitting Australia under RCP 4.5 and 8.5. Note: Predictions include stochastic uncertainty associated with future years and for unobserved commodity and infected exporter combinations.

**Figure 9.5.:**
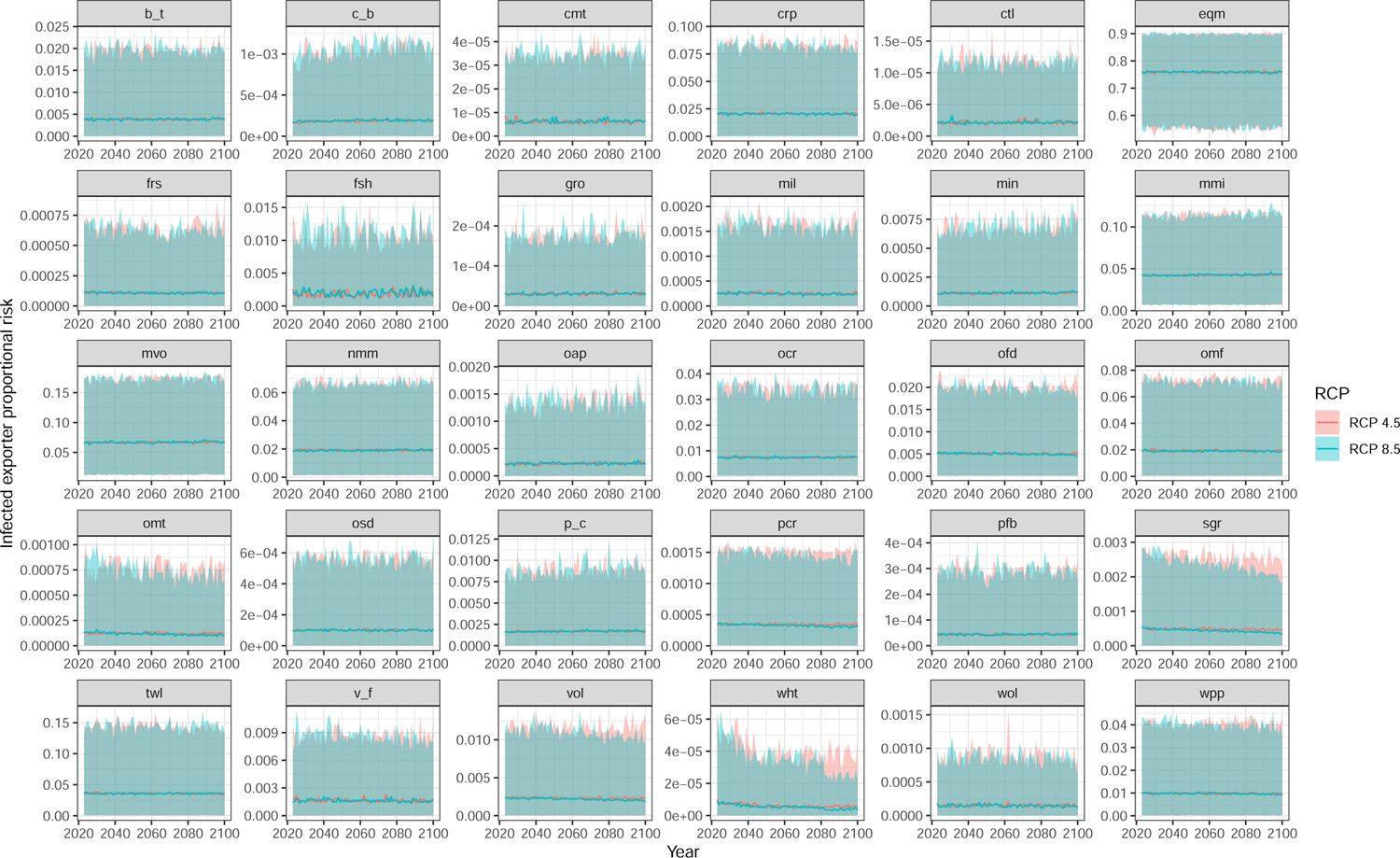
Annual mean (95% CI) commodity sector proportional contributions to Asian honey bee (*Apis cerana*) propagule pressure hitting Australia under RCP 4.5 and 8.5. Note: Predictions include stochastic uncertainty associated with future years and for unobserved commodity and infected exporter combinations.

## 10. Sheltering pest: Giant African snail (*Lissachatina fulica*)

### 10.1. Likelihoods of contamination

The giant African snail (*Lissachatina fulica*) contamination model revealed a weak positive effect of mean climate suitability on exporter contamination rates (Figure 10.1a). This implies that regions with higher average climate suitability tended to contain marginally higher contamination rates – albeit this effect was highly uncertain. The model also revealed that there was considerable variation within and between random grouping variables (Figure 10.1b). Specifically, contamination rates were most variable among exporters (Region SD = 2.91), followed by commodity sectors (Commodity SD = 1.92). The least amount of unexplained variation was among years (Year SD = 0.69).

**Figure 10.1.:**
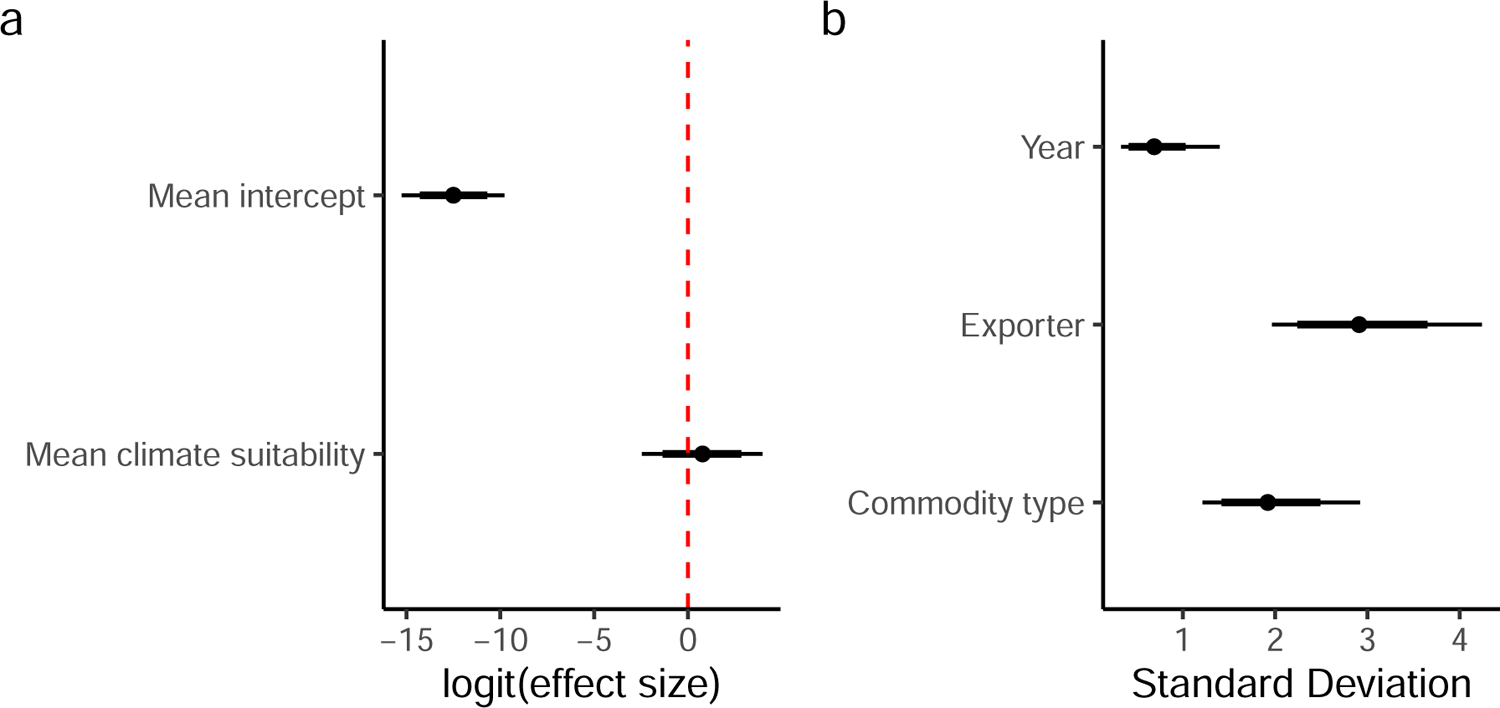
Giant african snail (*Lissachatina fulica*) coefficient and group variance plots. a) Model coefficients; b) Random effect standard deviations (i.e. measure of variability within each grouping variable). Error bars reflect 95% (thin) and 80% (thick) credible intervals.

Examination of baseline likelihoods for infected exporters (Figure 10.2a) revealed that Rest of Eastern Africa (xec) and Tanzania (tza) had on average the highest contamination rates. This was followed by the Rest of Oceania (xoc), Nigeria (nga), and the Rest of Southeast Asia (xse). By contrast, the United States (usa) had the lowest infected region contamination rate among infected regions.

Baseline likelihoods for commodity sector (Figure 10.2b) also showed that giant African snail contamination rates were highest in petroleum and coal products (p_c) and motor vehicles and associated parts (mvo). By contrast, processed rice (pcr) and sugar cane/beet (c_b) exhibited some of the lowest estimated contamination rates among modelled commodity sectors.

**Figure 10.2.:**
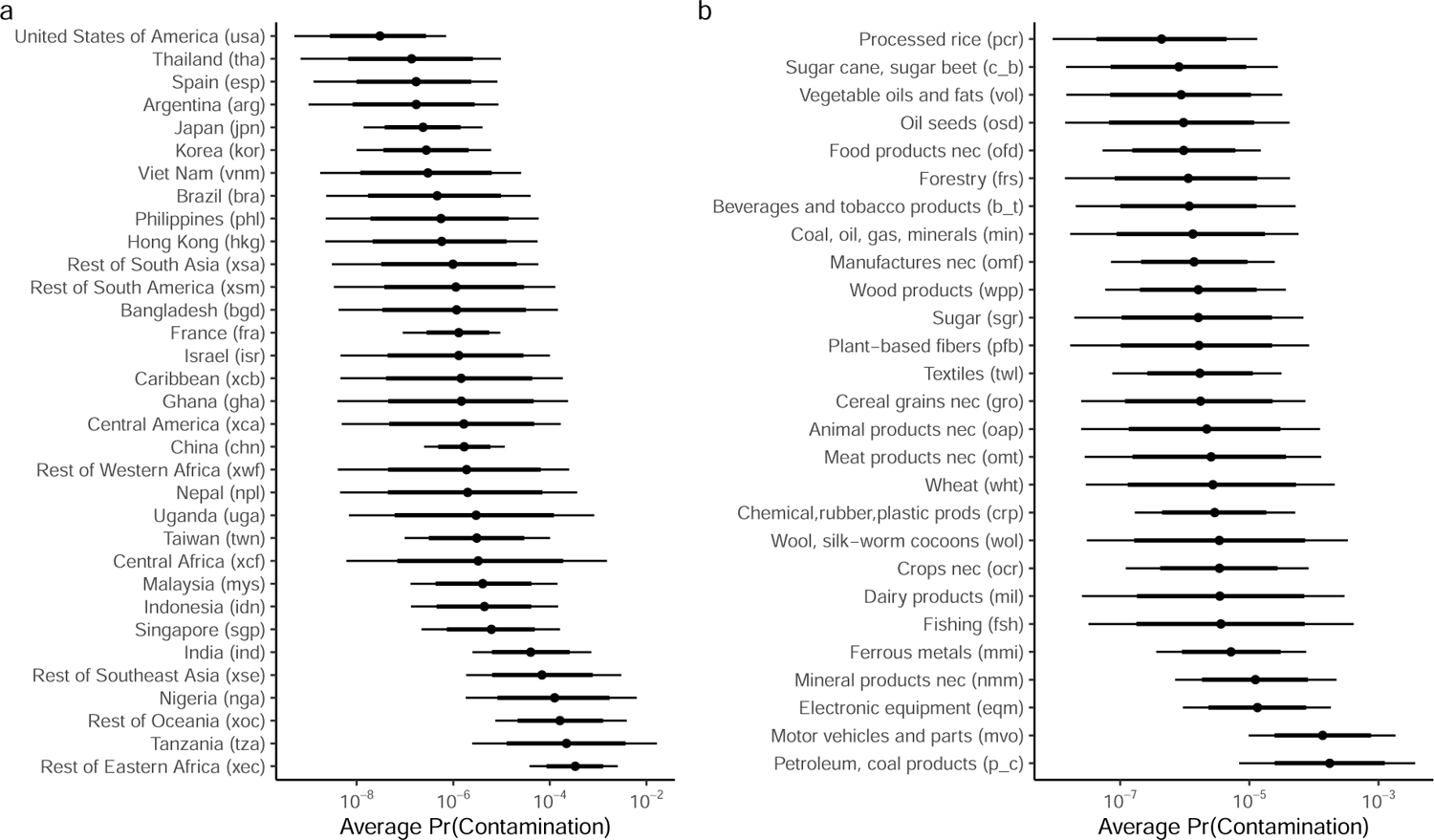
Giant african snail (*Lissachatina fulica*) baseline contamination likelihoods associated with (a) infected exporter; and (b) commodity type. Error bars reflect 95% (thin) and 80% (thick) credible invervals. Note: baseline likelihoods are back-transformed (anti-logit) intercept terms in the model and that do not account for exporter mean climate suitability. Giant African snail contamination rates between 2014 and 2021 were mostly stable, with marginally higher rates detected in 2019 (Figure 10.3).

Imports from Rest of Oceania (xoc) and Rest of southeast asia (xse) accounted for 11 of the top 20 estimated contamination likelihoods for 2023 (Table 10.1). In particular, imports of petroleum and coal products (p_c) from Rest of Oceania contained the highest estimated contamination likelihood with a 2.1% chance a single imported line was contaminated. This was followed by imports of motor vehicles and associated parts (mvo) from both xoc (1.2%) and Rest of Southeast Asia (xse; 0.76%). Other exporters that fell within the top 20 included: India (ind), Singapore (sgp), Indonesia (idn), Malaysia (mys) and Central America (xca). As climate suitability only had a weak (and highly uncertain) positive effect on contamination rates, likelihoods were predicted to only marginally change over time under both climate scenarios – with most exporting regions exhibiting declines in contamination rates by 2100 under both RCPs.

However, the climate suitability for this species is expected to change over time under both climate scenarios (Figure 6.5), with significant declines predicted for Central Africa and South America under both RCPs. This is likely to have long-term negative implications on the extent this species occupies in infected regions, and ultimately be have unmeasured implications on contamination rates from these regions (Table 10.2).

**Table 10.1.:** Giant African snail top 20 commodity sector by region contamination likelihoods for commodities with 100 or more imported lines into Australia and how they are predicted to change under RCP 4.5 and RCP 8.5 by 2050 and 2100. Numbers in brackets signify 95% credible limits.

| Exporter Commodity |  | 2023 | RCP 4.5 | RCP 4.5 | RCP 8.5 | RCP 8.5 |
| --- | --- | --- | --- | --- | --- | --- |
|  |  |  | 2050 | 2100 | 2050 | 2100 |
| xoc | p_c | 2.1e-02 (1.8e-03,8.1e-02) | 2.0e-02 (1.7e-03,7.9e-02) | 1.9e-02 (1.5e-03,7.8e-02) | 2.0e-02 (1.6e-03,8.4e-02) | 1.9e-02 (1.2e-03,8.5e-02) |
| xoc | mvo | 1.2e-02 (3.5e-03,3.5e-02) | 1.2e-02 (3.4e-03,3.4e-02) | 1.1e-02 (3.0e-03,3.4e-02) | 1.2e-02 (3.4e-03,3.4e-02) | 1.1e-02 (2.1e-03,3.6e-02) |
| xse | mvo | 7.6e-03 (4.8e-04,3.1e-02) | 7.4e-03 (4.6e-04,3.0e-02) | 7.1e-03 (4.2e-04,3.0e-02) | 7.3e-03 (4.7e-04,3.0e-02) | 6.8e-03 (3.4e-04,3.2e-02) |
| ind | p_c | 4.9e-03 (3.8e-04,2.0e-02) | 4.7e-03 (3.5e-04,2.0e-02) | 4.6e-03 (3.1e-04,2.0e-02) | 4.7e-03 (3.4e-04,2.0e-02) | 4.5e-03 (2.5e-04,2.0e-02) |
| ind | mvo | 2.9e-03 (7.8e-04,9.0e-03) | 2.8e-03 (7.1e-04,8.5e-03) | 2.7e-03 (6.1e-04,8.5e-03) | 2.7e-03 (7.1e-04,8.4e-03) | 2.6e-03 (4.5e-04,9.0e-03) |
| xoc | fsh | 2.7e-03 (4.5e-06,1.7e-02) | 2.6e-03 (4.3e-06,1.7e-02) | 2.5e-03 (4.1e-06,1.7e-02) | 2.5e-03 (4.8e-06,1.7e-02) | 2.4e-03 (3.5e-06,1.5e-02) |
| xec | nmm | 2.2e-03 (1.4e-04,9.3e-03) | 2.1e-03 (1.4e-04,9.3e-03) | 2.1e-03 (1.3e-04,9.3e-03) | 2.2e-03 (1.3e-04,9.0e-03) | 2.1e-03 (1.2e-04,8.8e-03) |
| xec | eqm | 2.1e-03 (2.0e-04,8.1e-03) | 2.1e-03 (1.9e-04,8.1e-03) | 2.1e-03 (1.9e-04,8.0e-03) | 2.1e-03 (1.8e-04,7.9e-03) | 2.0e-03 (1.7e-04,7.5e-03) |
| xoc | v_f | 1.7e-03 (3.9e-06,1.4e-02) | 1.6e-03 (3.8e-06,1.4e-02) | 1.6e-03 (3.7e-06,1.4e-02) | 2.0e-03 (4.1e-06,2.0e-02) | 1.9e-03 (2.8e-06,1.7e-02) |
| xoc | nmm | 1.4e-03 (1.8e-04,5.0e-03) | 1.3e-03 (1.7e-04,4.9e-03) | 1.3e-03 (1.5e-04,4.9e-03) | 1.3e-03 (1.5e-04,5.2e-03) | 1.3e-03 (1.0e-04,5.3e-03) |
| xoc | eqm | 1.3e-03 (3.3e-04,3.8e-03) | 1.2e-03 (3.2e-04,3.7e-03) | 1.2e-03 (2.8e-04,3.6e-03) | 1.2e-03 (3.0e-04,3.8e-03) | 1.1e-03 (1.9e-04,4.1e-03) |
| xse | v_f | 1.0e-03 (1.2e-06,9.0e-03) | 1.0e-03 (1.2e-06,8.8e-03) | 9.8e-04 (1.2e-06,8.5e-03) | 1.2e-03 (1.2e-06,1.0e-02) | 1.2e-03 (9.4e-07,9.3e-03) |
| sgp | p_c | 9.3e-04 (4.0e-05,4.2e-03) | 8.6e-04 (3.4e-05,4.1e-03) | 8.5e-04 (2.9e-05,4.1e-03) | 8.6e-04 (3.4e-05,4.2e-03) | 8.7e-04 (2.5e-05,4.5e-03) |
| xec | mmi | 8.5e-04 (7.4e-05,3.5e-03) | 8.4e-04 (7.3e-05,3.5e-03) | 8.4e-04 (7.2e-05,3.4e-03) | 8.4e-04 (7.2e-05,3.4e-03) | 8.1e-04 (6.9e-05,3.2e-03) |
| idn | p_c | 7.7e-04 (2.7e-05,3.8e-03) | 7.3e-04 (2.3e-05,3.6e-03) | 7.0e-04 (2.0e-05,3.6e-03) | 7.2e-04 (2.3e-05,3.7e-03) | 7.1e-04 (1.6e-05,3.9e-03) |
| xse | eqm | 7.7e-04 (4.6e-05,3.1e-03) | 7.5e-04 (4.3e-05,3.1e-03) | 7.2e-04 (4.0e-05,3.1e-03) | 7.4e-04 (4.5e-05,3.2e-03) | 6.9e-04 (3.3e-05,3.4e-03) |
| xec | ocr | 7.6e-04 (2.3e-05,3.8e-03) | 7.6e-04 (2.3e-05,3.7e-03) | 7.5e-04 (2.2e-05,3.7e-03) | 7.6e-04 (2.2e-05,3.8e-03) | 7.3e-04 (2.1e-05,3.7e-03) |
| xca | mvo | 6.5e-04 (1.8e-07,5.4e-03) | 6.5e-04 (1.8e-07,5.4e-03) | 6.5e-04 (1.8e-07,5.5e-03) | 6.6e-04 (1.8e-07,5.4e-03) | 6.6e-04 (1.8e-07,5.5e-03) |
| mys | p_c | 6.2e-04 (3.4e-05,2.7e-03) | 5.8e-04 (3.1e-05,2.6e-03) | 5.6e-04 (2.7e-05,2.5e-03) | 5.8e-04 (2.8e-05,2.6e-03) | 5.7e-04 (1.8e-05,3.0e-03) |
| xoc | oap | 6.2e-04 (3.0e-06,4.2e-03) | 6.0e-04 (3.0e-06,4.1e-03) | 5.7e-04 (2.7e-06,3.7e-03) | 5.8e-04 (2.9e-06,3.8e-03) | 5.5e-04 (2.6e-06,3.6e-03) |

**Table 10.2.:**
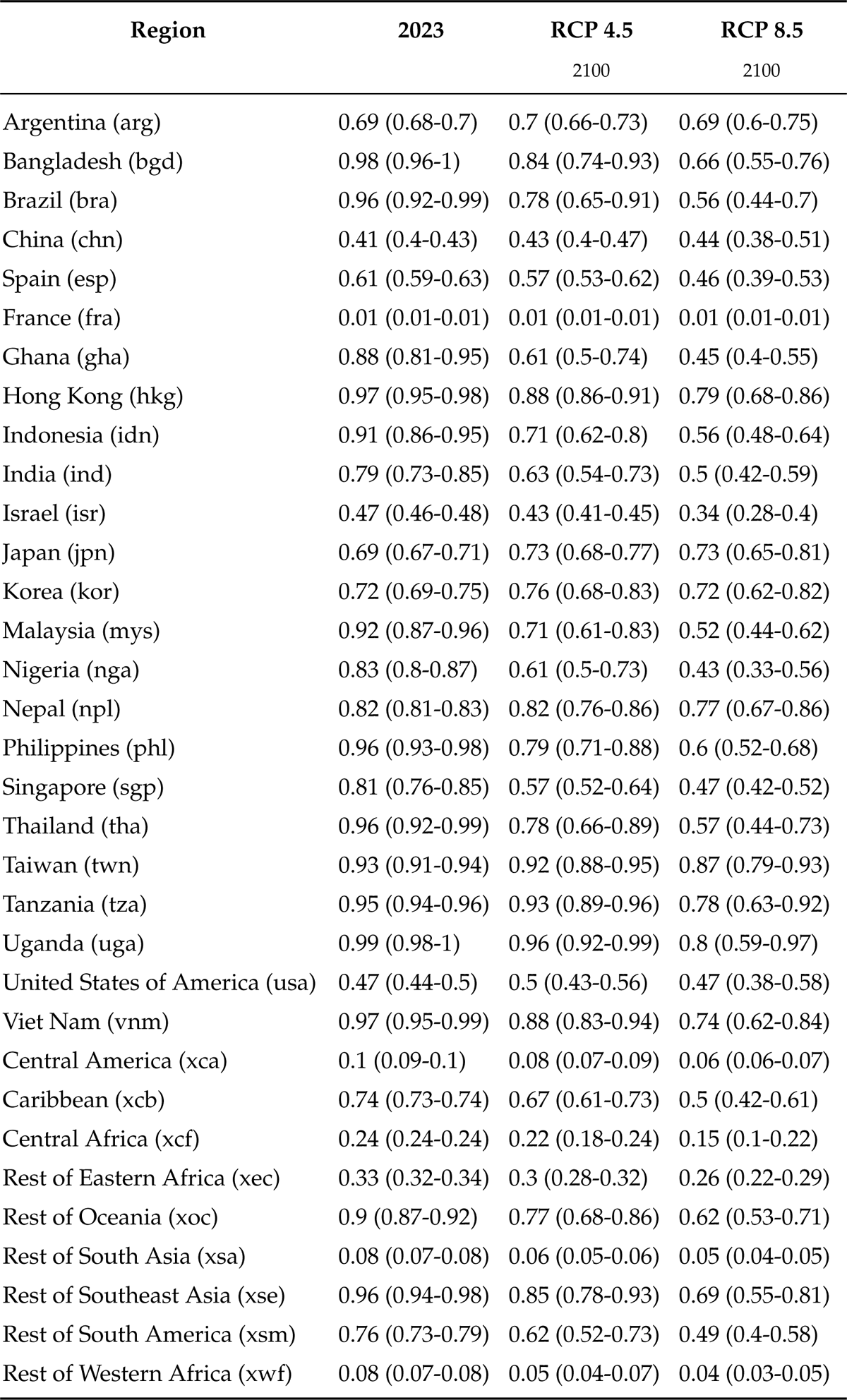
Giant african snail mean climate suitability for each infected exporter region in 2023 and 2100 for RCPs 4.5 and 8.5. Mean estimates are based on the ensembled annual climate suitabilities derived from 23 GCMs. Numbers in parentheses are the 10th and 90th percentiles derived from ensembles annual models.

### 10.2. Australia: Giant African snail propagule pressure and establishment exposure

Giant African snail (*Lissachatina fulica*) propagule pressure arriving at Australian borders is expected to marginally decline from 351 (95CI: 109-985) in 2023 to 319 (95CI: 91-953) and 302 (95CI: 71-1067) by 2100 for RCP 4.5 and RCP 8.5, respectively (Figure 10.4).

**Figure 10.3.:**
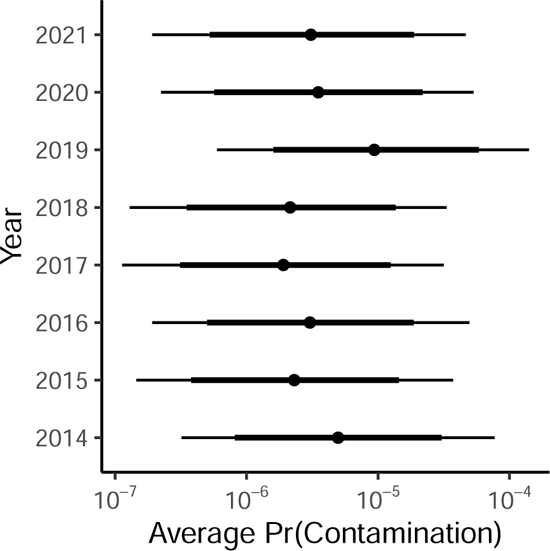
Giant african snail annual baseline contamination likelihoods. Error bars reflect 95% (thin) and 80% (thick) credible invervals. Note: baseline likelihoods are back-transformed (anti-logit) intercept terms in the model and that do not account for exporter mean climate suitability.

Estimated establishment exposure in Australia, is also expected to marginally decline under RCP 8.5 but remain relatively stable under RCP 4.5. This is because under RCP 8.5 there is a more significant decline in giant african snail suitable climatic conditions at locations where imported goods are expected to be unloaded (i.e., regions of highest human population density; Figure 6.5). As a consequence, the expected number of contaminated lines arriving at a climatically suitable locations within Australia will be lower.

**Figure 10.4.:**
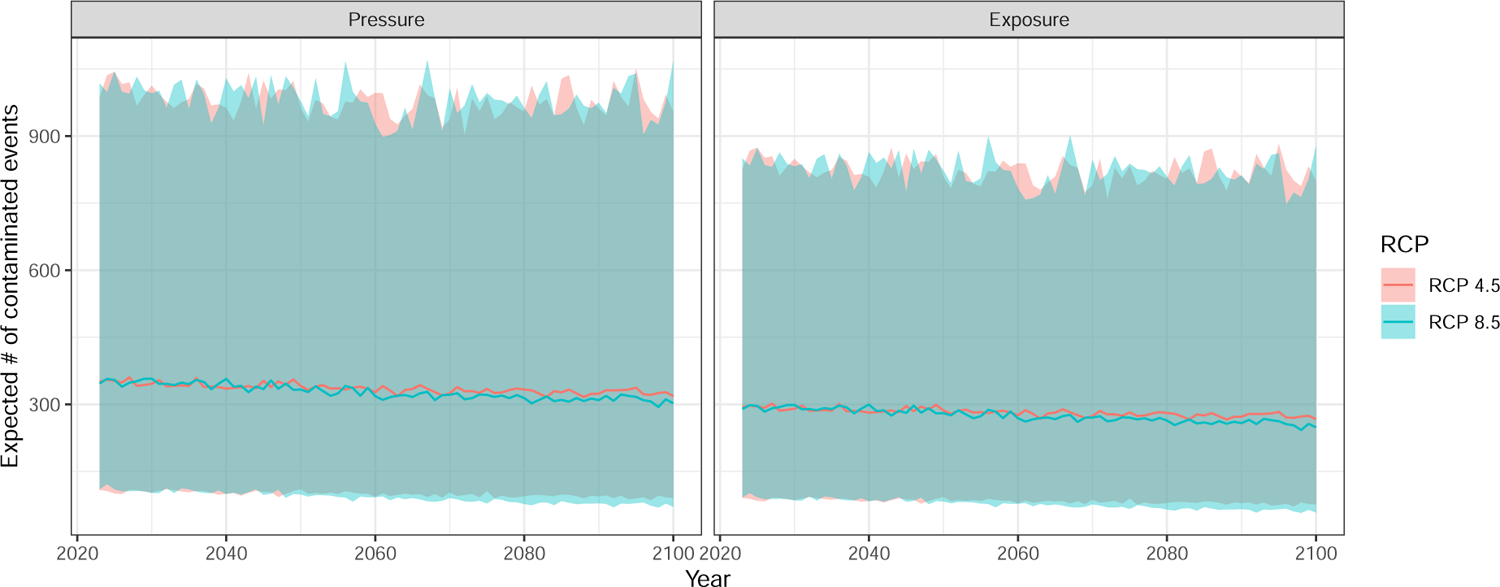
Annual mean (95% CI) giant African snail (*Lissachatina fulica*) propagule pressure (left) and establishment exposure (right) hitting Australia under RCP 4.5 and 8.5. Note: Predictions include stochastic uncertainty associated with future years and for unobserved commodity and infected exporter combinations.

### 10.3. Australia: Giant African snail risk attributable to different exporters & commodity sectors

The model indicates that India (ind) currently accounts for approxmately 40% (28- 53) of the total giant African snail propagule pressure hitting Australian borders (Figure 10.5). However, the proportion of risk attributable to India is expected to decline to 38% (24-53) and 35% (20-53) by 2100 for RCP 4.5 and 8.5, respectively. Other significant contributors to giant African snail propagule pressure in Australia were: China (chn; 14% (6-24)) and Taiwan (twn; 7% (1-19)), both of which are expected to become marginally greater contributors to risk under both future climate scenarios.

The risk attributable to different commodity sectors was also expected to remain relatively stable under both RCP scenarios (Figure 10.6). Imports of motor vehicles and associated parts (mvo) was the most significant contributor to giant African snail propagule pressure, accounting for approximately 47% (34-59) of the total number of predicted giant African snail contaminated lines entering Australia.

**Figure 10.5.:**
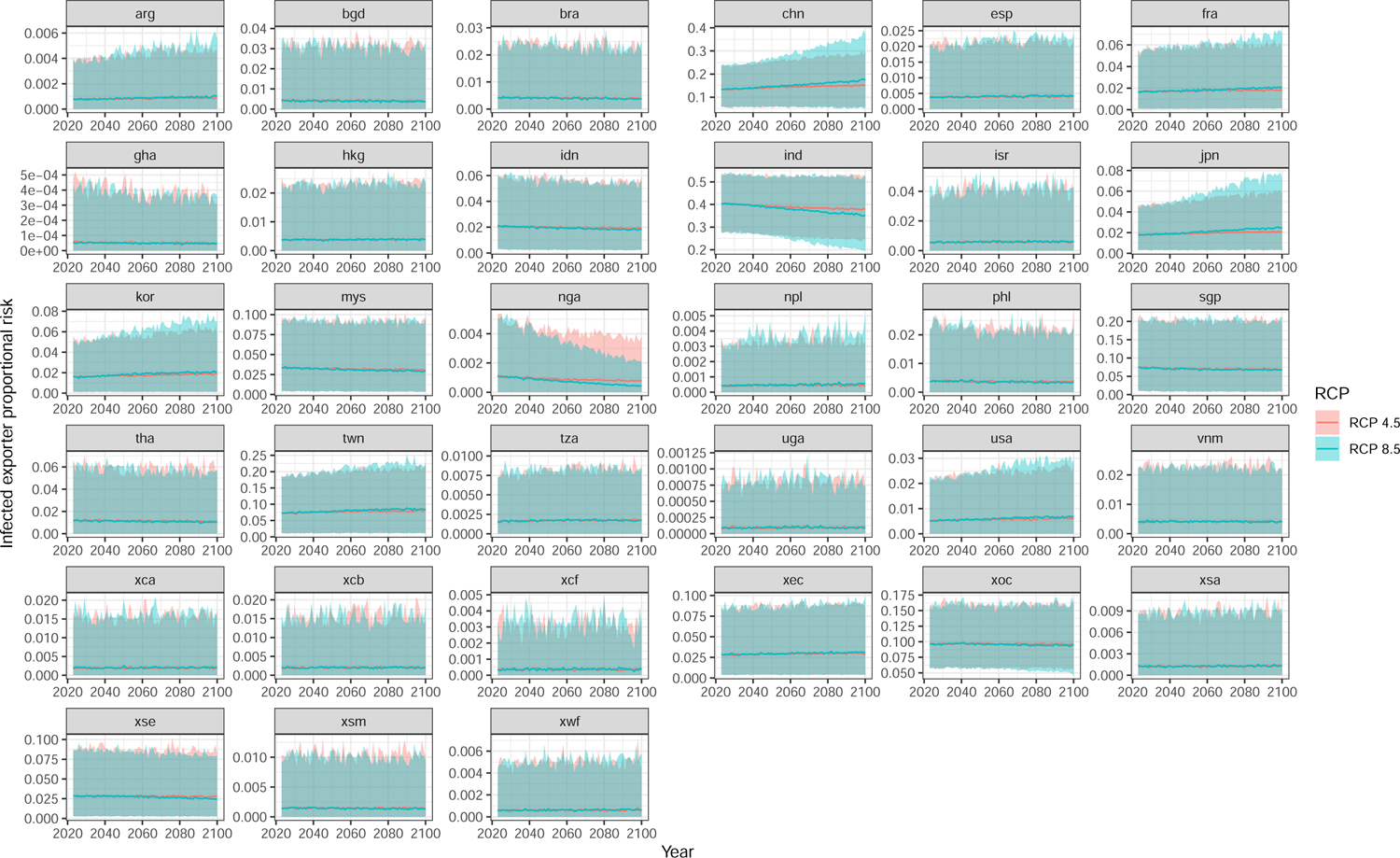
Annual mean (95% CI) exporter proportional contributions to giant African snail (*Lissachatina fulica*) propagule pressure hitting Australia under RCP 4.5 and 8.5. Note: Predictions include stochastic uncertainty associated with future years and for unobserved commodity and infected exporter combinations.

**Figure 10.6.:**
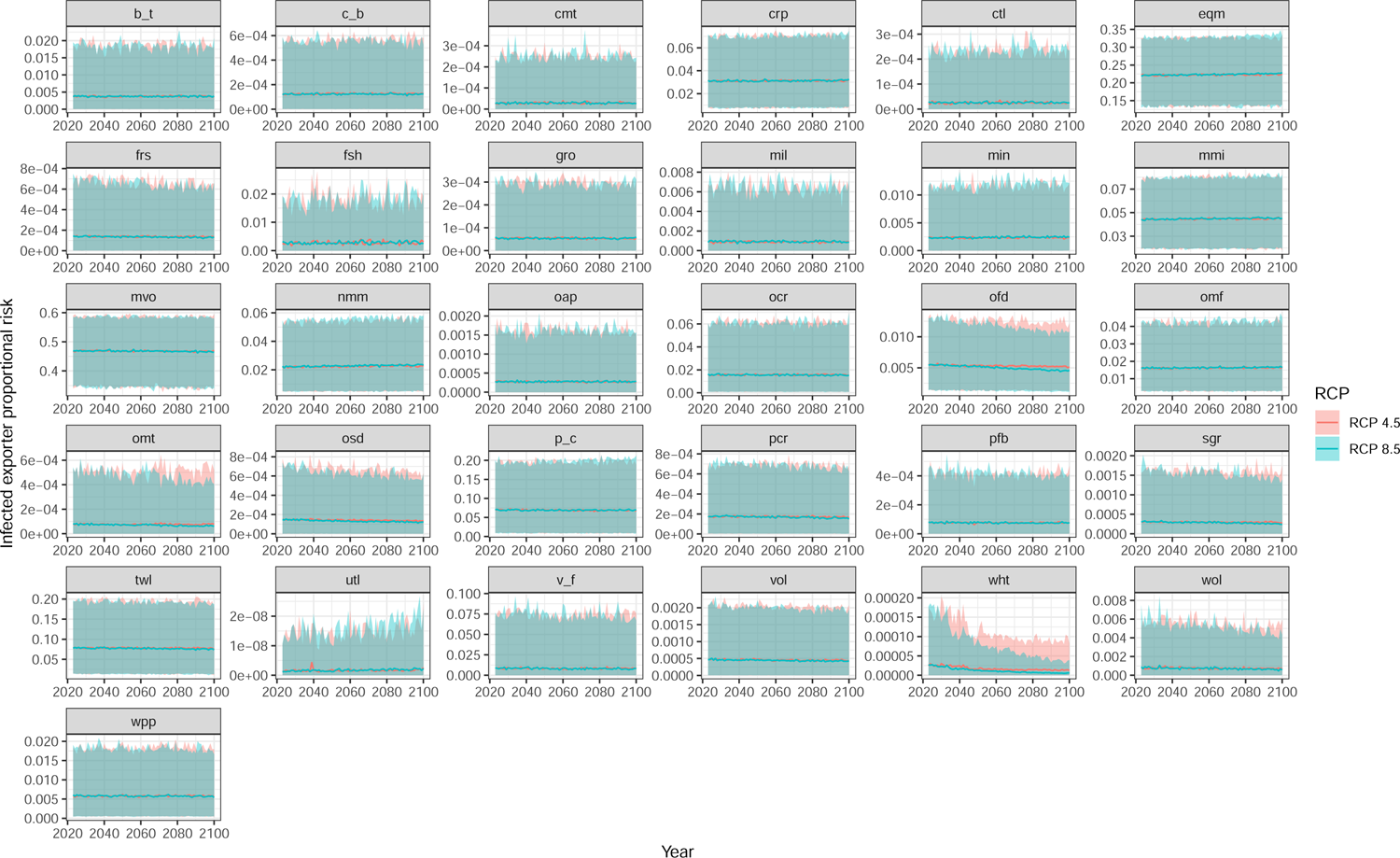
Annual mean (95% CI) commodity sector proportional contributions to giant African snail (*Lissachatina fulica*) propagule pressure hitting Australia under RCP 4.5 and 8.5. Note: Predictions include stochastic uncertainty associated with future years and for unobserved commodity and infected exporter combinations.

## 11. Internal storage pest: Khapra beetle (*Trogoderma granarium*)

### 11.1. Likelihoods of contamination

The Khapra beetle (*Trogoderma granarium*) contamination model revealed that there was considerable variation within each random grouping variable (Figure 11.1b). Specifically, estimated standard deviations in contamination rates among exporter (2.07), commodity sector (2.28) and year (2.37) were broadly similar – with year estimates being the most uncertain. Suggesting that there is similar variation among commodities, exporters, and years.

**Figure 11.1.:**
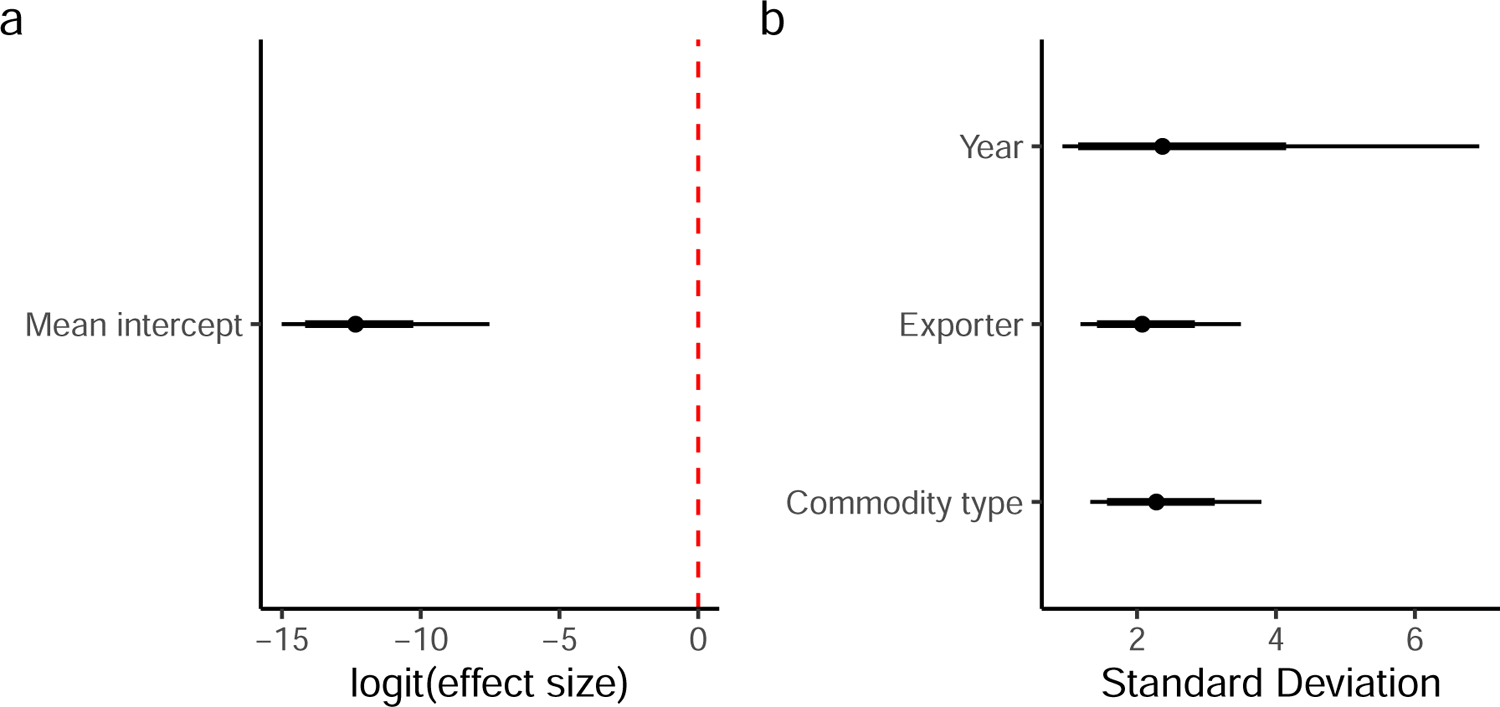
Khapra beetle (*Trogoderma granarium*) coefficient and group variance plots. a) Model coefficients; b) Random effect standard deviations (i.e. measure of variability within each grouping variable). Error bars reflect 95% (thin) and 80% (thick) credible intervals.

Examination of baseline likelihoods (Figure 11.2) revealed some variation in contamination rates across both exporters and commodity sectors. Specifically, the model indicated that Italy (`ita`), Pakistan (pak) and India (ind) had the highest average contamination rates, while Korea (kor) and Rest of the European Free Trade Association (`xef`) had some of the lowest estimated infected exporter rates. Similarly, there was also little variation among most commodity sectors. However, rates were higher for imports of motor vehicles and associated parts (mvo), processed rice (pcr), sugar (sgr), and food products not else classified (ofd).

It is important to note that Khapra beetle is not known to be established in mainland Italy. However, it is assumed to occur in Sardinia – an island region of Italy. Australian border screening data indicates that Khapra beetle is frequently detected on items originating from Italy. However, these detections are recorded to country of origin and not sub-regions within a country. Moreover, the detections attributed to Italy, may have originated from another source country. This is because Khapra is capable of persisting within containers for years, and thus, such an infected container may be used by different exporting countries prior to detection, making attribution of country of origin challenging. As such, modelled outputs for Khapra beetle should be interpreted with caution.

**Figure 11.2.:**
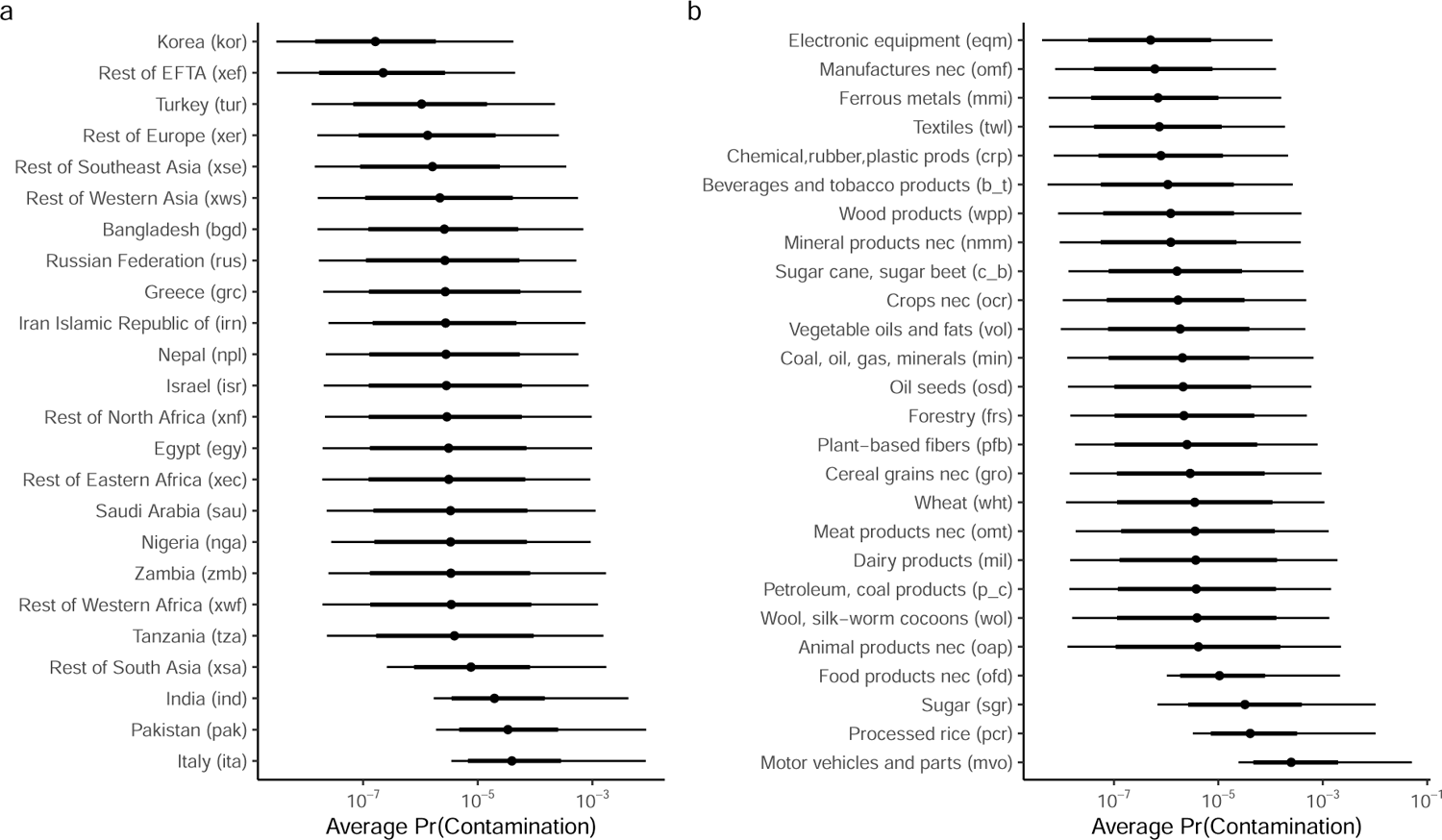
Khapra beetle (*Trogoderma granarium*) baseline contamination likelihoods associated with (a) infected exporter; and (b) commodity type. Error bars reflect 95% (thin) and 80% (thick) credible invervals. Note: baseline likelihoods are back-transformed (anti-logit) intercept terms in the model.

Imports of motor vehicles and associated parts (mvo) from infected exporters accounted for 10 of the top 20 estimated contamination likelihoods for 2023 (Table 11.1). Italy (`ita`) was also identified as the exporter contributing the most to the top 20, occupying 7 of the top 20 estimated contamination likelihoods (Table 11.1).

**Table 11.1.:** Khapra beetle top 20 commodity sector by region contamination likelihoods for commodities with 100 or more imported lines into Australia and how they are predicted to change under RCP 4.5 and RCP 8.5 by 2050 and 2100. Numbers in brackets signify 95% credible limits.

| Exporter Commodity |  | 2023 | RCP 4.5 | RCP 4.5 | RCP 8.5 | RCP 8.5 |
| --- | --- | --- | --- | --- | --- | --- |
|  |  |  | 2050 | 2100 | 2050 | 2100 |
| ita | mvo | 1.5e-03 (1.1e-05,8.2e-03) | 1.5e-03 (1.1e-05,8.2e-03) | 1.5e-03 (1.1e-05,8.2e-03) | 1.4e-03 (8.7e-06,8.2e-03) | 1.4e-03 (8.7e-06,8.2e-03) |
| ind | mvo | 7.9e-04 (5.2e-06,4.8e-03) | 7.9e-04 (5.2e-06,4.8e-03) | 7.9e-04 (5.2e-06,4.8e-03) | 7.3e-04 (4.2e-06,4.7e-03) | 7.3e-04 (4.2e-06,4.7e-03) |
| egy | mvo | 5.7e-04 (1.1e-07,4.6e-03) | 5.7e-04 (1.1e-07,4.6e-03) | 5.7e-04 (1.1e-07,4.6e-03) | 5.8e-04 (1.1e-07,4.4e-03) | 5.8e-04 (1.1e-07,4.4e-03) |
| xsa | mvo | 5.1e-04 (1.1e-06,4.0e-03) | 5.1e-04 (1.1e-06,4.0e-03) | 5.1e-04 (1.1e-06,4.0e-03) | 4.6e-04 (1.1e-06,3.6e-03) | 4.6e-04 (1.1e-06,3.6e-03) |
| xnf | mvo | 5.0e-04 (1.1e-07,4.3e-03) | 5.0e-04 (1.1e-07,4.3e-03) | 5.0e-04 (1.1e-07,4.3e-03) | 5.0e-04 (9.7e-08,3.7e-03) | 5.0e-04 (9.7e-08,3.7e-03) |
| rus | mvo | 4.5e-04 (9.8e-08,4.1e-03) | 4.5e-04 (9.8e-08,4.1e-03) | 4.5e-04 (9.8e-08,4.1e-03) | 3.5e-04 (1.2e-07,2.7e-03) | 3.5e-04 (1.2e-07,2.7e-03) |
| isr | mvo | 4.3e-04 (1.3e-07,3.5e-03) | 4.3e-04 (1.3e-07,3.5e-03) | 4.3e-04 (1.3e-07,3.5e-03) | 4.2e-04 (9.0e-08,3.5e-03) | 4.2e-04 (9.0e-08,3.5e-03) |
| xws | mvo | 3.0e-04 (8.7e-08,2.4e-03) | 3.0e-04 (8.7e-08,2.4e-03) | 3.0e-04 (8.7e-08,2.4e-03) | 2.8e-04 (1.2e-07,2.3e-03) | 2.8e-04 (1.2e-07,2.3e-03) |
| ita | pcr | 2.8e-04 (1.5e-06,1.9e-03) | 2.8e-04 (1.5e-06,1.9e-03) | 2.8e-04 (1.5e-06,1.9e-03) | 2.6e-04 (1.3e-06,1.8e-03) | 2.6e-04 (1.3e-06,1.8e-03) |
| pak | pcr | 2.6e-04 (1.1e-06,1.9e-03) | 2.6e-04 (1.1e-06,1.9e-03) | 2.6e-04 (1.1e-06,1.9e-03) | 2.4e-04 (9.6e-07,1.8e-03) | 2.4e-04 (9.6e-07,1.8e-03) |
| ita | p_c | 2.6e-04 (1.1e-08,1.8e-03) | 2.6e-04 (1.1e-08,1.8e-03) | 2.6e-04 (1.1e-08,1.8e-03) | 2.9e-04 (8.5e-09,1.7e-03) | 2.9e-04 (8.5e-09,1.7e-03) |
| ita | wol | 2.6e-04 (1.6e-08,2.0e-03) | 2.6e-04 (1.6e-08,2.0e-03) | 2.6e-04 (1.6e-08,2.0e-03) | 2.1e-04 (1.3e-08,1.3e-03) | 2.1e-04 (1.3e-08,1.3e-03) |
| ita | mil | 2.1e-04 (1.2e-08,1.3e-03) | 2.1e-04 (1.2e-08,1.3e-03) | 2.1e-04 (1.2e-08,1.3e-03) | 1.9e-04 (9.8e-09,1.3e-03) | 1.9e-04 (9.8e-09,1.3e-03) |
| ita | omt | 2.0e-04 (1.3e-08,1.4e-03) | 2.0e-04 (1.3e-08,1.4e-03) | 2.0e-04 (1.3e-08,1.4e-03) | 1.7e-04 (1.4e-08,1.2e-03) | 1.7e-04 (1.4e-08,1.2e-03) |
| xse | mvo | 1.9e-04 (8.0e-08,1.5e-03) | 1.9e-04 (8.0e-08,1.5e-03) | 1.9e-04 (8.0e-08,1.5e-03) | 1.8e-04 (6.3e-08,1.5e-03) | 1.8e-04 (6.3e-08,1.5e-03) |
| ind | sgr | 1.8e-04 (3.0e-07,1.5e-03) | 1.8e-04 (3.0e-07,1.5e-03) | 1.8e-04 (3.0e-07,1.5e-03) | 1.8e-04 (2.5e-07,1.3e-03) | 1.8e-04 (2.5e-07,1.3e-03) |
| ita | v_f | 1.5e-04 (1.3e-08,1.1e-03) | 1.5e-04 (1.3e-08,1.1e-03) | 1.5e-04 (1.3e-08,1.1e-03) | 1.5e-04 (1.1e-08,1.2e-03) | 1.5e-04 (1.1e-08,1.2e-03) |
| pak | v_f | 1.4e-04 (9.4e-09,9.3e-04) | 1.4e-04 (9.4e-09,9.3e-04) | 1.4e-04 (9.4e-09,9.3e-04) | 1.7e-04 (6.7e-09,1.1e-03) | 1.7e-04 (6.7e-09,1.1e-03) |
| ind | pcr | 1.3e-04 (7.9e-07,8.1e-04) | 1.3e-04 (7.9e-07,8.1e-04) | 1.3e-04 (7.9e-07,8.1e-04) | 1.2e-04 (7.4e-07,8.1e-04) | 1.2e-04 (7.4e-07,8.1e-04) |
| xer | mvo | 1.3e-04 (7.4e-08,1.1e-03) | 1.3e-04 (7.4e-08,1.1e-03) | 1.3e-04 (7.4e-08,1.1e-03) | 1.2e-04 (6.9e-08,1.0e-03) | 1.2e-04 (6.9e-08,1.0e-03) |

Note that because climate suitability was not included as a predictor of contamination rates for this species^1^, changes in contamination likelihoods over time and under different RCPs were minimal (Table 11.1).

### 11.2. Australia: Khapra beetle propagule pressure

Khapra beetle propagule pressure arriving at Australian borders is not expected the change substantially from contemporary estimates of approximately 169 (95CI: 1-923) contaminated events per year (Figure 11.3).

**Figure 11.3.:**
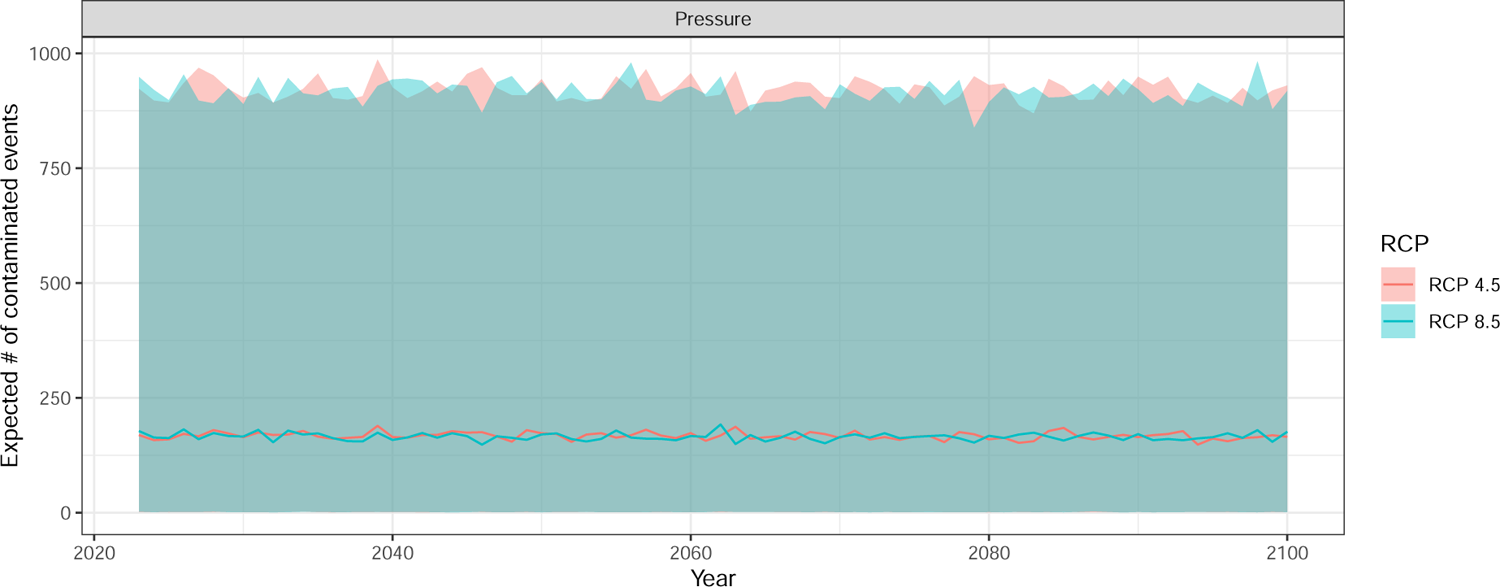
Annual mean (95% CI) Khapra beetle (*Trogoderma granarium*) propagule pressure hitting Australia under RCP 4.5 and 8.5. Note: Predictions include stochastic uncertainty associated with future years and for unobserved commodity and infected exporter combinations.

### 11.3. Australia: Khapra beetle risk attributable to different exporters & commodity sectors

The model indicates that Italy (`ita`) accounts for approximately 77% (63-87) of the total Khapra beetle propagule pressure hitting Australian borders (Figure 11.4). This proportion of risk is also expected to marginally increase under RCP 8.5 to 79% (67-88) by 2100. This increase is associated with expected increases in imports of high risk commodity types such as motor vehicles (mvo; Figure 5.2) and processed rice (pcr; Figure 5.3). The other major contributor to Khapra beetle propagule pressure in Australia was India (ind). However, this contribution is expected to marginally decline under RCP 8.5 from 17% (8-28)) to 14% (7-26)) by 2100.

The risk attributable to different commodity sectors was predicted to remain relatively stable under both RCP scenarios (Figure 11.5). Imports of motor vehicles and associated parts (mvo) was the greatest contributor to Khapra beetle propagule pressure, accounting for approximately 83% (63-93) of the total number of predicted Khapra beetle contaminated lines entering Australia.

**Figure 11.4.:**
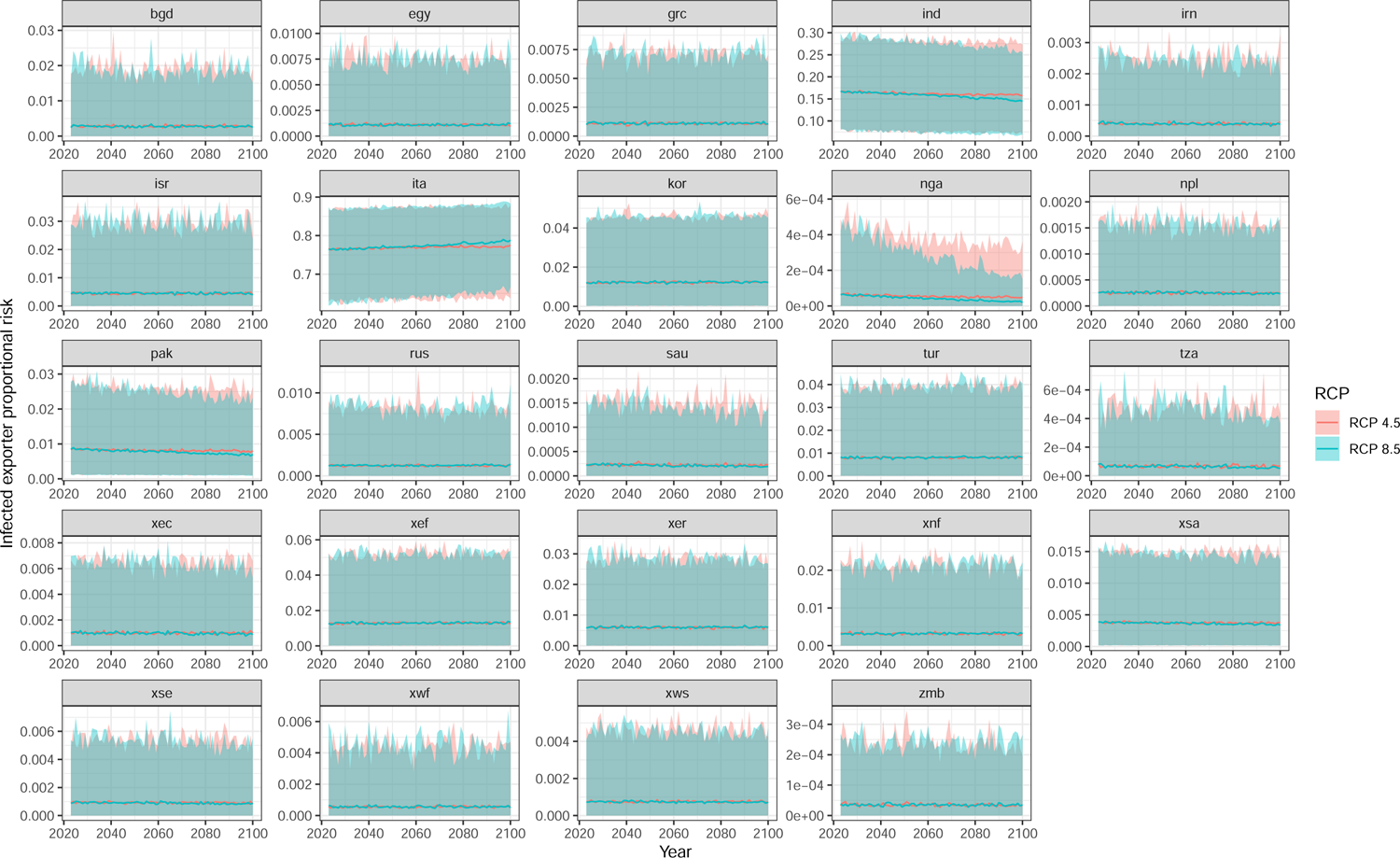
Annual mean (95% CI) exporter proportional contributions to Khapra beetle (*Trogoderma granarium*) propagule pressure hitting Australia under RCP 4.5 and 8.5. Note: Predictions include stochastic uncertainty associated with future years and for unobserved commodity and infected exporter combinations.

**Figure 11.5.:**
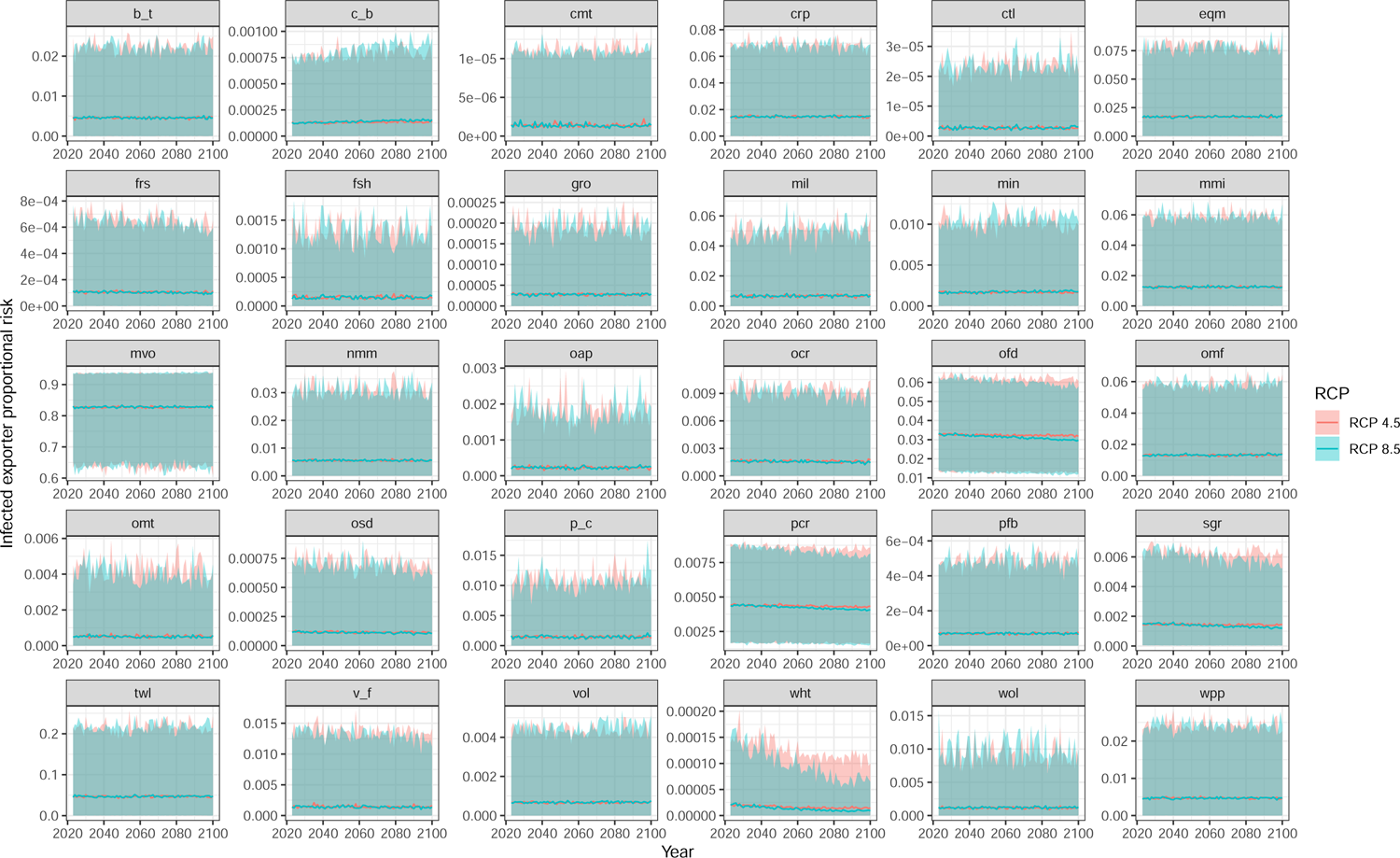
Annual mean (95% CI) commodity sector proportional contributions to Khapra beetle (*Trogoderma granarium*) propagule pressure hitting Australia under RCP 4.5 and 8.5. Note: Predictions include stochastic uncertainty associated with future years and for unobserved commodity and infected exporter combinations.

## 12. Future opportunities

There are substantial opportunities to improve and extend the models. Currently, the CGE imposes climate related sectorial damages based on a limited set of damage functions, notably: limited loss in areable land due to sea-level rise, temperature-related crop productivity losses and heat stress losses on agricultural and labour productivity. However, a rapidly changing climate is expected to have a wide range of direct and indirect impacts that are not yet accounted for within the CGE model. As a consequence, the estimated GDP damages and impacts on trade and biosecurity risk are likely to be underestimated. To better account for these other damages, the CGE model can be improved by incorporating additional damage functions such as climate impacts on water stress, sea level rise and storm surge, and the risk of extreme events (e.g. bushfires and inland flooding). The CGE model can also be improved by incorporating long-term socioeconomic trajectories which have the potential to substantially alter the composition of Australian imports and exports. These trajectories are likely to exert as large or larger impacts than climate change on trade patterns over the rest of the century, particularly in regions undergoing demographic decline. Future work will hence seek to incorporate more comprehensive climate change damages *as well as* long-term socioeconomic trends.

Similarly, the contamination model can be improved and extended through the inclusion of additional border interception data from other countries such as New Zealand and the addition of other predictor variables (e.g. region specific estimates of threat area of occupancy or duration pest has been present). Doing so, will improve estimates (and associated uncertainty) of country by commodity contamination rates observed across a range of different regions. Lastly, the contamination model may also be extended from a threat-specific model to a multi-threat model, increasing the utility of the model by allowing it to make predictions of contamination rates for new or emerging threats. This could be achieved by either undertaking a functional group analysis, or by directly incorporating covariate information on species morphological (e.g. size, wing-span) and physiological (e.g. temperature tolerance, generation time, and fecundity) traits that may contribute to hitch-hiking risk. In the sections below, we briefly describe these (and other) opportunties to extend and improve the model.

### Improve trade model damage functions

Although economic damages from global warming in this report are considerable, the damage functions used are still very limited, concentrating only on losses in agricultural and labour productivity from heat stress and limited sea level rise in terms of losses in arable land. This can be improved with more elaborate damage functions and further calibration of the impacts of heat stress. For example, economic losses from floods and fires, air pollution, and damages to biodiversity need to be included. Major impacts will also accrue from sea level rise and storm surge, likely swamping the impact of heat stress for many countries near the coast or in coastal regions. Indeed, countries with manufacturing facilities and hubs near coastlines may be disproportionately impacted by sea-level rise. Finally, the impacts of water stress to both green and blue water sources needs to be calibrated. Blue water is the primary source for water for irrigated agriculture, impacting not only productivity but food security concerns. The team at CEBRA/CEER has already begun some work on sea level rise and storm surge, along with the impacts of blue water stress, but much more primary research remains.

### Simulate socioeconomic pathways

The scenarios employed within this analysis simulate the impact of some aspects of climate change on selected industries through a long-term time horizon. Within a horizon extending to 2100 major socioeconomic shifts in demography and GDP are expected to occur. Many such shifts (e.g., demographic transitions to low fertility) occur heterogeneously and will alter trade and production trends at a magnitude as large as or greater than climate change. Scenarios such as the Shared Socioeconomic Pathways (Riahi *et al*., 2017) provide qualitative storylines and quantitative trajectories to examine potential socioeconomic futures. The team at CEBRA/CEER is in an ideal position to model these pathways in addition to climate change damages in order to provide a more comprehensive outlook for long-term Australian trade relations. Quantitative trajectories for such scenarios in terms of GDP (Dellink *et al*., 2017) and population (Commission *et al*., 2018; Kc, 2020) have already been simulated within a recursive-dynamic model at a smaller aggregation than the model used in this study (Cantele, 2023). Within the context of the current analysis, incorporation of socioeconomic pathways will compound economic losses for select regions while faster growing economies may to a limited extent offset damages attributable to climate change.

### Australia-China bilateral trade

For example, Chinese fertility levels have remained below the 2.1 replacement level for nearly three decades and the current population level is widely regarded as peak. Regardless of ongoing initiatives, a precipitous decline in population is already baked into current projections (see Figure 12.1). As the largest exporter to Australia, the impact of a declining Chinese population has implications for production within labour supply sensitive economic sectors of growth. Falling private household demand in China is also likely to indirectly impact global trade flows. Under such a future environment there is the potential for key imports to Australia such as manufacturing goods to be displaced to other regions (e.g., Southeast Asia). It is worth reiterating here that neither physical capital depreciation due to coastal inundation nor the cost of moving manufacturing away from vulnerable coastal areas is simulated in the current model. This is particularly relevant given the potential impacts of sea level rise on the Chinese eastern seaboard as well as the whole of Southeast Asia.

**Figure 12.1.:**
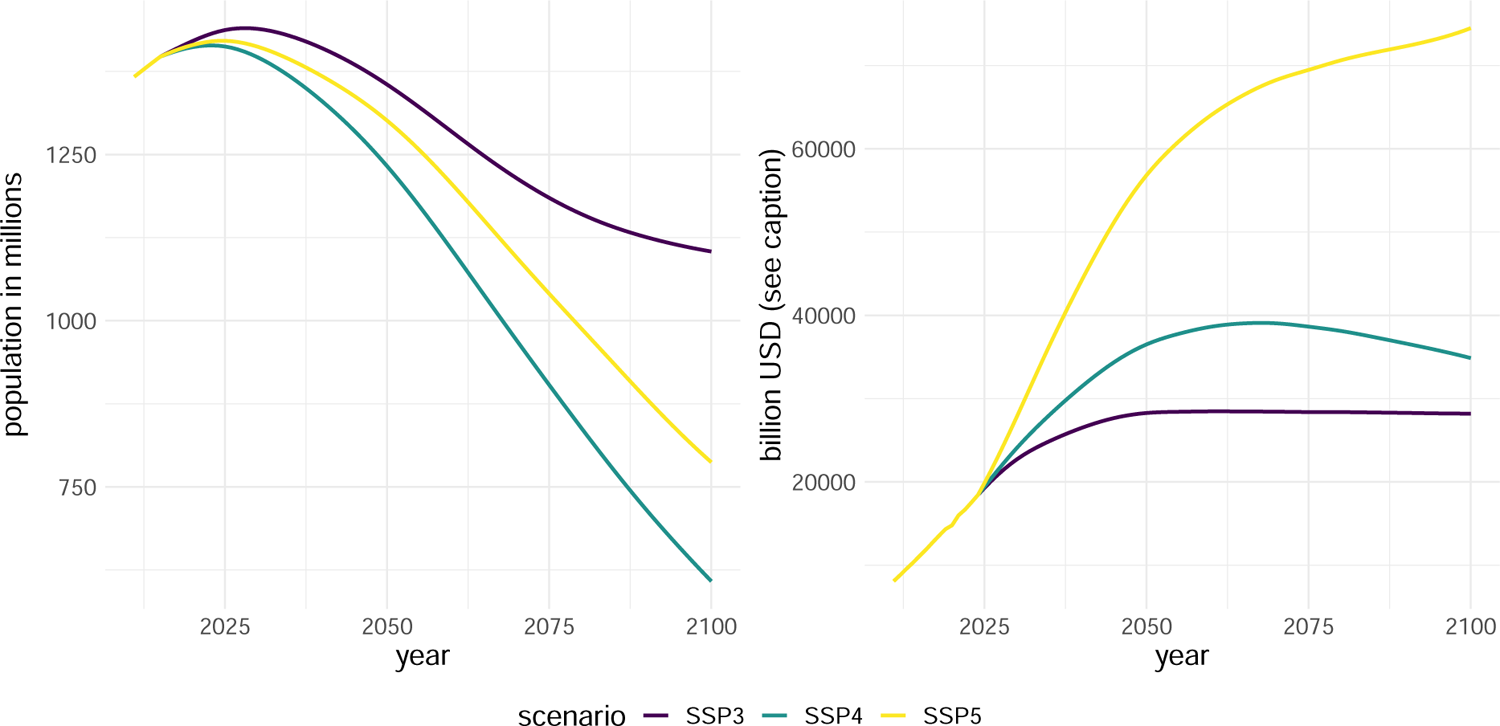
Demographic and GDP trajectories for China under SSP3, SSP4, and SSP5. Demographic data from Lutz et al. (2018) and Kc (2020). GDP data from World Bank WDI and Global Economic Prospects through 2024 - author calculations from 2024 (Cantele 2023), denominated in constant 2015USD.

### Account for contamination viability & leakage rates

Currently, the model estimates the likelihood of an imported line being contaminated with a given threat, which in turn, is multiplied by the expected number of lines imported into a country to derive an estimate of propagule pressure hitting a region’s border. Such estimates are almost certaintly an over-estimate of the true propagule pressure. This is because in most cases, there is a low likelihood that an imported line contains a viable population that survives the journey and is large enough to result in a potential establishment. To account for such viability would require detailed data on the number of surviving individuals detected at the border per contamination incident for each region by commodity combination. Incorporating such an effect may become achievable if additional data is collected at the border (e.g., numbers of surviving individuals rather than just presence/absence). An interim alternative could be to use expert elicitation to estimate the proportion of region by commodity contaminated events likely to be viable.

Similarly, the model also does not account for the proportion of contaminated imports that may “leak” through border inspections. To incorporate such an effect would require an understanding of each region’s border screening effectiveness & effort. This is likely to vary considerably among commodity types, modes of transport and across regions. Ideally, such estimates would be derived through independent and random repeated sampling of inspected goods. Alternatively, expert elicitation can also be used to provide estimates.

While the inclusion of both leakage and viability rates will reduce the over-estimation of region-specific propagule pressure and establishment exposure. Obtaining such estimates (whether by data or expert elicitation) for all 70 regions will likely be impractical. As such, alternative methods or assumptions should be considered when incorporating these processes.

### Simulate threat global spread

The current model estimates threat propagule pressure and establishment exposure based on exporters known to have established populations of the threat. The model does not currently simulate spread into unoccupied regions. This however is feasible as the model estimates both the propagule pressure hitting a region’s border as a function of trade with known infected regions, but also approximates the number of propagules likely to arrive at a climatically suitable location. By extending the model to forecast spread, one could approximate potential regional damages to various asset classes as well as assess what marginal benefits can be obtained from different border and pre-border interventions.

### Multi-species contamination models

The current pest-trade model is based on threat-specific contamination likelihoods and climate suitability layers. However, there is potential to make this model multi-threat focused by building multi-species contamination models. This could be achieved through the use of species traits (Palma *et al*., 2021) coupled with species random effect terms in the contamination model. Such models could be developed for various hitchhiking functional groups (e.g., flies, spiders, bees, ants, moths, bugs, and snails) and provide great advantage in predicting risk associated with new and emerging threats, where interception data is absent.

### Extend trade and contamination models to account for tourist movements

It is important to note that this model only considers trade flows, and thus, propagule pressure, associated with trade. Currently the model does not account for air passenger movement, illegal trade, natural spread, or other pathways of entry. Modelling natural spread is likely to be a difficult exercise as it requires precise GIS locations of known established populations coupled with accurate estimates of dispersal kernels – data that may not be easily sourced from all countries. By contrast, estimating the propagule pressure related to air passenger movements may be achievable through the combination of passenger inspection data and global air passenger movement data, which is increasingly being made available. It is possible to extend the GTAP framework to also forecast changes in tourist movements, which are already projected to be substantial (e.g., tourist arrivals to Australia are estimated to fall considerably under RCP 8.5 with losses to the Great Barrier Reef and excessive heat and flooding in the eastern States). Doing so would allow the model to account for another major pathway of pest and disease entry.

### Reduce uncertainty by adding predictors and additional data to contamination model

As seen in model outputs, some predictions are highly uncertain. Some of this uncertainty could be better accounted for if we were to add additional predictor variables to the contamination model. Additional predictors at the regional level (e.g. area pest occupies, or time the pest has been present), commodity level (e.g. covariates related to commodity susceptibility) could greatly reduce model uncertainty.

Model uncertainty could also be reduced by adding additional border interception data from other countries. New Zealand border interception data would be the logical first step, as they have similar standards of biosecurity screening at the border.

### Model validation

Models such as the one proposed in this report are notoriously difficult to validate due to the nature of the problem it attempts to estimate – rare events – coupled with there being very few independent datasets available to accurately test model predictive performance. One possibility would be to examine how well current and historical temporal patterns of establishment align with current and hind-cast predictions from the model. This validation exercise would bring additional challenges in that it requires:

1. estimates of changes in national income of GDP globally and by country as a result of already apparent global warming impacts;
2. validation of baseline line GDP measures and commodity sector shares by alternative (past) baselines;
3. the pest/disease to have only spread among countries predominately as a function of international trade or in a way that is correlated with international trade;
4. a detailed understanding of the time line of incursions and which country they originated from;
5. estimates of how interception/contamination rates have changed among countries and tariff types over time;
6. an understanding of how effective each countries biosecurity system is at mitigating the pest entry and establishment; and
7. historical estimates of trade flows among countries.

Compiling a dataset that would meet all of the above conditions to accurately test the model’s predictive performance would be a substantial undertaking. However, with the increased use of DNA analyses coupled with more sophisticated collation of border surveillance statistics among World Trade Organisation member countries such a dataset may be much easier to produce in future.

The contamination model sub-model could be partially validated by testing how well it can predict new Australian border screening data (e.g. data from 2022 to 2024). Alternatively, it could assessed by comparing its predictions on region propagule pressure with border interception data of other countries. However, using such border interception data from other countries, and the counts derived from it, will be strongly dependent on the border inspection effort and sensitivity. As such, such comparisons should only be done with regions that have well developed border inspection methods and estimates of screening effort (i.e., number of lines inspected vs number of lines imported).

Another approach is to fit the model using a different countries interception data and assess whether the relative rankings of exporter and commodity risk is the same. But again, this approach will be highly dependent on the sensitivity of the screening approaches and the surveillance effort imposed.

An easier task is to test recent trade flows in terms of the pattern of imports and exports between countries to determine if they are consistent with GTAP data. Our preliminary investigation indicates that the GTAP database is consistent in this sense but more work could be done here.

## Acknowledgements

This report is a product of the Centre of Excellence for Biosecurity Risk Analysis (C-EBRA). In preparing this report, the authors acknowledge the financial and other support provided by the Australian Department of Agriculture, Fisheries and Forestry and its predecessors, the New Zealand Ministry for Primary Industries and the University of Melbourne. We thank Dr John Baumgartner for providing programming advice and Estibaliz Palma who contributed to the development of conceptual diagrams. We also provide special thanks to the project sponsor Peter Gooday and department collaborators: Shalan Scholfield, Justin Billings, Tony Arthur, Brian Garms, Bethany Stone, Rachel Slatyer and Chris Wark. Lastly, we thank the two external reviewers and two departmental reviewers of this report, your feedback was invaluable.

## A. Posterior predictive checks for BMSB

**Figure A.1.:**
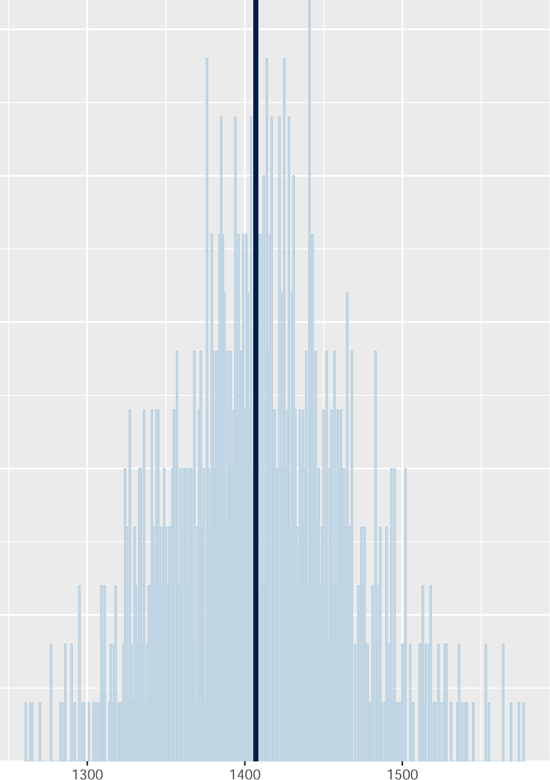
Observed vs predicted total numbers of BMSB contaminated lines across all infected exporters, commodities and years. Observed counts shown as dark blue line. Samples of posterior predictive distribution are shown as light blue bars. Note: Observed and predicted counts are based on realised number of lines inspected. This will be an underestimate of the true contamination count as not all lines are inspected.

**Figure A.2.:**
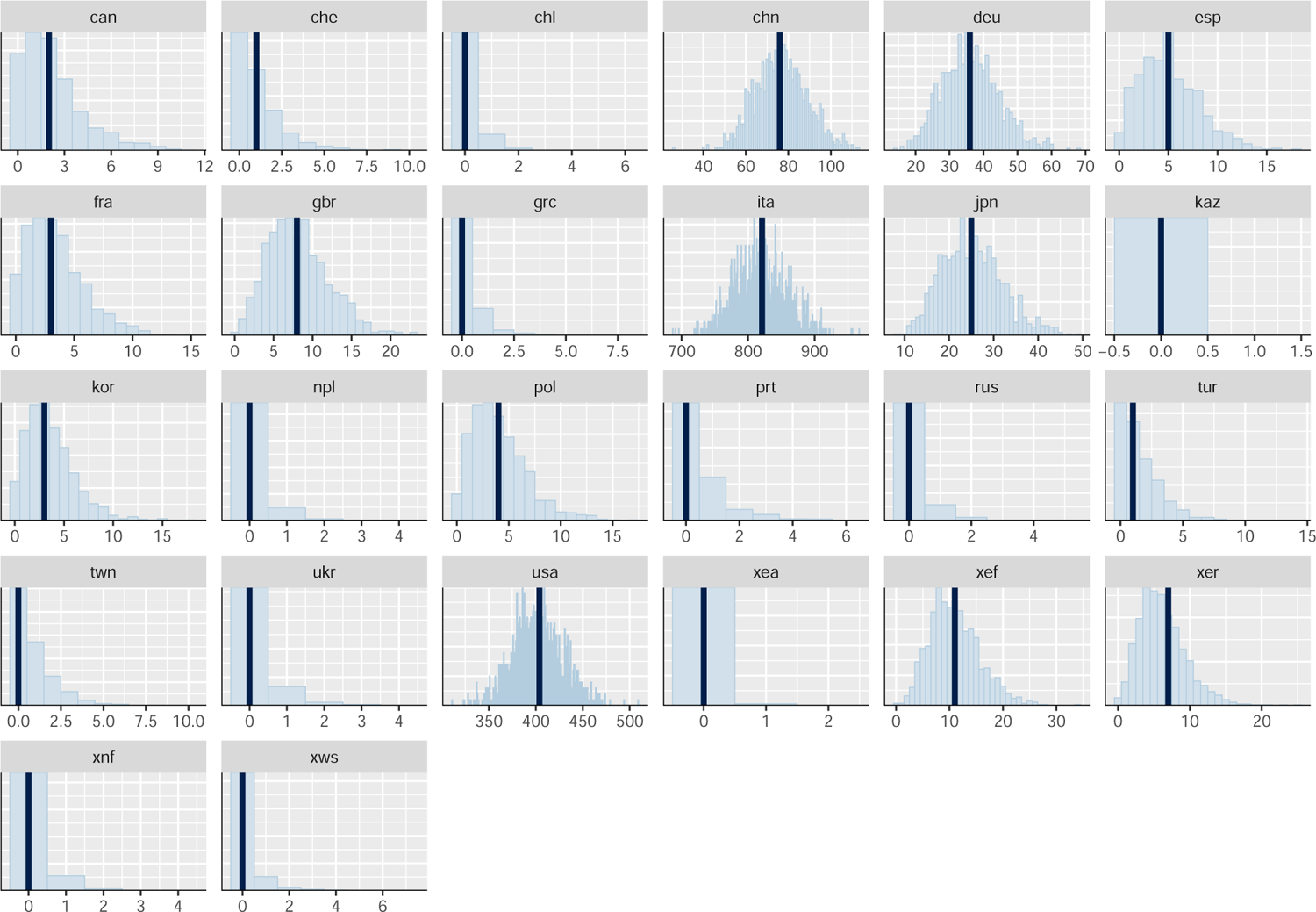
Observed vs predicted total numbers of BMSB contaminated lines across all years and commodity types for each infected exporter. Observed counts shown as dark blue line. Samples of posterior predictive distribution are shown as light blue bars. Note: Observed and predicted counts are based on realised number of lines inspected. This will be an underestimate of the true contamination count as not all lines are inspected.

**Figure A.3.:**
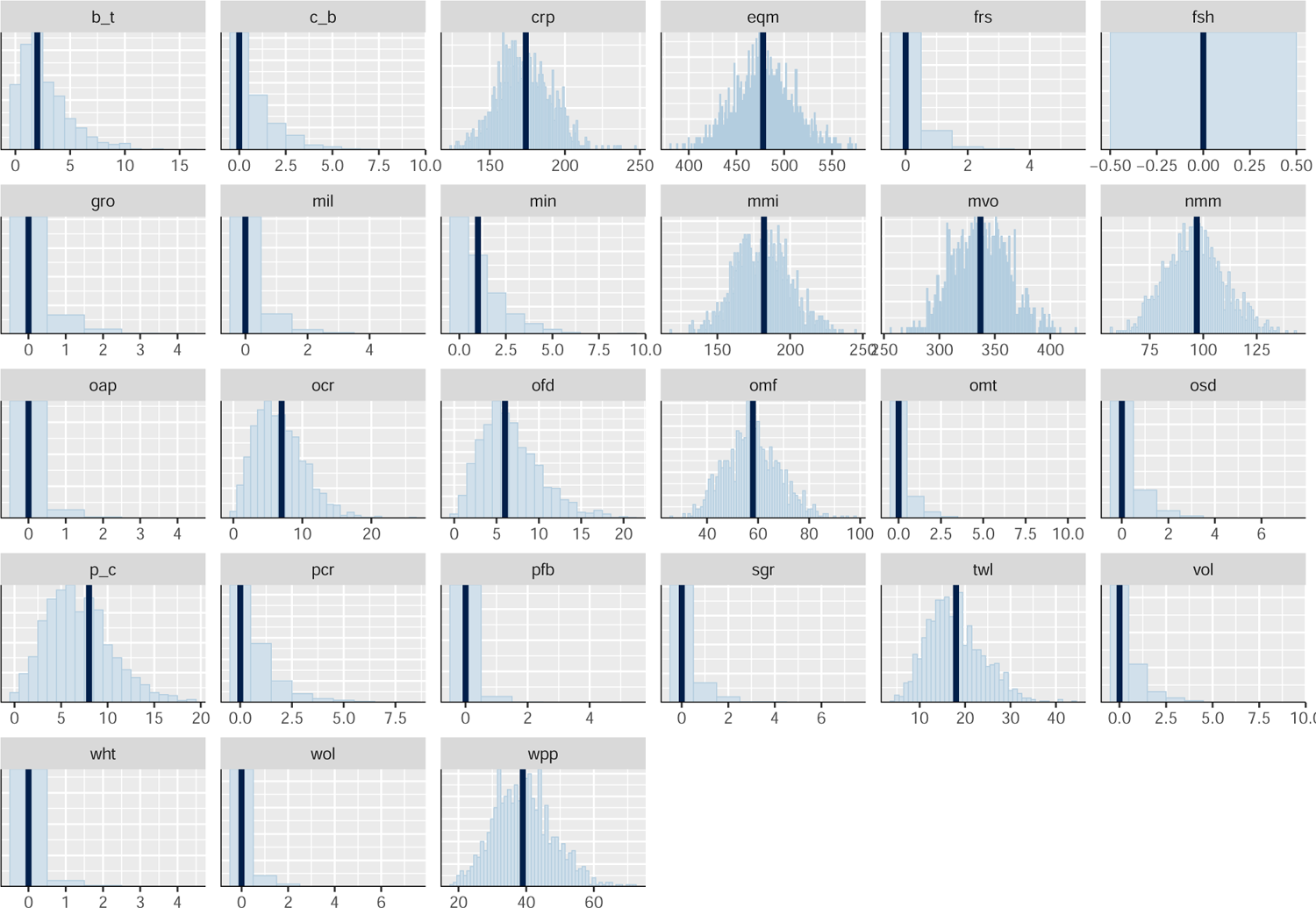
Observed vs predicted total numbers of BMSB contaminated lines across all infected exporters and years for each commodity sector. Observed counts shown as dark blue line. Samples of posterior predictive distribution are shown as light blue bars. Note: Observed and predicted counts are based on realised number of lines inspected. This will be an underestimate of the true contamination count as not all lines are inspected.

**Figure A.4.:**
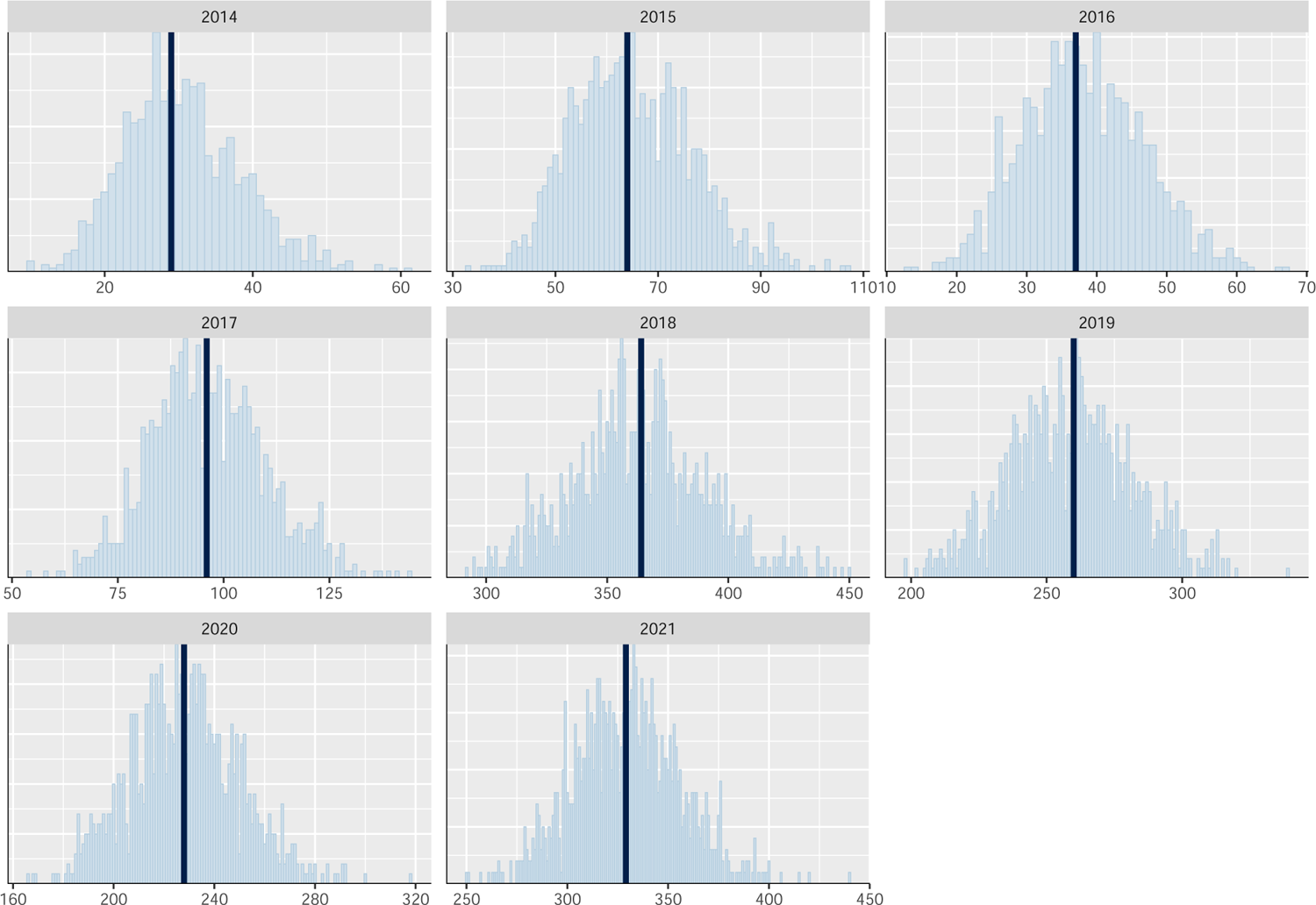
Observed vs predicted total numbers of BMSB contaminated lines across all infected exporters and commodity types for each year. Observed counts shown as dark blue line. Samples of posterior predictive distribution are shown as light blue bars. Note: Observed and predicted counts are based on realised number of lines inspected. This will be an underestimate of the true contamination count as not all lines are inspected.

## B. Posterior predictive checks for Spongy moth

**Figure B.1.:**
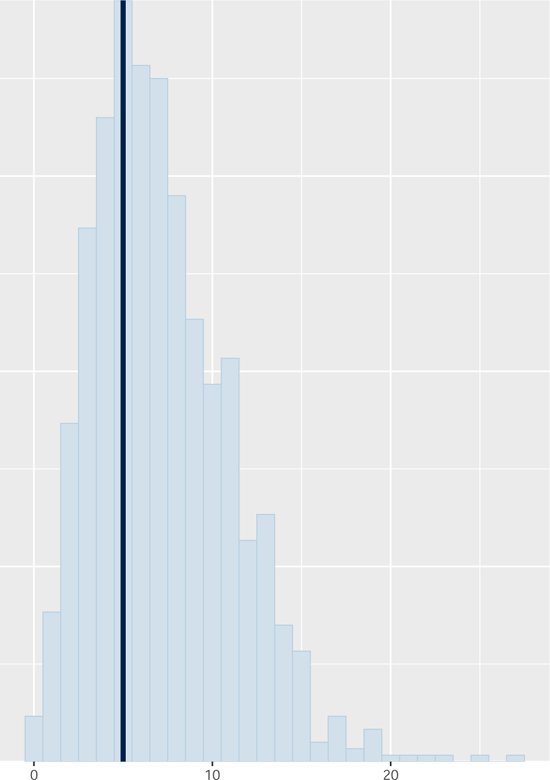
Observed vs predicted total numbers of Spongy moth contaminated lines across all infected exporters, commodities and years. Observed counts shown as dark blue line. Samples of posterior predictive distribution are shown as light blue bars. Note: Observed and predicted counts are based on realised number of lines inspected. This will be an underestimate of the true contamination count as not all lines are inspected.

**Figure B.2.:**
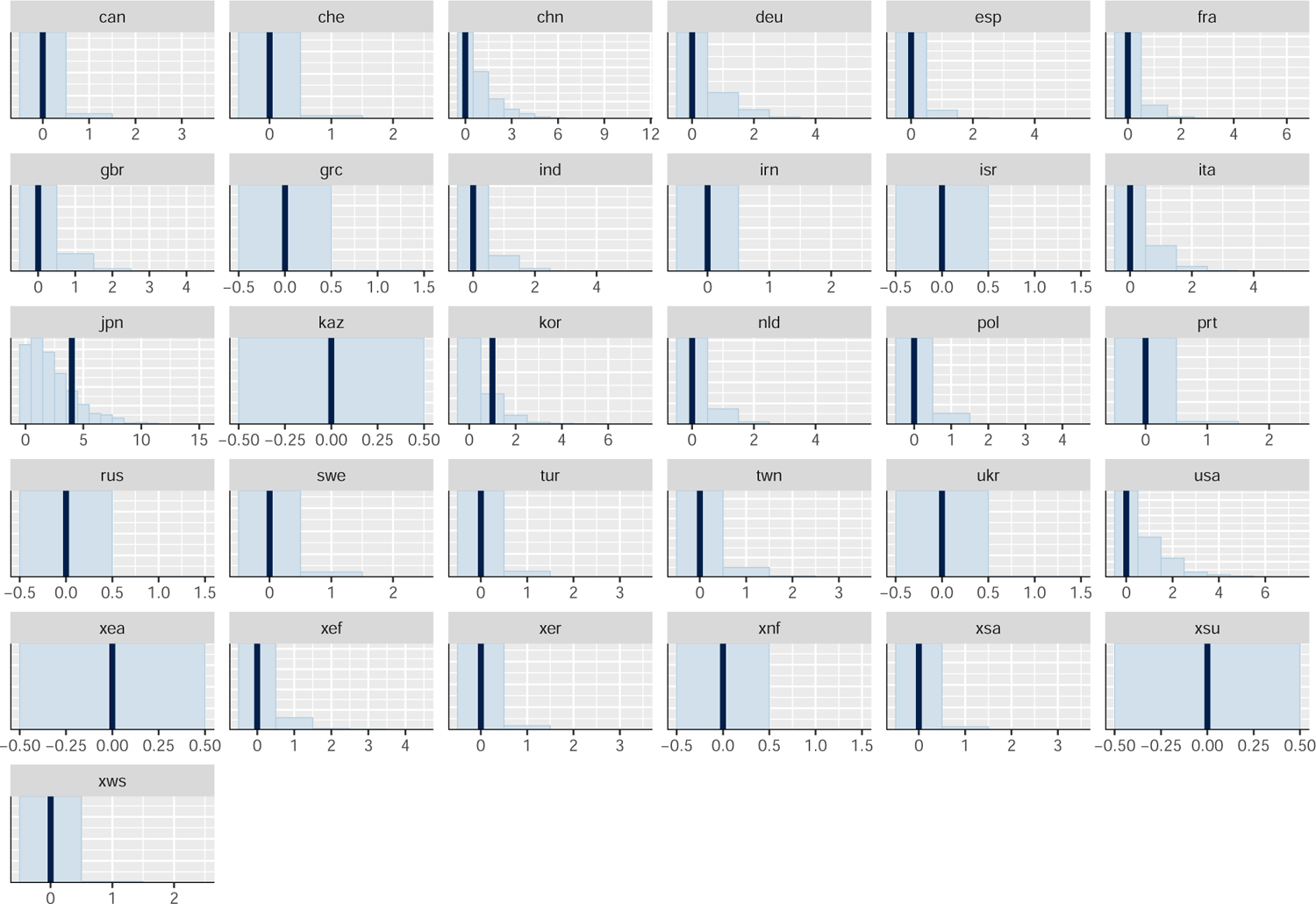
Observed vs predicted total numbers of Spongy moth contaminated lines across all years and commodity types for each infected exporter. Observed counts shown as dark blue line. Samples of posterior predictive distribution are shown as light blue bars. Note: Observed and predicted counts are based on realised number of lines inspected. This will be an underestimate of the true contamination count as not all lines are inspected.

**Figure B.3.:**
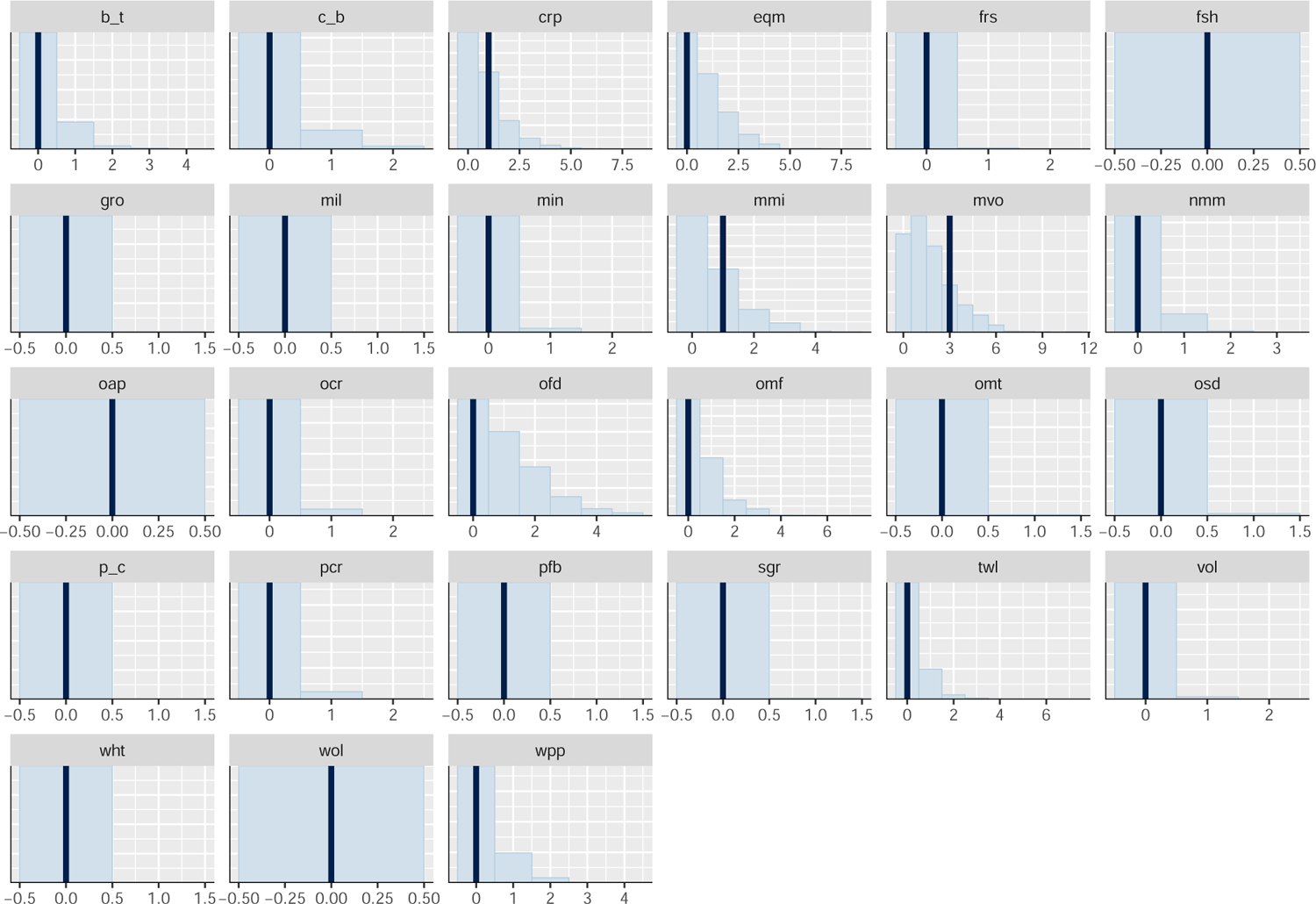
Observed vs predicted total numbers of Spongy moth contaminated lines across all infected exporters and years for each commodity sector. Observed counts shown as dark blue line. Samples of posterior predictive distribution are shown as light blue bars. Note: Observed and predicted counts are based on realised number of lines inspected. This will be an underestimate of the true contamination count as not all lines are inspected.

**Figure B.4.:**
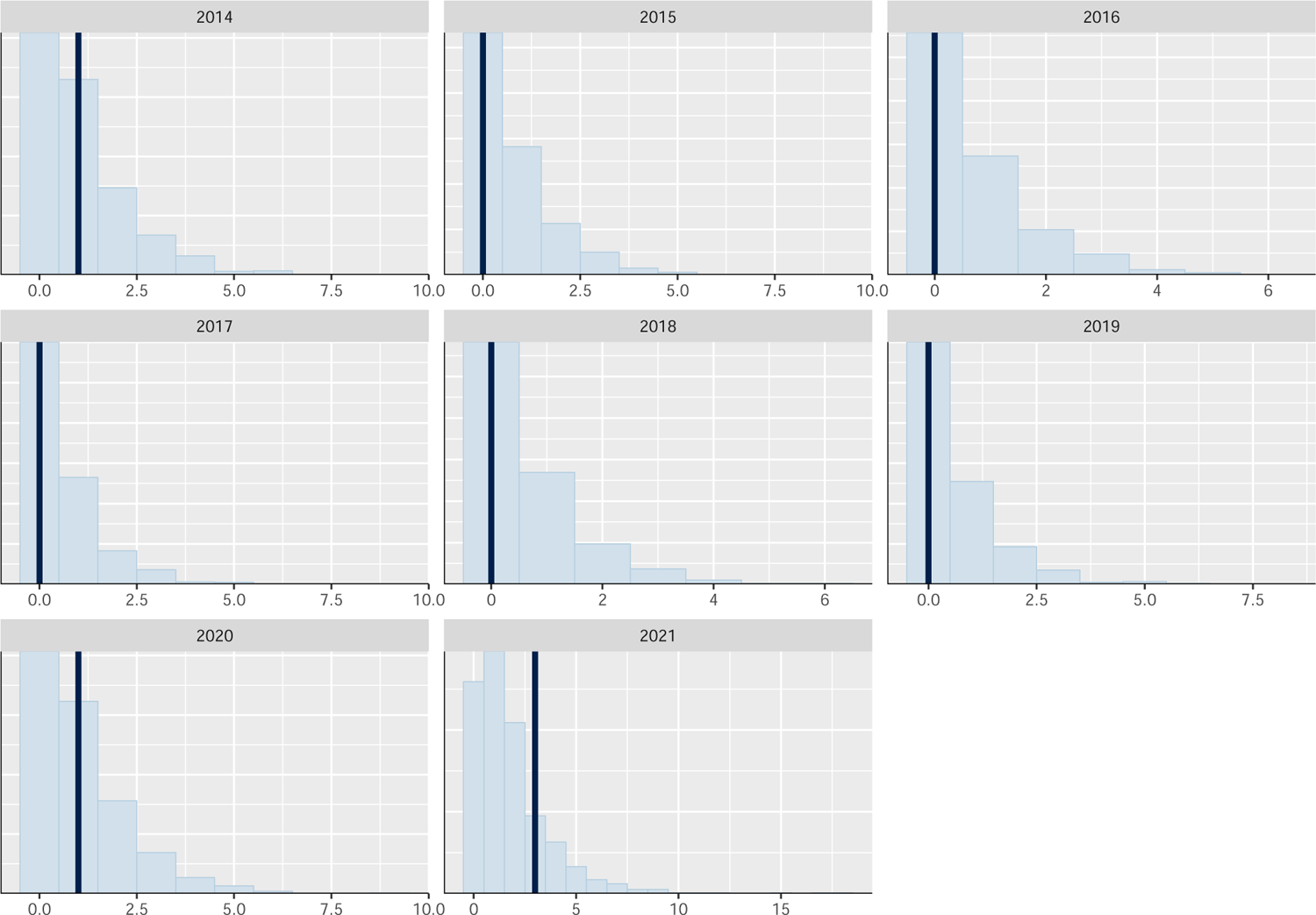
Observed vs predicted total numbers of Spongy moth contaminated lines across all infected exporters and commodity types for each year. Observed counts shown as dark blue line. Samples of posterior predictive distribution are shown as light blue bars. Note: Observed and predicted counts are based on realised number of lines inspected. This will be an underestimate of the true contamination count as not all lines are inspected.

## C. Posterior predictive checks for Asian honey bee

**Figure C.1.:**
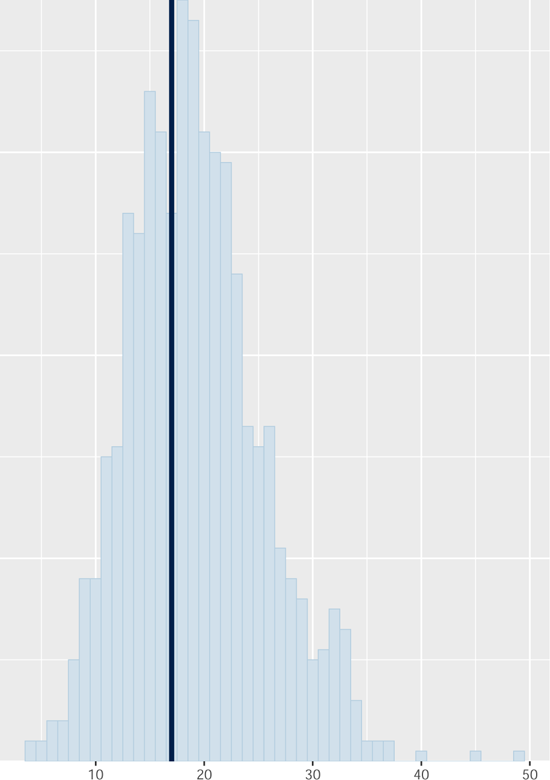
Observed vs predicted total numbers of Asian honey bee contaminated lines across all infected exporters, commodities and years. Observed counts shown as dark blue line. Samples of posterior predictive distribution are shown as light blue bars. Note: Observed and predicted counts are based on realised number of lines inspected. This will be an underestimate of the true contamination count as not all lines are inspected.

**Figure C.2.:**
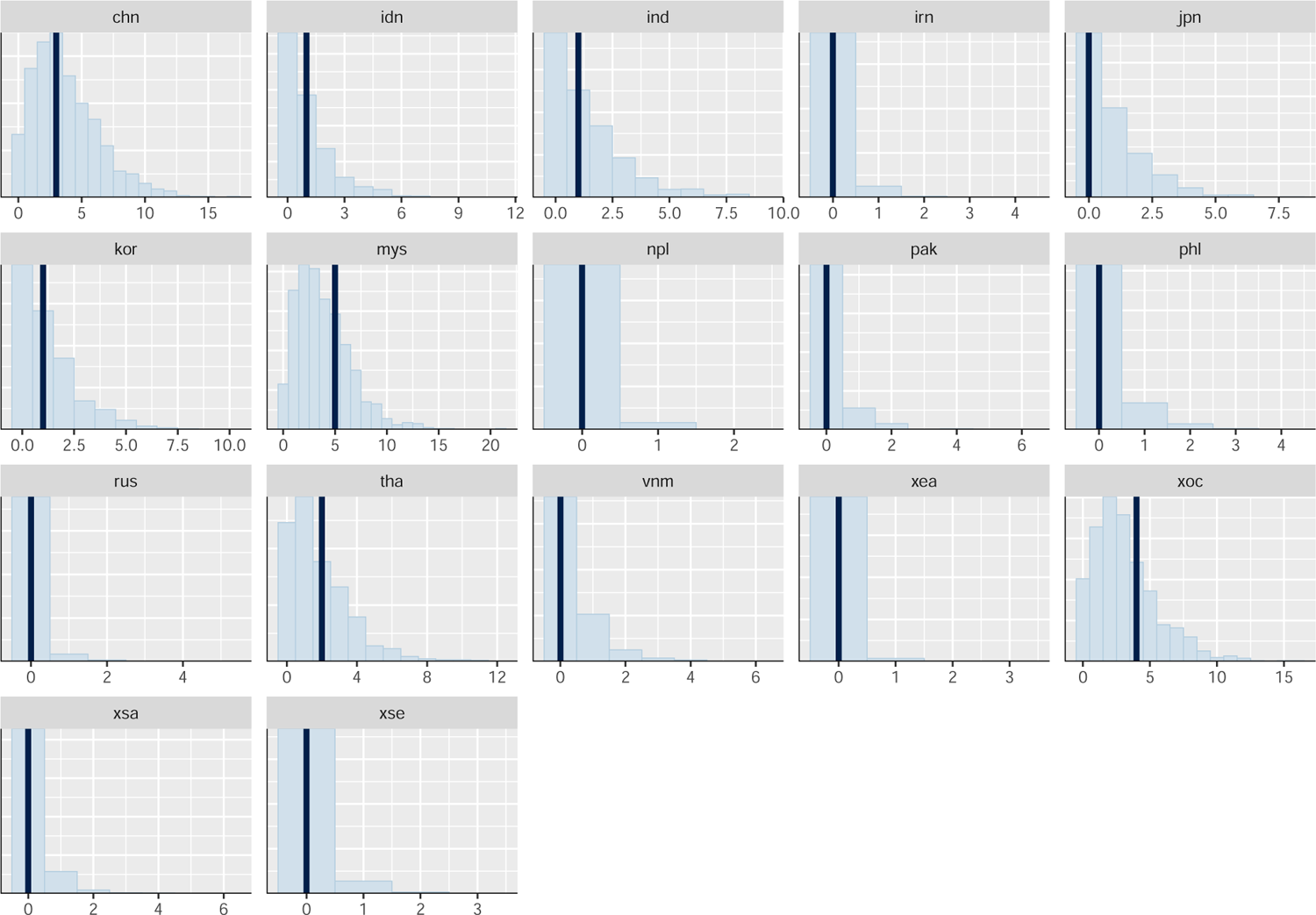
Observed vs predicted total numbers of Asian honey bee contaminated lines across all years and commodity types for each infected exporter. Observed counts shown as dark blue line. Samples of posterior predictive distribution are shown as light blue bars. Note: Observed and predicted counts are based on realised number of lines inspected. This will be an underestimate of the true contamination count as not all lines are inspected.

**Figure C.3.:**
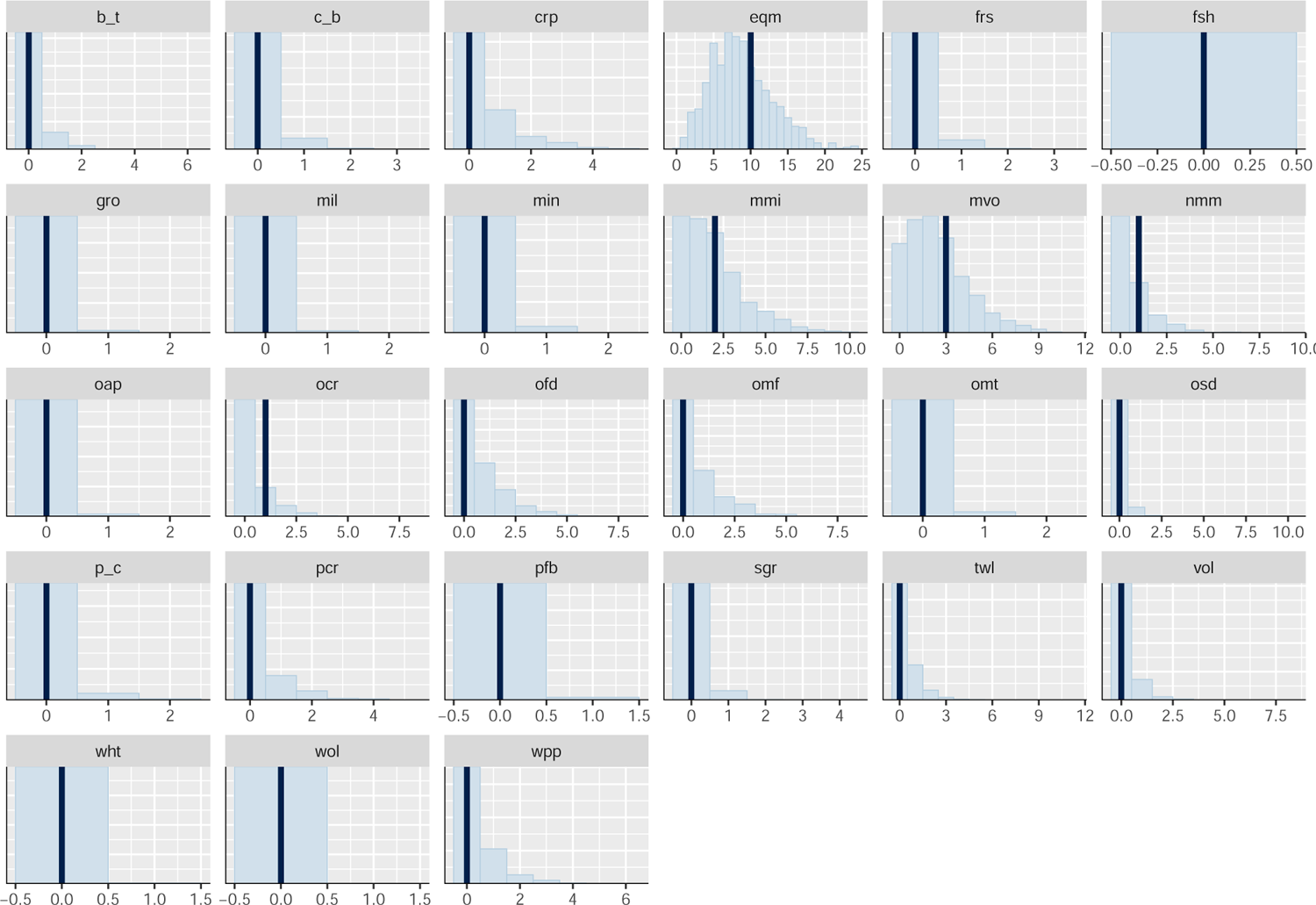
Observed vs predicted total numbers of Asian honey bee contaminated lines across all infected exporters and years for each commodity sector. Observed counts shown as dark blue line. Samples of posterior predictive distribution are shown as light blue bars. Note: Observed and predicted counts are based on realised number of lines inspected. This will be an underestimate of the true contamination count as not all lines are inspected.

**Figure C.4.:**
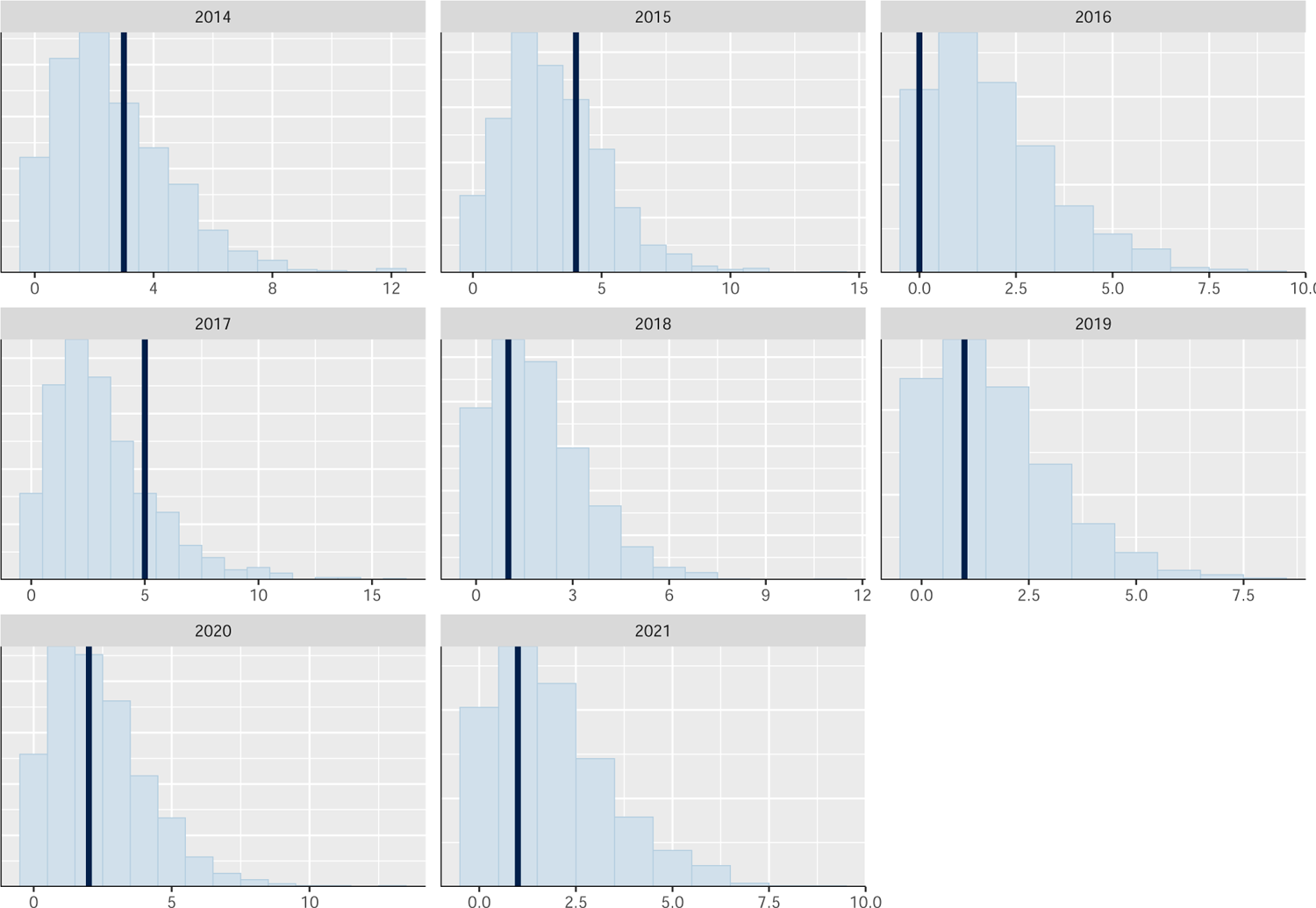
Observed vs predicted total numbers of Asian honey bee contaminated lines across all infected exporters and commodity types for each year. Observed counts shown as dark blue line. Samples of posterior predictive distribution are shown as light blue bars. Note: Observed and predicted counts are based on realised number of lines inspected. This will be an underestimate of the true contamination count as not all lines are inspected.

## D. Posterior predictive checks for Giant African snail

**Figure D.1.:**
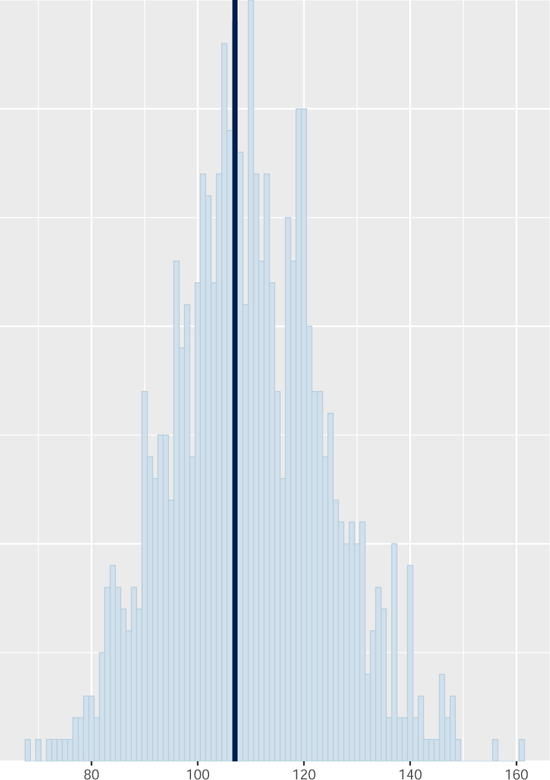
Observed vs predicted total numbers of Giant African snail contaminated lines across all infected exporters, commodities and years. Observed counts shown as dark blue line. Samples of posterior predictive distribution are shown as light blue bars. Note: Observed and predicted counts are based on realised number of lines inspected. This will be an underestimate of the true contamination count as not all lines are inspected.

**Figure D.2.:**
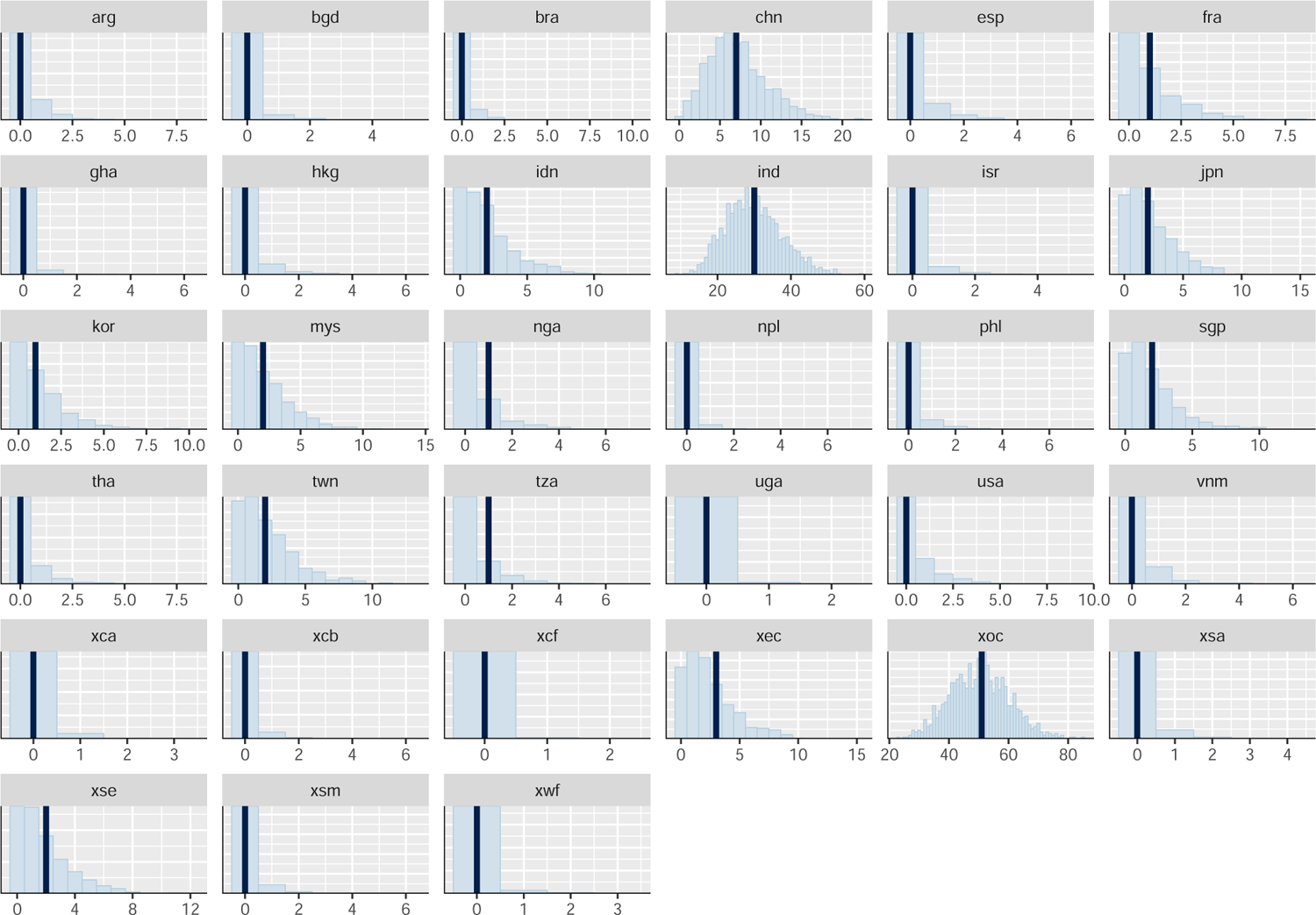
Observed vs predicted total numbers of Giant African snail contaminated lines across all years and commodity types for each infected exporter. Observed counts shown as dark blue line. Samples of posterior predictive distribution are shown as light blue bars. Note: Observed and predicted counts are based on realised number of lines inspected. This will be an underestimate of the true contamination count as not all lines are inspected.

**Figure D.3.:**
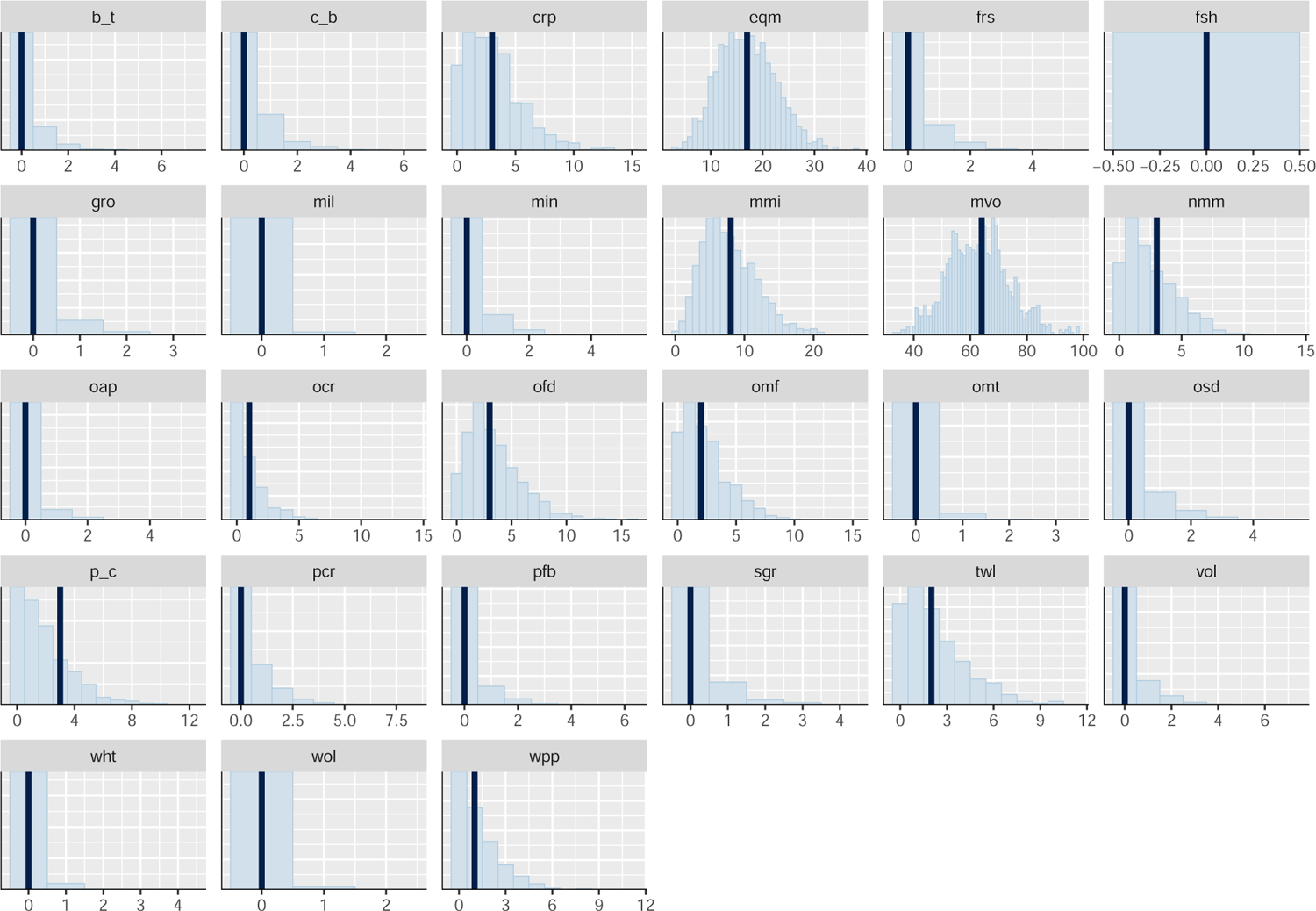
Observed vs predicted total numbers of Giant African snail contaminated lines across all infected exporters and years for each commodity sector. Observed counts shown as dark blue line. Samples of posterior predictive distribution are shown as light blue bars. Note: Observed and predicted counts are based on realised number of lines inspected. This will be an underestimate of the true contamination count as not all lines are inspected.

**Figure D.4.:**
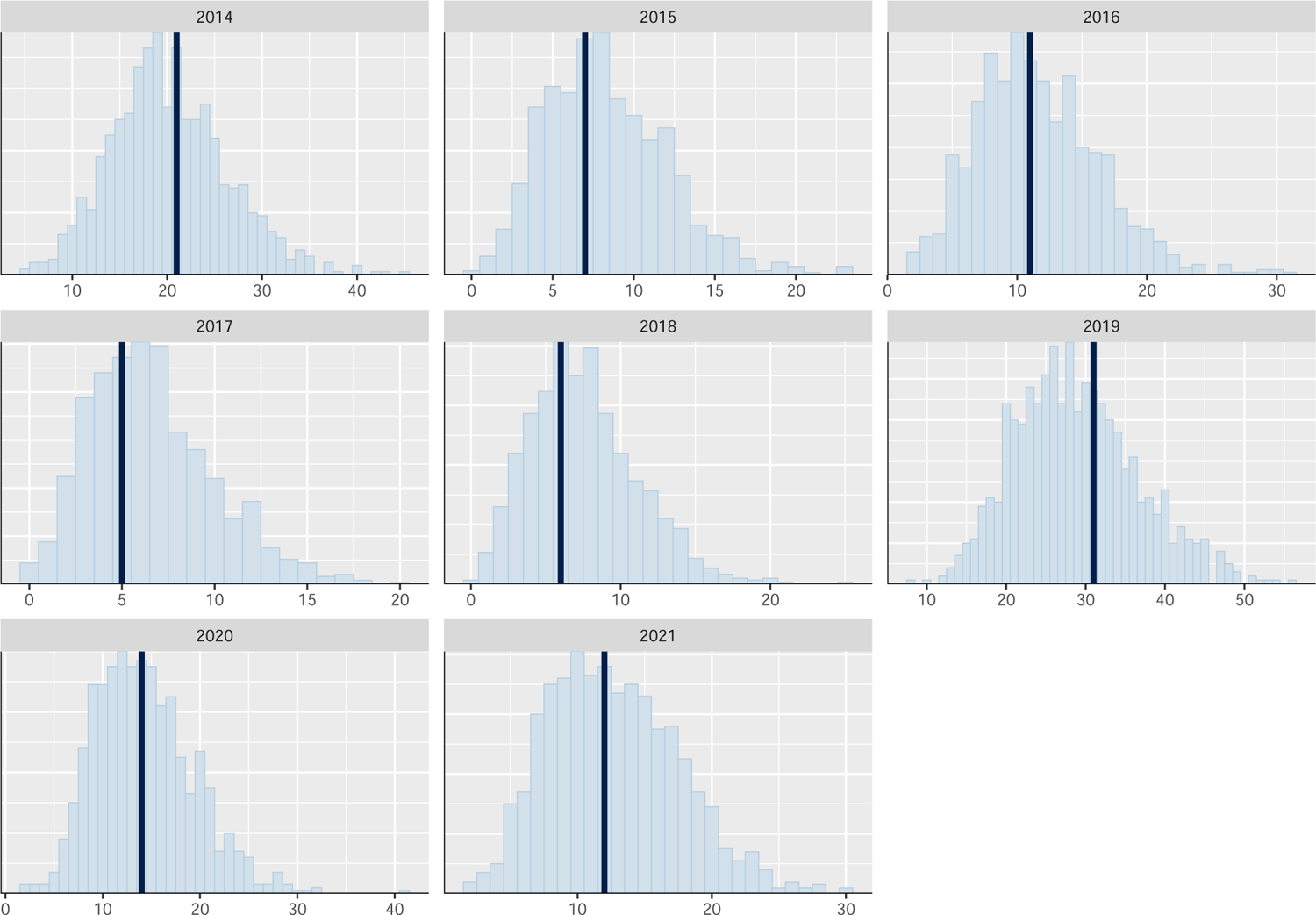
Observed vs predicted total numbers of Giant African snail contaminated lines across all infected exporters and commodity types for each year. Observed counts shown as dark blue line. Samples of posterior predictive distribution are shown as light blue bars. Note: Observed and predicted counts are based on realised number of lines inspected. This will be an underestimate of the true contamination count as not all lines are inspected.

## E. Posterior predictive checks for Khapra beetle

**Figure E.1.:**
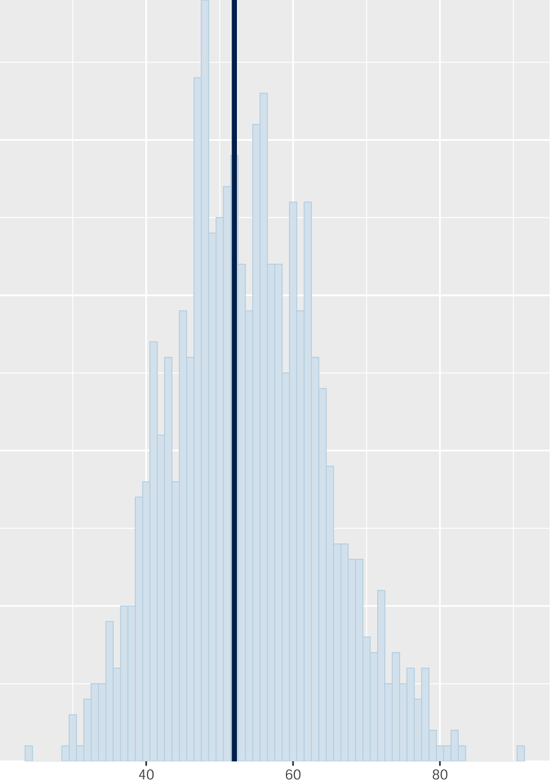
Observed vs predicted total numbers of Khapra beetle contaminated lines across all infected exporters, commodities and years. Observed counts shown as dark blue line. Samples of posterior predictive distribution are shown as light blue bars. Note: Observed and predicted counts are based on realised number of lines inspected. This will be an underestimate of the true contamination count as not all lines are inspected.

**Figure E.2.:**
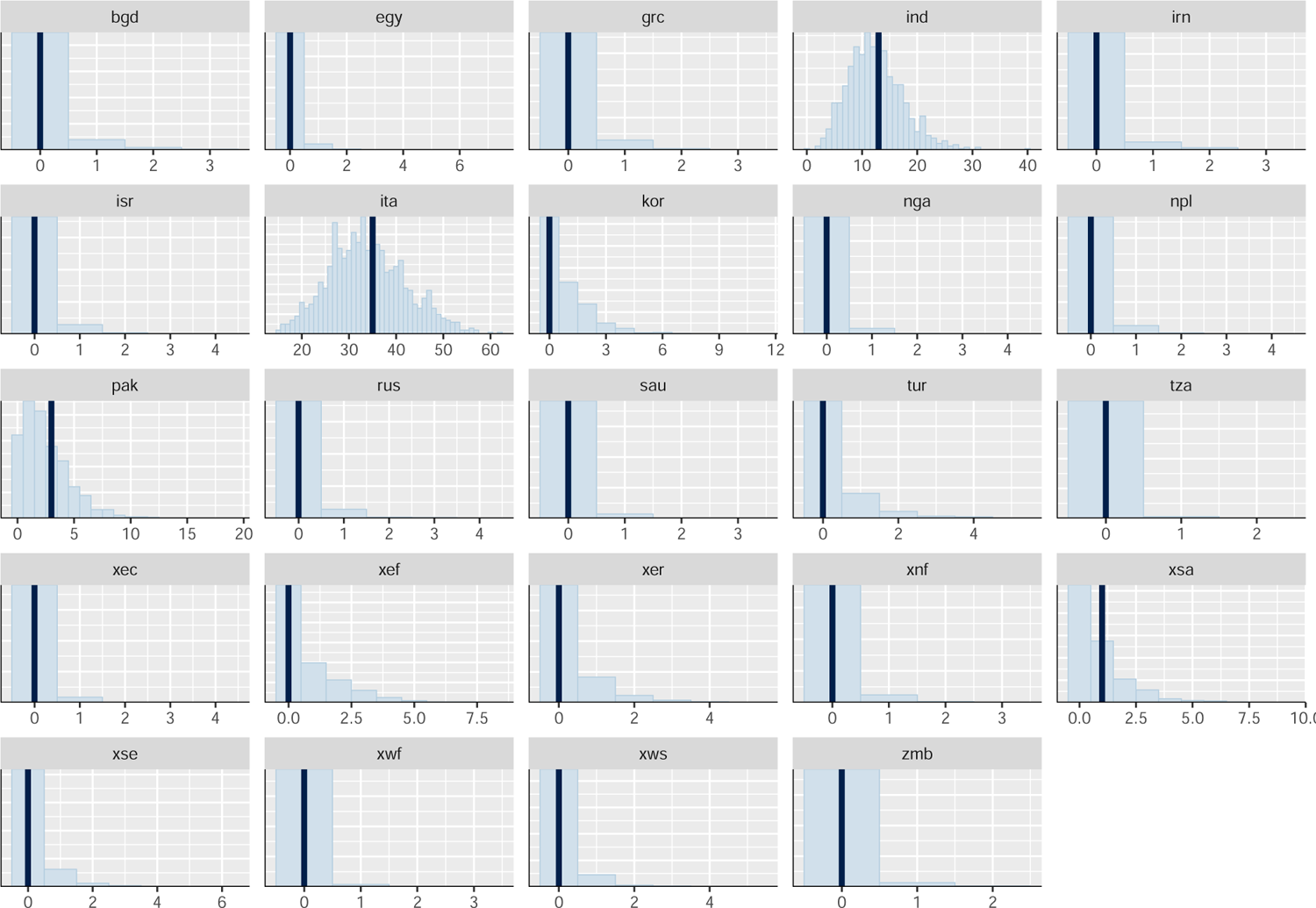
Observed vs predicted total numbers of Khapra beetle contaminated lines across all years and commodity types for each infected exporter. Observed counts shown as dark blue line. Samples of posterior predictive distribution are shown as light blue bars. Note: Observed and predicted counts are based on realised number of lines inspected. This will be an underestimate of the true contamination count as not all lines are inspected.

**Figure E.3.:**
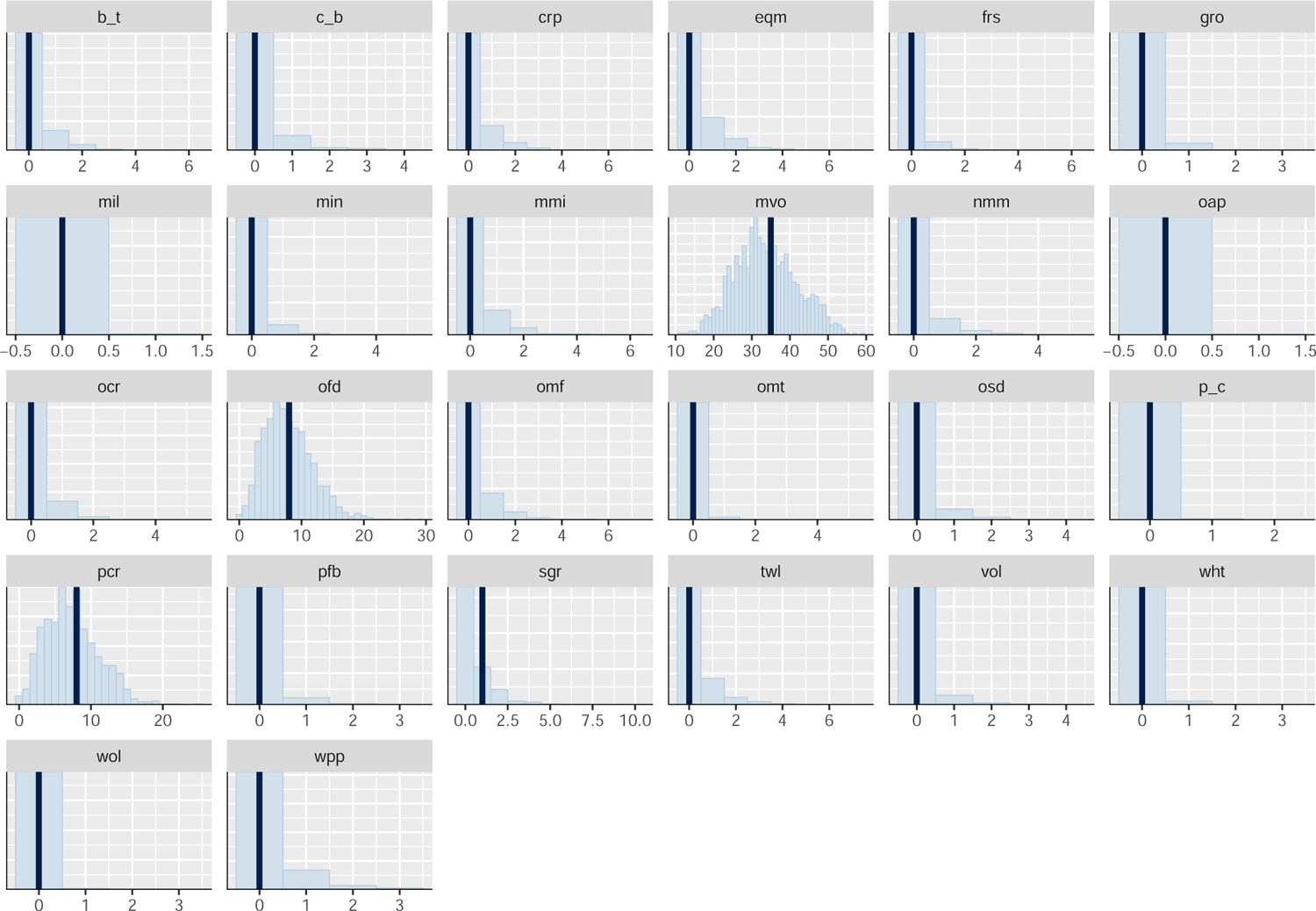
Observed vs predicted total numbers of Khapra beetle contaminated lines across all infected exporters and years for each commodity sector. Observed counts shown as dark blue line. Samples of posterior predictive distribution are shown as light blue bars. Note: Observed and predicted counts are based on realised number of lines inspected. This will be an underestimate of the true contamination count as not all lines are inspected.

**Figure E.4.:**
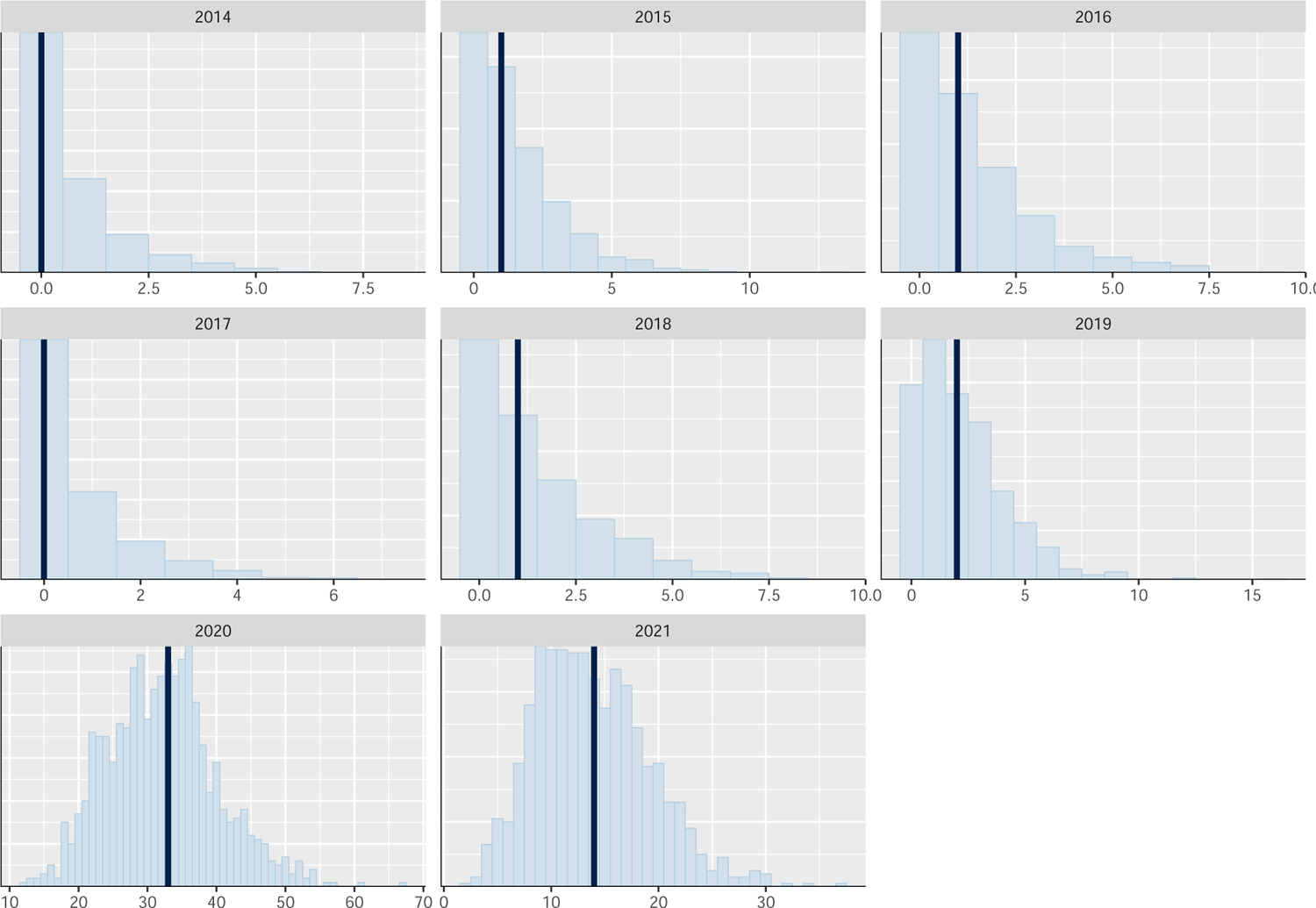
Observed vs predicted total numbers of Khapra beetle contaminated lines across all infected exporters and commodity types for each year. Observed counts shown as dark blue line. Samples of posterior predictive distribution are shown as light blue bars. Note: Observed and predicted counts are based on realised number of lines inspected. This will be an underestimate of the true contamination count as not all lines are inspected.

## Footnotes

1 (https://apps.cebra.unimelb.edu.au/trade-dashboard/)

2 Study codes: *bgd*, *ind*, *npl*, *pak*, and *xsa* within Table 4.1.

3 Study codes: *idn*, *mys*, *phl*, *sgp*, *tha*, *vnm*, and *xse* within Table 4.1.

4 Note: zip file is approximately 900mb: (https://mediaflux.researchsoftware.unimelb.edu.au:443/mflux/share.mfjp?_token=seBBIbIl4Wf1u3yvyg1211282383971&browser=true&filename=interactive_maps.zip)

5 Note: As Khapra beetle is a threat of stored food items, and thus, not exposed to ambient climate conditions, we did not estimate climate-induced impacts on contamination rates or establishment exposure for this species.

6 Readily available via the online platform: Biosecurity Commons (https://www.biosecuritycommons.org.au)

1 See Cantele, et al. (2021) for a comprehensive introduction to CGE models.

1 The heterogeneity of damages across countries from global warming is nicely illustrated in Piontek *et al*. (2021), see figure 4 and supporting text in the article, which is developed from Kompas *et al*. (2018).

2 (http://live.magicc.org/)

3 (http://www.pik-potsdam.de/$\sim$mmalte/rcps/)

1 An expanded shock regime encompassing the impacts of bush fires, flooding, and sea level rise and storm surge on coastal capital and environmental infrastructure will result in far larger damages.

1 Note: Identification of country of origin can be challenging, especially for threats such as Khapra beetle, which allow it to persist in subfloors of containers for years. As such, in some cases, country of origin is a ‘best approximation’ and as a consequence, results derived from its use should be interpreted with caution.

1 excluding: cns, fin, osg, rmk, svs, trd, tsp, utl. pdr was also excluded due to neglible imports of Paddy rice into Australia.

3 (https://www.gtap.agecon.purdue.edu/resources/download/11724.xlsx)

4 (gbif.org)

5 Often records with no coordinates are entered into databases as 0,0

6 GCMs included in model ensemble were: ACCESS-CM2, ACCESS-ESM1-5, CanESM5, CanESM5- CanOE, CMCC-ESM2, CNRM-CM6-1, CNRM-CM6-1-HR, CNRM-ESM2-1, EC-Earth3-Veg, EC-Earth3- Veg-LR, FIO-ESM-2-0, GISS-E2-1-G, GISS-E2-1-H, HadGEM3-GC31-LL, INM-CM4-8, INM-CM5-0, IPSL-CM6A-LR, MIROC-ES2L, MIROC6, MPI-ESM1-2-HR, MPI-ESM1-2-LR, MRI-ESM2-0, UKESM1- 0-LL.

7 Note: we also extracted ensembled lower and upper 10th and 90th percentiles as well as standard deviations. However, we opted to use the mean estimates as there was little variation in climate suitability predictions across GCMs)

8 For those interested in undertaking expert elicitation to derive such parameters, CEBRA in collaboration with the Australian National University have recently developed an easy-to-use web platform for undertaking the IDEA protocol (Courtney Jones *et al*., 2023). This platform can be accessed at: https://www.ideacology.com

9 Note: zip file is approximately 900mb (https://mediaflux.researchsoftware.unimelb.edu.au:443/mflux/share.mfjp?_token=seBBIbIl4Wf1u3yvyg1211282383971&browser=true&filename=interactive_maps.zip)

1 While these estimates may appear high, it is important to acknowledge that they are the expectations across all imported lines into Australia, not just those inspected. Most imported lines exhibit relatively low rates of inspection (Median proportion of lines inspected = 6.4%). Moreover, these estimates do not account for the viability of a contamination, which in most instances is likely to be low (i.e. low survivorship or low contaminated population sizes.)

1 It is a pest of stored foodstuff, and thus, rarely subject to ambient climate conditions

## Bibliography

Aguiar A, Narayanan B, McDougall R (2016) An Overview of the GTAP 9 Data Base. Journal of Global Economic Analysis, 1, 181–208. doi:10.21642/JGEA.010103AF. URL https://jgea.org/ojs/index.php/jgea/article/view/23.

Araújo MB, Rozenfeld A (2014) The geographic scaling of biotic interactions. Ecography, 37, 406–415.

Babiker M, Gurgel A, Paltsev S, Reilly J (2009) Forward-looking versus recursive- dynamic modeling in climate policy analysis: A comparison. Economic Modelling, 26, 1341–1354. doi:10.1016/j.econmod.2009.06.009. URL https://linkinghub.elsevier.com/retrieve/pii/S0264999309001035.

Barry S, Elith J, Heersink D, Caley P, Kearney M, Tenant P, Arthur A (2015) Final report: CEBRA 1402B Tools and approaches for invasive species distribution modelling for surveillance. Tech. rep., CSIRO.

Breiner FT, Nobis MP, Bergamini A, Guisan A (2017) Optimizing ensembles of small models for predicting the distribution of species with few occurrences. Methods in Ecology and Evolution, 9, 802–808.

Brooks SP, Gelman A (1998) General methods for monitoring convergence of iterative simulations. Journal of Computational and Graphical Statistics, 7, 434–455.

Burges HD (1963) Studies on the Dermestid beetle Trogoderma granarium Everts. VI.—Factors inducing diapause. Bulletin of Entomological Research, 54, 571–587.

Bürkner PC (2021) Bayesian item response modeling in R with brms and Stan. Journal of Statistical Software, 100, 1–54. doi:10.18637/jss.v100.i05.

Camac JS, Baumgartner JB, Garms B, Robinson A, T K (2021a) Estimating trading partner exposure risk to new pests or diseases. Tech. Rep. CEBRA project: 190606, Centre of Excellence for Biosecurity Risk Analysis.

Camac JS, Baumgartner JB, Hester S, Subasinghe R, Collins S (2021b) Using edmaps and Zonation to inform multi-pest early-detection surveillance designs. Tech. Rep. CEBRA project: 20121001, Centre of Excellence for Biosecurity Risk Analysis.

Camac JS, Robinson A, Elith J (2020) Developing pragmatic maps of establishment likelihood for plant pests. Tech. Rep. CEBRA project: 170607, Centre of Excellence for Biosecurity Risk Analysis.

Cantele M (2023) A glimpse into land-use futures: Simulating socioeconomic pathways within a temporally recursive, land-use explicit computable general equilibrium model. Ph.D. thesis, University of Melbourne, Parksville, Australia. URL https://minerva-access.unimelb.edu.au/items/2a002d14-e52e-4b54-b674-5d5b5de18308.

Cantele M, Bal P, Kompas T, Hadjikakou M, Wintle B (2021) Equilibrium Modeling for Environmental Science: Exploring the Nexus of Economic Systems and Environmental Change. Earth’s Future, 9. doi:10.1029/2020EF001923. URL https://onlinelibrary.wiley.com/doi/10.1029/2020EF001923.

Commission E, Centre JR, Lutz W, Stilianakis N, Stonawski M, Goujon A, Samir K (2018) Demographic and human capital scenarios for the 21st century – 2018 assessment for 201 countries. Publications Office. 10.2760/835878.

Courtney Jones SK, Geange SR, Hanea A, et al. (2023) Ideacology: An interface to streamline and facilitate efficient, rigorous expert elicitation in ecology. Methods in Ecology and Evolution, 14, 2019–2028. 10.1111/2041-210X.14017. URL https://besjournals.onlinelibrary.wiley.com/doi/abs/10.1111/2041-210X.14017.

DAFF (2020) Climatch v2.0. URL https://climatch.cp1.agriculture.gov.au.

Dellink R, Chateau J, Lanzi E, Magné B (2017) Long-term economic growth projections in the Shared Socioeconomic Pathways. Global Environmental Change, 42, 200–214. doi:10.1016/j.gloenvcha.2015.06.004. URL https://linkinghub.elsevier.com/retrieve/pii/S0959378015000837.

Dodd A, Baumgartner J, Kompas T (2021) Final report: Stakeholder perceptions of and options for refining the cebra ‘value’ model for use within dawe. Tech. Rep. CEBRA project 20100401, Centre of Excellence for Biosecurity Risk Analysis.

Dodd AJ, McCarthy MA, Ainsworth N, Burgman MA (2015) Identifying hotspots of alien plant naturalisation in Australia: approaches and predictions. Biological Invasions, 18, 631 – 645. doi:10.1007/s10530-015-1035-8.

Drake JM (2015) Range bagging: a new method for ecological niche modelling from presence-only data. Journal of The Royal Society Interface, 12, 20150086–9.

Elith J (2017) Predicting distributions of invasive species. In: Invasive species risk assessment and management (eds. Robinson AP, Walshe T, Burgman MA, Nunn M), pp. 93–129. Cambridge University Press.

Fick SE, Hijmans RJ (2017) WorldClim 2: new 1-km spatial resolution climate surfaces for global land areas. International Journal of Climatology, 37, 4302–4315.

Garrard GE, Bekessy SA, McCarthy MA, Wintle BA (2008) When have we looked hard enough? A novel method for setting minimum survey effort protocols for flora surveys. Austral Ecology, 33, 986–998.

Garrard GE, McCarthy MA, Williams NSG, Bekessy SA, Wintle BA (2012) A general model of detectability using species traits. Methods in Ecology and Evolution, 4, 45–52.

Gelman A, Hill J (2007) Data analysis using regression and multilevel/hierarchical models. Cambridge University Press, Cambridge.

Glotter MJ, Pierrehumbert RT, Elliott JW, Matteson NJ, Moyer EJ (2014) A simple carbon cycle representation for economic and policy analyses. Climatic Change, 126, 319–335. doi:10.1007/s10584-014-1224-y.

Ha PV, Kompas T, Nguyen HTM, Long CH (2017) Building a better trade model to determine local effects: A regional and intertemporal GTAP model. Economic Modelling, 67, 102–113. doi:10.1016/j.econmod.2016.10.015.

Hemming V, Burgman MA, Hanea AM, McBride MF, Wintle BC (2018) A practical guide to structured expert elicitation using the idea protocol. Methods in Ecology and Evolution, 9, 169–180. 10.1111/2041-210X.12857. URL https://besjournals.onlinelibrary.wiley.com/doi/abs/10.1111/2041-210X.12857.

Hertel TW, ed. (1997) Global Trade Analysis: Modelling and Applications. Cambridge University Press, Cambridge, New York.

Higgins SI, Richardson DM (2014) Invasive plants have broader physiological niches. Proceedings of the National Academy of Sciences, 111, 10610–10614.

Hill M, Caley P, Camac J, Elith J, Barry S (2022) A novel method accounting for predictor uncertainty and model transferability of invasive species distribution models. *bioRxiv*. doi:10.1101/2022.03.14.483865. URL https://www.biorxiv.org/content/early/2022/03/17/2022.03.14.483865.

Hulme PE (2009) Trade, transport and trouble: managing invasive species pathways in an era of globalization. Journal of Applied Ecology, 46, 10–18. 10.1111/j.1365-2664.2008.01600.x. URL https://besjournals.onlinelibrary.wiley.com/doi/abs/10.1111/j.1365-2664.2008.01600.x.

Kc S (2020) Updated demographic SSP4 and SSP5 scenarios complementing the SSP1-3 scenarios published in 2018. Working Paper WP-20-016, International Institute Of Applied System Analysis, Laxenburg, Austria.

Kearney MR, Porter WP (2020) Nichemapr – an r package for biophysical modelling: the ectotherm and dynamic energy budget models. Ecography, 43, 85–96. doi: 10.1111/ecog.04680. URL https://onlinelibrary.wiley.com/doi/abs/10.1111/ecog.04680.

Kompas T, Ha PV (2021) Climate change impacts on regional Australia. Centre for Environmental and Economic Research, University of Melbourne.

Kompas T, Ha PV, Nhu CT (2018) The effects of climate change on GDP by country and the global economic gains from complying with the paris climate accord. Earth’s Future, 6, 1153–1173. doi:10.1029/2018EF000922.

Kompas T, Van Ha P (2019) The ‘curse of dimensionality’ resolved: The effects of climate change and trade barriers in large dimensional modelling. Economic Modelling, 80, 103–110. doi:10.1016/j.econmod.2018.08.011. URL https://www.sciencedirect.com/science/article/pii/S0264999317310386.

Kriticos D, Maywald GF, Yonow T, Zurcher EJ, Herrmann NI, Sutherst RW (2015) Climex Version 4: Exploring the effects of climate on plants, animals and diseases. CSIRO.

Kriticos DJ, Kean JM, Phillips CB, Senay SD, Acosta H, Haye T (2017) The potential global distribution of the brown marmorated stink bug, Halyomorpha halys, a critical threat to plant biosecurity. Journal of Pest Science, 90, 1–11.

Li C, Kompas T (2022) The impact of heat stress on agricultural productivity. Centre for Environmental and Economic Research, University of Melbourne.

Martin T (2017) Surveillance for detection of pests and diseases: how sure can we be of their absence? In: Invasive species risk assessment and management (eds. Robinson AP, Walshe T, Burgman MA, Nunn M), pp. 348–384. Cambridge.

Martin TG, Murphy H, Liedloff A, et al. (2015) Buffel grass and climate change: a framework for projecting invasive species distributions when data are scarce. Biological Invasions, 17, 3197–3210.

Meinshausen M, Raper S, Wigley TML (2011a) Emulating coupled atmosphere-ocean and carbon cycle models with a simpler model, MAGICC6: Part I – Model Description and Calibration. Atmospheric Chemistry and Physics, 11, 1417–1456. doi: doi:10.5194/acp-11-1417-2011.

Meinshausen M, Smith S, Calvin K, et al. (2011b) The RCP greenhouse gas concentrations and their extensions from 1765 to 2300. Climatic Change, 109, 213.

Meinshausen M, Wigley T, Raper S (2011c) Emulating atmosphere-ocean and carbon cycle models with a simpler model, MAGICC6–Part 2: Applications. Atmospheric Chemistry and Physics, 11, 1457–1471.

Moss RH, Edmonds JA, Hibbard KA, et al. (2010) The next generation of scenarios for climate change research and assessment. Nature, 463, 747–756.

Nauels A, Meinshausen M, Mengel M, Lorbacher K, Wigley TML (2017) Synthesizing long-term sea level rise projections – the MAGICC sea level model v2.0. Geoscientific Model Development, 10, 2495–2524. doi:10.5194/gmd-10-2495-2017.

Nazarenko L, Schmidt GA, Miller RL, et al. (2015) Future climate change under RCP emission scenarios with GISS ModelE2. Journal of Advances in Modeling Earth Systems, 7, 244–267. doi:10.1002/2014MS000403.

Nicholls ZRJ, Meinshausen M, Lewis J, et al. (2020) Reduced complexity model intercomparison project phase 1: Protocol, results and initial observations. Preprint, Climate and Earth System Modeling. doi:10.5194/gmd-2019-375.

Olén NB, Lehsten V (2022) High-resolution global population projections dataset developed with cmip6 rcp and ssp scenarios for year 2010–2100. Data in Brief, 40, 107804. 10.1016/j.dib.2022.107804. URL https://www.sciencedirect.com/science/article/pii/S2352340922000166.

Palma E, Vesk PA, White M, Baumgartner JB, Catford JA (2021) Plant functional traits reflect different dimensions of species invasiveness. Ecology, 102, e03317. 10.1002/ecy.3317. URL https://esajournals.onlinelibrary.wiley.com/doi/abs/10.1002/ecy.3317.

Phillips SJ, Anderson RP, Schapire RE (2006) Maximum entropy modeling of species geographic distributions. Ecological Modelling, 190, 231–259.

Piontek F, Drouet L, Emmerling J, et al. (2021) Integrated perspective on translating biophysical to economic impacts of climate change. Nature Climate Change, 11, 563– 572.

Riahi K, Grübler A, Nakicenovic N (2007) Scenarios of long-term socio-economic and environmental development under climate stabilization. Technological Forecasting and Social Change, 74, 887–935. doi:10.1016/j.techfore.2006.05.026.

Riahi K, Rao S, Krey V, etal (2011) RCP 8.5-A scenario of comparatively high greenhouse gas emissions. Climatic Change, 1, 33–57.

Riahi K, van Vuuren DP, Kriegler E, et al. (2017) The Shared Socioeconomic Pathways and their energy, land use, and greenhouse gas emissions implications: An overview. Global Environmental Change, 42, 153–168. doi:10.1016/j.gloenvcha.2016.05.009. URL https://linkinghub.elsevier.com/retrieve/pii/ S0959378016300681.

Rice KB, Bergh CJ, Bergmann EJ, et al. (2014) Biology, Ecology, and Management of Brown Marmorated Stink Bug (Hemiptera: Pentatomidae). Journal of Integrated Pest Management, 5, 1–13.

Romanello M, Napoli Cd, Green C, et al. (2023) The 2023 report of the Lancet Countdown on health and climate change: the imperative for a health-centred response in a world facing irreversible harms. The Lancet. doi:10.1016/s0140-6736(23)01859-7.

Roson R, Sartori M (2016) Estimation of climate change damage functions for 140 regions in the GTAP 9 database. Journal of Global Economic Analysis, 1, 78–115.

Seebens H, Bacher S, Blackburn TM, et al. (2021) Projecting the continental accumulation of alien species through to 2050. Global Change Biology, 27, 970–982. 10.1111/gcb.15333. URL https://onlinelibrary.wiley.com/doi/abs/10.1111/gcb.15333.

Seebens H, Blackburn TM, Dyer EE, et al. (2017) No saturation in the accumulation of alien species worldwide. Nature Communications, 8, 1 – 9. doi:10.1038/ncomms14435.

Thomson AM, Calvin KV, Smith SJ, et al. (2011) RCP4. 5: A pathway for stabilization of radiative forcing by 2100. Climatic change, 109, 77–94.

Thuiller W, Richardson DM, Pyšek P, Midgley GF, Hughes GO, Rouget M (2005) Niche-based modelling as a tool for predicting the risk of alien plant invasions at a global scale. Global Change Biology, 11, 2234–2250.

Valentin RE, Nielsen AL, Wiman NG, Lee DH, Fonseca DM (2017) Global invasion network of the brown marmorated stink bug, Halyomorpha halys. Scientific Reports, 7, 1–12.

Wintle BA, Walshe TV, Parris KM, McCarthy MA (2012) Designing occupancy surveys and interpreting non-detection when observations are imperfect. Diversity and Distributions, 18, 417–424.

Zizka A, Silvestro D, Andermann T, et al. (2019) CoordinateCleaner: Standardized cleaning of occurrence records from biological collection databases. Methods in Ecology and Evolution, 289, 110–8.

